# Incubation of palatable food craving is associated with brain-wide neuronal activation in mice

**DOI:** 10.1101/2022.05.31.494210

**Authors:** Rajtarun Madangopal, Eric R. Szelenyi, Joseph Nguyen, Megan B. Brenner, Olivia R. Drake, Diana Pham, Aniruddha Shekara, Michelle Jin, Jia Jie Choong, Connor Heins, Lauren E. Komer, Sophia J. Weber, Bruce T. Hope, Yavin Shaham, Sam A. Golden

**Affiliations:** Intramural Research Program, National Institute on Drug Abuse, Baltimore, MD, USA; University of Washington, Department of Electrical and Computer Engineering, Seattle, Washington, USA; Department of Biological Structure, University of Washington, Seattle, WA, USA.; University of Washington, Center of Excellence in Neurobiology of Addiction, Pain, and Emotion (NAPE), Seattle, Washington, USA

**Author notes:** Correspondence to: Sam A. Golden, Ph.D., Department of Biological Structure, University of Washington, Seattle, WA 98195. These authors contributed equally (Rajtarun Madangopal, Eric R. Szelenyi). **Author Contributions:** RM, ES, YS, BTH and SAG designed the experiments; RM, MJ, CH, LEK, SJW, and SAG ran the behavioral experiments and collected data; RM, MJ, CH, LEK, SJW, MBB, ORD, AS, and DP analyzed behavioral data; RM, MBB, and SAG performed whole-brain immunohistochemistry and clearing; RM, ES, and SAG optimized imaging protocols, RM, ES, MJ, JN, JJC, and SAG established analysis pipelines; RM, ES, MBB, ORD, AS, DP and JJC analyzed data; RM, ES, YS, BTH and SAG wrote the paper. All authors reviewed and approved the final version prior to submission.

**Keywords:** addiction, incubation, whole-brain analysis, mice, Fos

## Abstract

Studies using rodent models have shown that relapse to drug or food seeking increases progressively during abstinence, a phenomenon termed ‘incubation of craving’. Mechanistic studies of incubation of craving have focused on specific neurobiological targets within pre- selected brain areas. Recent methodological advances in whole-brain immunohistochemistry, clearing, and imaging now enable unbiased brain-wide cellular resolution mapping of regions and circuits engaged during learned behaviors. However, these whole brain imaging approaches were developed for mouse brains while incubation of drug craving has primarily been studied in rats and incubation of food craving has not been demonstrated in mice. Here, we established a mouse model of incubation of palatable food craving and examined food reward seeking after 1, 15, and 60 abstinence days. We then used the neuronal activity marker Fos with intact brain mapping procedures to identify corresponding patterns of brain-wide activation. Relapse to food seeking was significantly higher after 60 abstinence days than after 1 or 15 days. Using unbiased ClearMap analysis, we identified increased activation of multiple brain regions, particularly corticostriatal structures, following 60, but not 15 abstinence days. We used orthogonal SMART2 analysis to confirm these findings within corticostriatal and thalamocortical subvolumes and applied expert-guided registration to investigate subdivision and layer-specific activation patterns. Overall, we (1) identified novel brain-wide activity patterns during incubation of food seeking using complementary analytical approaches, and (2) provide a single-cell resolution whole-brain atlas that can be used to identify functional networks and global architecture underlying incubation of food craving.

**Significance Statement:** Relapse to reward seeking progressively increases during abstinence, a phenomenon termed incubation of craving. Mechanistic studies of incubation can lead to novel relapse treatments. However, previous studies have primarily used rat models and targeted region-by-region analyses and a brain-wide functional atlas of incubation of reward seeking is lacking. We established a behavioral procedure for incubation of palatable food seeking in mice and applied whole-brain activity mapping with Fos as a neuronal activity marker to identify the functional connectome of this incubation. Like rats, mice showed incubation of food seeking during abstinence. Using two complementary activity mapping approaches, we identified a brain-wide pattern of increased neural activation that mirrored incubation of food seeking after 60, but not 15, days of abstinence.

## Introduction

Studies using rodent models have shown that non-reinforced drug or food seeking progressively increases during abstinence in the home cage (1–3). This phenomenon, termed *‘incubation of craving’*, was first identified in rats after cocaine self-administration (4–7), and has since been demonstrated for other drugs such as heroin (8), methamphetamine (9), alcohol (10), and nicotine (11), as well as for non-drug rewards such as sucrose (12, 13), standard chow pellets (14), and high-carbohydrate pellets (15). These preclinical findings mirror reports of incubation of cue-induced drug craving and physiological responses in human drug-users (16–18) and could provide avenues to identify cellular and molecular mechanisms underlying persistent relapse vulnerability to both unhealthy palatable food (3, 19) and addictive drugs (1, 2, 20–22).

Immediate early gene (IEG, eg. Fos, Arc, Zif) expression serves as a proxy for strongly activated neurons, and quantification of Fos-positive (Fos+) cells is routinely used to identify changes in neural activation patterns after exposure to different unconditioned and conditioned stimuli (23–27). Previous activity-mapping studies of incubation of drug and food craving have identified several brain regions (Supplementary Table S1) relevant to (1) relapse (increased Fos expression during relapse tests vs. home cage controls), and/or (2) incubation (higher Fos expression during late abstinence test vs. early abstinence test) (1, 3, 5, 28, 29). These studies used targeted one-by-one regional quantification of Fos+ cell counts in thin sectioned tissue samples and focused on changes in specific pre-determined brain areas. Thus, it is currently unknown whether brain-wide activity patterns, including multi-regional patterns, are altered during incubation of reward seeking during abstinence.

To address this knowledge gap, we leveraged recent developments in brain-wide activity mapping approaches, including whole mouse brain immunofluorescent staining and clearing (30–35), light sheet fluorescence microscopy (LSFM) (36, 37), and open-source analysis tools (38–41) to investigate changes in brain-wide activation patterns during incubation of palatable food seeking in mice.

We trained food-sated CD-1 male mice to self-administer palatable high-carbohydrate food pellets (42, 43) for 7 days and then tested them for relapse to food seeking after 1, 15, or 60 abstinence days. We perfused and extracted their brains 90 min after the relapse tests (or directly from homecage as a baseline activity control), labeled ‘active’ Fos+ nuclei across intact mouse brains using an optimized iDISCO+ Fos immunofluorescent staining protocol (41), and imaged Fos immunofluorescence at single cell resolution using LSFM. We used the ClearMap pipeline (39, 40) for unbiased mapping of “incubation-associated” neural activation patterns across the entire anterior-posterior axis of the mouse brain. We also updated the SMART analysis package (41) to conduct targeted analysis of neural activation patterns within LSFM coronal subvolumes and used SMART2 to cross-validate and extend our ClearMap findings within a subset of corticostriatal and thalamocortical brain regions and subdivisions.

## Materials and Methods

### Subjects

We used male (n = 60) 4-6-month-old sexually experienced CD-1 mice (Charles River Lab, CRL), weighing ∼40 g prior to food self-administration training. We confirmed with CRL animal- facility staff that all sexually experienced CD-1 males had equal access to receptive females. Per CRL’s procedure male mice are pair-housed with several females from PD28 until purchase.

Pregnant females are switched with new non-pregnant females, with no break between cycles and male mice that do not successfully breed are removed from the breeding pool and not made available for purchase. We excluded 14 mice due to failure to acquire food self-administration.

The mice had free access to food and water in the homecage and were maintained on a reverse 12:12 h light-dark cycle (light off at 8 am). We only used male mice because the behavioral study was performed before the implementation of the NIH Sex as a Biological Variable Guideline.

### Apparatus

We trained and tested all mice in standard Med Associates operant chambers, enclosed in ventilated sound-attenuating cubicles. Each chamber was equipped with a stainless-steel grid floor and two side-walls, each with three modular operant panels. We used a houselight located on one side of the chamber to illuminate the chamber during the training and test sessions. Two levers served as operant manipulanda – (1) a non-retractable lever on the same side as the houselight served as the inactive lever and (2) a retractable lever on the side opposite the houselight served as the active lever; both levers were positioned 2.4 cm above the grid floor. We placed a yellow LED light for the food-paired conditioned stimulus (CS) above the active lever and equipped the central panel on the same side with a pellet receptacle connected to a pellet dispenser. Presses on the active lever (only extended during food self-administration sessions or food-seeking tests) resulted in delivery of 20-mg food pellets and a 2-s light CS (bright yellow LED), while presses on the inactive lever had no programmed consequences.

### Behavioral procedures

The experimental timeline is shown in Figure 1A. Details of the food self-administration procedure, abstinence phase, and relapse test are provided below.

**Figure 1.**
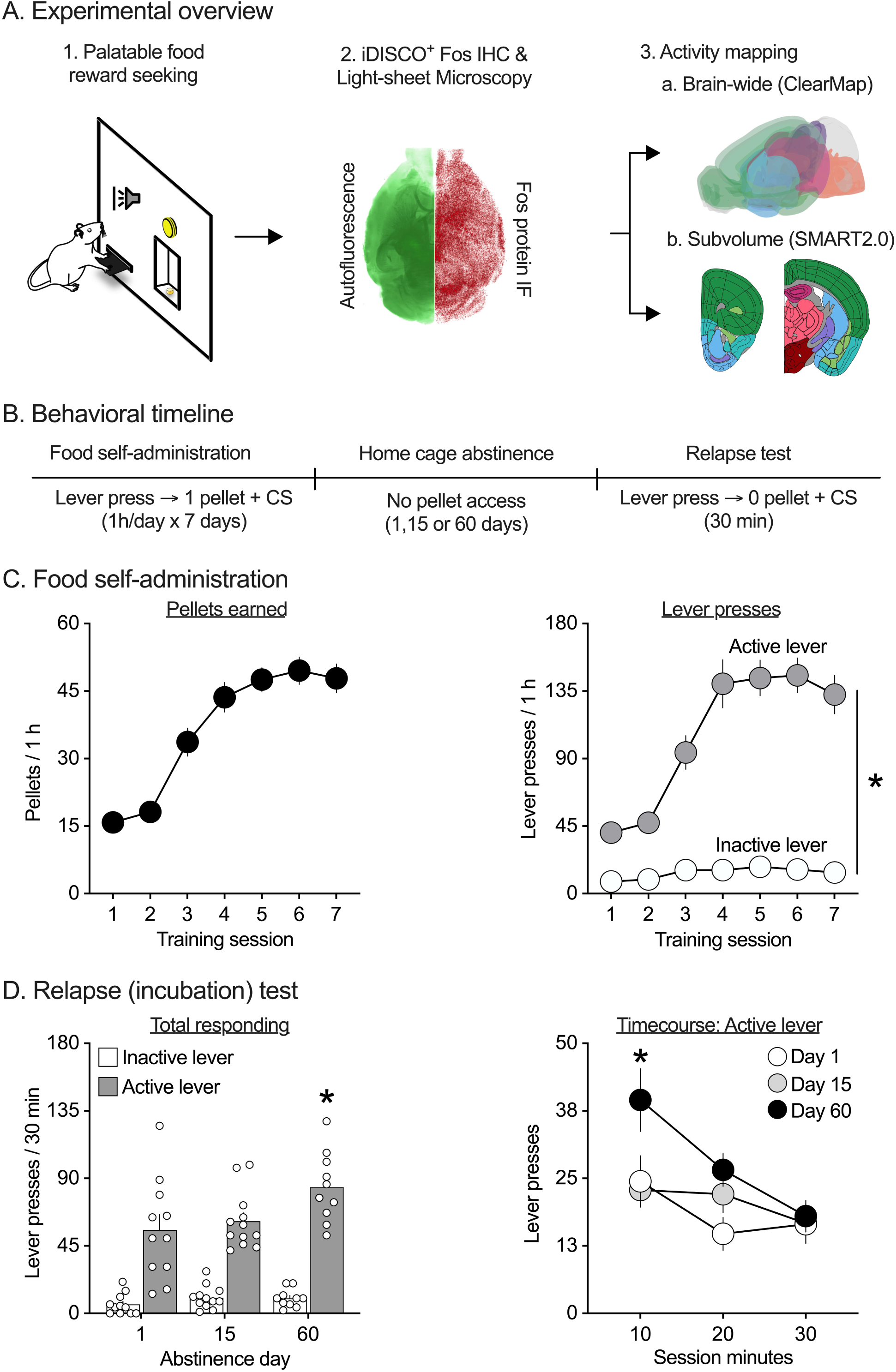
Incubation of palatable food seeking in male CD1 mice. (**A**) Experimental overview. (**B**) Timeline of food self-administration training, abstinence, and relapse tests. (**C**) <u>Food self-administration training.</u> Mice learned to self-administer palatable food pellets over 7 sessions. Mean (±SEM) number of food pellets earned (left panel) and lever presses (right panel) during each 1-h session. * Significant difference (p<0.05) between active and inactive lever presses (n=46). (**D**) <u>Relapse (incubation) test.</u> Responding on active but not inactive lever progressively increased during abstinence. Mean (±SEM) number of lever presses during the entire 30-min relapse test session (left panel) and binned 10-min timecourse of active lever presses (right panel). * Significant difference (p<0.05) from day 1. See Supplementary Table S3 for a detailed listing of all statistical outputs relating to this figure.

#### Food self-administration

The food self-administration procedure is based on our previous study (43). We gave mice free access to regular chow and water in their homecages during this phase and other phases of the experiment. Prior to the operant training sessions, we gave all mice one 30-min session of food-magazine training. During this session mice received 15 evenly spaced (every 2 min) deliveries of a 20-mg palatable food pellet (TestDiet, Catalogue #1811142, 12.7% fat, 66.7% carbohydrate, and 20.6% protein) paired with a 2-s discrete cue light (Food-paired CS). Next, we trained the mice to lever press for palatable food reward during one 1-h session per day. The start of a session was signaled by the illumination of the house light followed 10-s later by the presentation of the central retractable active lever for 60 min. The houselight remained on for the duration of the session and served as a discriminative stimulus that signaled availability of the palatable food upon lever press.

Throughout the session, responses on the active lever were rewarded under a fixed-ratio-1 (FR1), 20-s timeout (TO) reinforcement schedule – active lever presses resulted in illumination of the food-paired CS for 2-s followed by the delivery of a palatable food pellet. Additional active responses during the 20-s timeout had no programmed consequence. Responses on the inactive lever had no programmed consequences throughout the session. We recorded (1) the total number of active lever presses, (2) the total number of inactive lever presses and (3) the total number of food pellet rewards earned during the entire session. Lever-press data for initial training sessions was not recorded for 2 mice due to technical malfunction so we recorded their values as zero to include them in statistical analysis. We gave mice at least 7 training sessions to acquire stable food self-administration behavior before moving to the homecage forced abstinence phase.

We used the above mentioned TestDiet pellets, because both mice and rats prefer this pellet over other nutritional or flavor compositions and show reliable acquisition of food self-administration without any food deprivation (42, 44). Additionally, food-stated CD-1 male mice strongly preferred these food pellets over operant aggression self-administration (43), and food-sated male and female rats strongly prefer these pellets over methamphetamine, heroin, and fentanyl self-administration in rats (45, 46).

#### Forced abstinence

During the abstinence phase, we housed mice in individual cages in the animal facility for 1, 15 or 60 days with no access to palatable food pellets. We gave mice ad libitum access to regular chow and water during this phase and handled them once per week.

#### Relapse tests

Following abstinence, we tested mice for non-reinforced food seeking during a 30-min relapse test. During the test, responses on the active lever resulted in presentation of the Food-paired CS on the same FR1 – 20-s TO reinforcement schedule but were not reinforced with food pellets (extinction conditions). After the test, we returned mice to their homecage for 60-min prior to perfusions and brain tissue collection (n=11 for Day 1, n=12 for Day 15 and n=10 for Day 60). At each incubation timepoint, we also collected brains from food-trained mice directly from the homecage (n=4 for Homecage Day 1, n=6 for Homecage Day 15, n=3 for Homecage Day 60) and collapsed them into a single group (Homecage, n=13) to serve as baseline controls for whole-brain analysis. We matched mice from the four groups for food-reinforced responding during the training phase.

### Whole brain Fos immunohistochemistry (IHC)

We used a modified version of the iDISCO+ protocol for intact mouse brain Fos IHC (41). We processed 37 brains across the 4 groups following perfusions (n=11 for Homecage, n=9 for Day 1, n=9 for Day 15 and n=8 for Day 60) based on perfusion quality, level of intactness, and behavioral data. Details of the sample collection, pretreatment, immunolabeling and clearing steps are provided in the sections below.

#### Sample collection

We anesthetized the mice with isoflurane and perfused them transcardially with 200 ml of 0.1 M phosphate buffered saline (PBS, pH 7.4) followed by 400 ml of 4% paraformaldehyde in PBS (4% PFA, pH 7.4). We extracted brains, post-fixed them for an additional 24-h in 4% PFA at 4°C, and then stored them in PBS with 0.1% sodium azide at 4°C prior to processing.

#### Sample pretreatment with methanol

We used 15 ml conical tubes for sample pretreatment and gently mixed tubes on a rotating mixer (Daigger Scientific, EF24935). We first washed the brains in PBS (3 x 30 min) at room temperature (RT) to remove 4% PFA. We then dehydrated samples in ascending concentrations of methanol (MeOH) in deionized H2O (dH2O) – 0%, 20%, 40%, 60%, 80%, 100%, 100% MeOH (RT, 1.5 h each). Next, we incubated samples in 66% Dichloromethane (DCM)/33% MeOH (RT, 1 x 8 h followed by overnight) for delipidation. We then washed samples in 100% MeOH (RT, 2 x 3 h each), and bleached them by incubating in a chilled H2O2 /H2O/MeOH solution (1 volume 30% H2O2 to 5 volumes 100% MeOH) overnight at 4°C. Next, we rehydrated the samples in descending concentrations of MeOH in dH2O - 80%, 60%, 40%, 20%, 0% MeOH (RT, 1.5 h each). We then washed the samples first in PBS (RT, 1 x 1 h), then three times in a buffer containing PBS with 0.5% TritonX-100 (PTx0.5) at 37°C (2 x 1 h, followed by 1 x overnight).

#### Immunolabeling

We used 1.5 ml Nalgene cryotubes for immunolabeling and 15 ml conical centrifuge tubes for permeabilization, blocking and wash steps. We performed staining over 7 days – we started with a lower initial concentration on day 1 (1° - 1:1000; 2° - 1:500), stepped up the concentration of antibody using booster doses (1° - 1:1000; 2° - 1:500) over the next 4 days, and let the samples incubate at the final concentration (1° - 1:200; 2° - 1:100) for an additional 2 days. We gently mixed the sample containers by rotation and always filled to the top to prevent oxidation. We first incubated the samples in permeabilization buffer containing 78.6% PTx0.5, 1.4% Glycine, and 20% Dimethyl Sulfoxide (DMSO), and then in blocking buffer containing 84% PTx0.5, 6% Normal Donkey Serum (NDS), and 10% DMSO (37°C, 2 d each). Next, we incubated the samples in primary (1°) antibodies (anti-cFos: Phospho-c-Fos (Ser32) (D82C12) XP® Rabbit mAb, Cell Signaling Technology, #5348S Lot 1; RRID#: AB_10557109) diluted in 1° antibody buffer containing 92% PTwH0.5, 3% NDS, and 5% DMSO (37°C, 7 d). We then performed washes in a buffer containing PBS with 0.5% Tween-20, and 10ug/ml Heparin (PTwH0.5) over 4 days (37°C, 4 x 12 h followed by 2 x 1 d). We then incubated the samples in secondary (2°) antibodies (Alexa Fluor® 647 AffiniPure F(ab’)₂ Fragment Donkey Anti-Rabbit IgG (H+L), Jackson ImmunoResearch Labs, 711-606-152, Lot 128806, RRID:AB_2340625; Alexa Fluor® 488 AffiniPure F(ab’)₂ Fragment Donkey Anti-Chicken IgY (IgG) (H+L), Jackson ImmunoResearch Labs, 703-546-155, Lot 127495, RRID:AB_2340375) diluted in 2° antibody buffer containing 97% PTwH0.5, and 3% NDS (37°C, 7 d). Finally, we performed washes in PTwH0.5 over 4 days (37°C, 4 x 8-12 h followed by 2 x 1 d).

#### Clearing

We used 15 ml conical centrifuge tubes for dehydration and delipidation, followed by glass vials with Teflon (PTFE) caps for clearing and refractive index matching. We dehydrated samples using ascending concentrations of MeOH in dH2O – 0%, 20%, 40%, 60%, 80%, 100%, 100% MeOH (RT, 1.5 h each) and performed delipidation using 66% dichloromethane (DCM)+33% MeOH (RT, 2 x 3 h, followed by 1 x overnight). Next, we washed the samples in 100% DCM (RT, 2 x 3 h) to remove MeOH and incubated them in 100% dibenzyl ether (DBE) for clearing and refractive index matching (RT, 2 x 3 h, followed by 1 x overnight).

### Whole-brain imaging and analysis

We imaged stained and cleared intact mouse brains using LSFM. We used the 3D rendering software Arivis Vision 4D (3.0.0) to stitch image tiles, manually corrected coronal alignment where necessary, and exported the images as TIFF files for whole-brain analysis. We analyzed the data using ClearMap (40) and SMART (41) analysis pipelines as described below.

#### Light-sheet fluorescent microscopy imaging (LSFM)

We used a light-sheet microscope (UltraMicroscope II with Infinity Corrected Objective Lenses, Miltenyi Biotec) with an attached camera (Andor Zyla sCMOS), a 1.1x/0.1NA objective (MI PLAN; LaVision BioTec), and a non-corrected dipping cap. Imaging parameters and acquisition order were controlled through ImspectorPro software (v 7.1.4). We mounted cleared Fos-stained brains in coronal orientation (olfactory bulb side up) using a custom sample platform and imaged at 2.2x effective magnification (1.1x objective x 2x magnification slider) in DBE. We acquired images for autofluorescence (Excitation: 488 nm laser, Emission: 535/43 bandpass filter) and Fos-IHC (Excitation: 647 nm laser, Emission: 690/50 bandpass filter) in separate 2 x 1 tiled scans (scan order: z-x-y). We used the following fixed parameters for acquisition: exposure = ∼100 ms; sheet NA = 0.16; sheet thickness = 3.89 um; sheet width = 70%; zoom = 2x; dynamic horizontal focus = 5 (Fos channel only); dynamic horizontal focus processing = blend; merge light-sheet = blend; 488 nm laser power = 20%; 647 nm laser power = 50%. Final image pixel resolution was 2.956 um X x 2.956 um Y x 3 um Z. Resulting tiles were stitched into full size coronal planes using Arivis Vision 4D (3.0.0) and exported as TIFFs. During the analyses, we observed that illumination of the entire coronal plane by the light-sheet during the first imaged tile scan led to significant photo-bleaching of the second imaged tile in all samples. Therefore, we only used the first imaged hemisphere from each sample for analysis and mirrored outputs for all visualizations. We excluded 3 brains due to insufficient clearing and staining, and 2 brains due to technical issues during image acquisition. We analyzed LSFM data for 32 brains across the 4 groups (n=11 for Homecage, n=8 for Day 1, n=8 for Day 15, and n=5 for Day 60).

#### ClearMap analysis

We used the open-source program ClearMap 1.0 (40) for whole-brain volumetric analysis on a dedicated machine (Intel Xeon® CPU E5-2650 v4 @ 2.20GHz x 48; 4 x GeForce GTX 1080 Ti/PCle/SSE2; 256GB RAM). We downsampled autofluorescence image stacks for each sample and registered them to a common 25 µm isotropic serial two-photon (STP) tomography reference template (47). We manually validated registration for each sample by post-hoc inspection of overlaid reference template and post-transformation image stacks in ImageJ. Three brains failed registration and were excluded from further analysis in ClearMap.

Next, we used the spot-detection method in ClearMap to automate Fos+ cell detection across all images. We used the same spot detection filter parameters for Fos+ cell detection across all samples (illumination correction: mean scaling; cell shape detection: threshold (150); Find intensity: Mean and size (3,3,3); Background removal: pixel size (5, 5); DoG filter: pixel size (6,6,11); Detect cell shape: threshold (150) and then applied a voxel size threshold (50, 200000) to constrain the size of detected Fos+ cells.

We validated this Fos+ cell detection procedure against ground truth manual cell detection performed by 2 expert raters across five separate 300 um x 300 um regions of interest (ROIs) from 3 sample image volumes. For each sample, we first isolated an 81 mm coronal image stack (26 image z-stack) at ∼1.46 from bregma along the anterior-posterior axis, and within it selected 5 ROIs encompassing a wide range of Fos+ cell density and background fluorescence signal. Two expert raters performed ground truth manual annotation of Fos+ cells in each FOV using ITK-SNAP software (48) version 3.8.0 (http://www.itksnap.org), resulting in ∼3500 manually annotated Fos+ cells. We used the FIJI image analysis package (49) to overlay the automated ClearMap annotation over the expert annotation and used Analyze Objects and Image Calculator plugins to determine expert-rated Fos+ cell counts, ClearMap-rated Fos+ cell counts and overlap. We calculated precision (ratio of correctly predicted Fos+ cells to all predicted cells), recall (ratio of correctly predicted Fos+ cells to expert annotated Fos+ cells), and F-score (harmonic mean of precision and recall) in Microsoft Excel (Supplementary Figure S1A).

We warped all ClearMap detected Fos+ cells into the reference space by applying transformation coordinates from the registration step and obtained counts for individual brain regions based on the Allen Brain Institute atlas ontology provided with the ClearMap installation package. We extracted Fos+ cell counts for all 1205 annotations and used custom python scripts to generate summed counts within regions of interest (ROIs) for analysis of activity changes between groups.

#### SMART2 analysis

We used an updated version of the open-source R package SMART (41) (SMART2) for expert-guided registration and volumetric activity mapping within two coronal subvolumes selected from ClearMap analysis (subvolume 1: AP +1.55 to AP +1.75 relative to Bregma; subvolume 2: AP -1.08 to AP -1.28 relative to Bregma). SMART extends the WholeBrain analytical framework (38) to volumetric analysis using mouse LSFM datasets and allows user-guided refinement of registration prior to automated Fos+ cell detection. We followed the steps outlined in the updated online tutorial (https://github.com/sgoldenlab/SMART2) for analysis. First, we set up sample information (animal ID, initials, paths, z spacing, registration step (space between z images), most anterior AP coordinate and z image number, most posterior AP coordinate and z image number), and file paths for each sample volume using the functions setup_pl(), im_sort(), and get_savepaths(). Next, we used the function choice() to align the entire sample volume to internal reference atlas plates along the anterior-posterior axis and used the function interpolate() to identify z-stack numbers corresponding to substacks of interest.

Next, we used the interpolated values to select 4 consecutive reference plates (100 mm spacing) for analysis of each coronal subvolume. We selected 2 ‘internal plates’ to register the subvolume of interest (subvolume 1: ABA atlas plate numbers 36 and 37 corresponding to AP +1.6 and AP +1.7; subvolume 2: ABA atlas plate numbers 65 and 66 corresponding to AP -1.13 and AP -1.23) and specified 2 additional ‘outer plates’ to enable identification of Fos+ cells corresponding to 50 mm on either side of the registered internal plates.

This resulted in a final analyzed subvolume of 200 mm thickness in atlas space. We used the functions regi_loop() and filter_loop() for user-guided registration correction between autofluorescence channel images and atlas reference plates. We used the same initial parameters for all samples [alim = c(50, 50), threshold.range = c(50000, 60000L), eccentricity = 999L, Max = 3025, Min = 0, brain.threshold = 400, resize = 0.25, blur = 7, downsample = 2] and applied additional correspondence points to improve the automated registration output.

We then used the function seg_loop() for Fos+ cell detection throughout the subvolume of interest and the function clean_duplicates() to correct for duplicate detection of Fos+ cells across adjacent z-slices. We applied the same segmentation filter parameters for all samples [alim = c(4, 100), threshold.range = c(500, 2200), eccentricity = 300, Max = 2200, Min = 100, brain.threshold = 400, resize = 0.25, blur = 7, downsample = 2]. We validated the automated Fos+ cell detection against ground truth expert annotation as described in the previous section and calculated precision, recall, and F-score in Microsoft Excel (Supplementary Figure S1B).

Finally, we used the forward_warp() function to warp detected Fos+ cells into the reference atlas space and obtained counts for individual brain regions based on the Allen Mouse CCF ontology provided in the WholeBrain package. We extracted Fos+ cell counts and used the get_rois() function to generate summed counts by ROI. We also implemented additional isolate_dataset(), cell_count_compilation(), get_groups(), and voxelize() functions to generate Fos+ cell counts and Fos+ cell density heatmaps for individual samples, and to output summary counts by experimental group. We provide these additional scripts as part of the updated SMART2 package repository (https://github.com/sgoldenlab/SMART2). We also provide a Docker installation image for rapid and user-friendly installation (https://hub.docker.com/repository/docker/goldenneurolab/wholebrain_smart2).

### Statistical analyses

We analyzed behavioral and whole-brain Fos data using IBM SPSS Statistics (version 28) and GraphPad Prism (version 9.3). For the behavioral and whole brain analysis, alpha (significance) level was set at 0.05, two-tailed. We tested the data for sphericity and homogeneity of variance when appropriate. When the sphericity assumption was not met, we adjusted the degrees of freedom using the Greenhouse-Geisser correction. Because our analyses yielded multiple main effects and interactions, we report only those that are critical for data interpretation. See Supplementary Table S2 for a listing of number of subjects/samples included in each phase of the study, and Supplementary Tables S3-S11 for statistical analyses and summary statistics.

#### Behavior

We analyzed two behavioral measures during food self-administration training - (1) the total number of lever presses on active and inactive lever (denoted as *lever presses*), and (2) the total number of food pellet rewards earned (denoted as *rewards*) during a 1-h self-administration session. Following training, we tested separate groups of mice for palatable food reward-seeking after 1, 15 or 60 days of abstinence and analyzed non-reinforced responding (*lever presses*) during the 30-min food seeking test. We describe the within- and between-subjects factors in the mixed ANOVAs we used to analyze the behavioral data in the Results section and in Supplementary Table S3.

#### ClearMap analysis

We processed the 32 brains using the ClearMap pipeline, of which 3 failed the registration to the reference atlas. We analyzed the remaining 29 brains (n=9 for Homecage, n=7 for Day 1, n=8 for Day 15 and n=5 for Day 60) as described below. We computed Fos+ cell counts within regions at two levels of the atlas hierarchy - (1) 10 major anatomical divisions, and (2) 56 subregions across the brain based on hierarchical relationships defined in the Allen Mouse Common Coordinate Framework (50). We selected regions within each level such that there were no parent-child relationships and/or overlapping spatial footprints between them. We transformed the raw Fos+ cell counts to Z-scores prior to the statistical analyses to normalize the data and account for differences in volume across regions of interest. We computed Z-scores for each region of interest relative to the homecage group’s values using the formula z = (x – μ) / σ, where x is the sample Fos+ cell count, μ is the mean Fos+ cell count of the homecage group, and σ is the standard deviation of the homecage group. For each level, we first used 2-way mixed ANOVA (GLM procedure in SPSS) with the within-subject factor of Region and the between-subjects factor of Group (Homecage, Day 1, Day 15, Day 60). We followed up on significant main effects and interactions with 1-way ANOVAs within each region and used Tukey HSD test for post-hoc comparisons between Groups. We provide the statistical outputs for all analysis pertaining to ClearMap in Supplementary Tables S4 and S5.

#### SMART2 analysis

In addition to the 29 brains used for ClearMap analysis, we used manual registration correction within each subvolume to recover and analyze brains that failed ClearMap registration. We analyzed 32 brains across the 4 groups (n=11 for Homecage, n=8 for Day 1, n=8 for Day 15 and n=5 for Day 60) for subvolume 1 and 31 brains across the 4 groups (n=10 for Homecage, n=8 for Day 1, n=8 for Day 15 and n=5 for Day 60) for subvolume 2. We extracted Fos+ cell counts within regions of interest at three levels of the atlas hierarchy - (1) major anatomical regions (subvolume 1: 5 regions; subvolume 2: 8 regions), (2) main subregions (subvolume 1: 18 subregions; subvolume 2: 28 regions), and (3) internal subdivisions within the subregions (subvolume 1: 43 subdivisions; subvolume 2: 64 subdivisions). Similar to ClearMap analysis, we selected regions within each level such that there were no parent-child relationships and/or overlapping spatial footprints between them. We transformed the raw Fos+ cell counts to Z-scores prior to statistical testing. For each level, we first used 2-way mixed ANOVA (GLM procedure in SPSS) with the within-subject factor of Region and the between-subjects factor of Group (Homecage, Day 1, Day 15, Day 60). We followed up on significant main effects and interactions with 1-way ANOVAs within each region and used Tukey HSD test for posthoc comparisons between Groups. We also generated spatial maps of Fos+ cell densities (mean across group) for each subvolume to aid visualization of differences in activation patterns between groups. We provide statistical outputs for all analysis pertaining to SMART2 subvolumes 1 and 2 in Supplementary Tables S6-S8 and S9-S11, respectively.

## Results

The experimental design and timeline of behavioral training, intact mouse brain activity labeling, and brain-wide activity mapping is shown in Figure 1A. See Supplementary Table S2 for a listing of number of subjects/samples included in each phase of the study.

### Incubation of palatable food seeking in male CD1 mice

We determined whether the time-dependent increases in food seeking during abstinence (incubation of food craving), previously observed in rats (3, 19), generalize to mice. The experimental timeline is shown in Figure 1A-B. We trained food sated CD-1 male mice to lever press for high carbohydrate food pellets for 7 days. Next, we tested different groups for relapse to food seeking in the presence of contextual (food self-administration chamber) and discrete (light cue paired with food delivery) cues after 1, 15, or 60 days of homecage forced abstinence. Statistical outputs for all analyses pertaining to training and relapse tests are provided in tabular format as Supplementary Table S3.

#### Training phase

The mice showed reliable food self-administration as indicated by increased responding on the food-paired lever during the daily sessions (Figure 1C). The repeated-measures ANOVA of *rewards*, which included the within-subjects factor of Training session (sessions 1-7), showed a significant effect of this factor (F3.2,144.7= 67.2, p<0.001). The repeated measures ANOVA of *lever presses*, which included the within-subjects factors of Training session and Lever (inactive, active), showed a significant interaction between the two factors (F3.6,160.3= 28.9, p<0.001).

#### Relapse (incubation) tests

Non-reinforced presses on the previously active lever were significantly higher after 60 abstinence days than after 1 or 15 days (Figure 1D), indicating ‘incubation of food craving.’ The mixed ANOVA of total lever presses, which included the between-subjects factor of Abstinence day (1, 15, 60) and Lever, showed a significant interaction between the two factors (F2,30=3.4, p=0.045). This incubation effect was primarily due to higher lever presses in the day 60 group during the first 10 min of the test session (Figure 1D right panel). Post-hoc group differences (Tukey test) are depicted in Figure 1D.

### Unbiased intact brain-wide activity mapping of incubation of palatable food-seeking

We processed brains using the ClearMap pipeline and extracted Fos+ cell counts within regions of interest (ROIs) at two levels of the atlas hierarchy - (1) 10 major anatomical divisions (Fig. 2A, Fig. 2B right panel), and (2) 56 subregions of interest (Fig. 2C) based on hierarchical relationships defined in the Allen Mouse Common Coordinate Framework (50). We selected regions within each level such that there were no parent-child relationships and/or overlapping spatial footprints between them and performed z-score normalization (relative to homecage group mean) for each ROI prior to statistical analyses (Fig. 2C, center and left panels).

**Figure 2.**
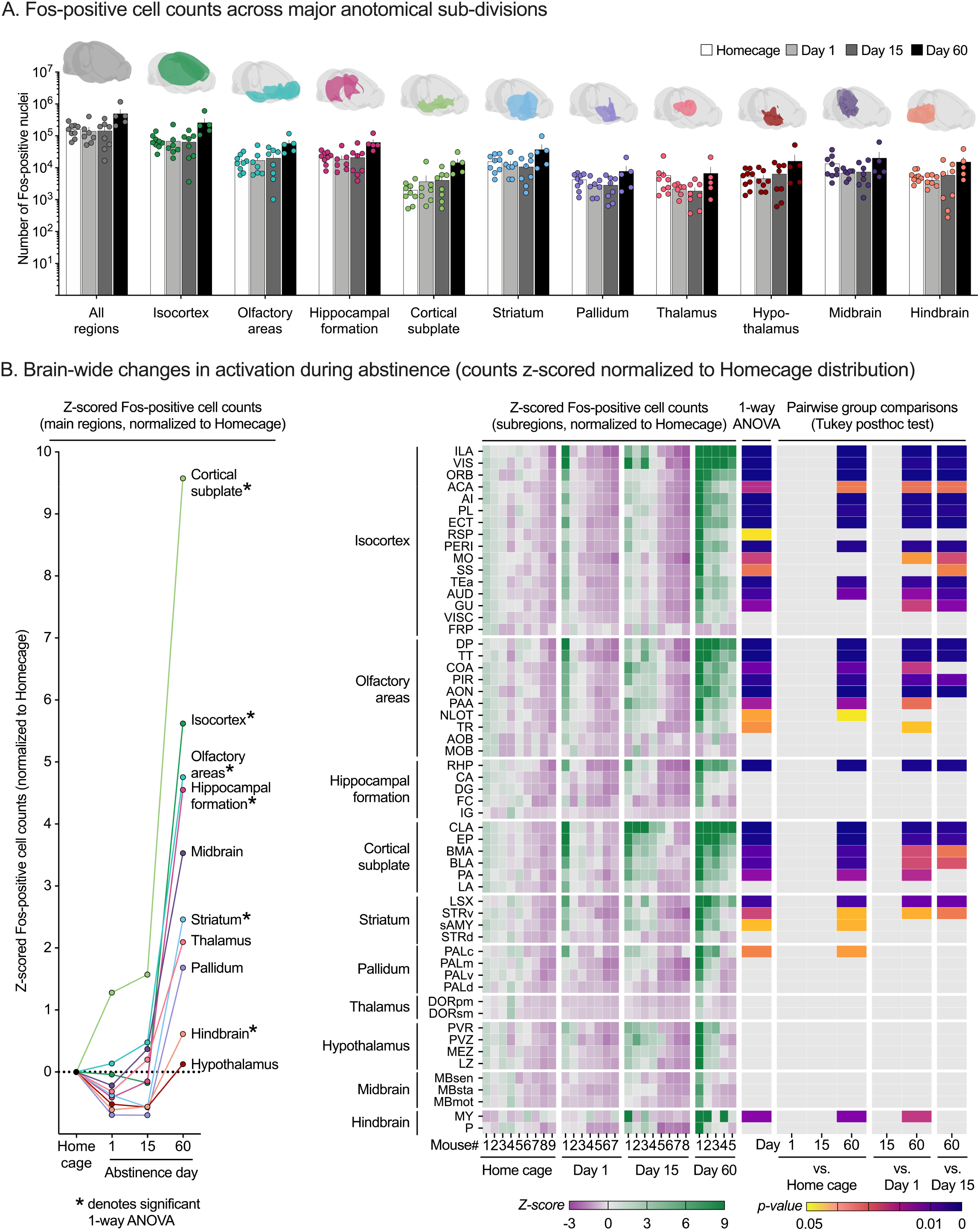
Unbiased brain-wide activity mapping of incubated relapse to palatable food-seeking using ClearMap. (**A**) Mean (±SEM) Fos+ cell counts across the whole brain and for 10 major anatomical sub- divisions. (**B**) Brain-wide changes in activation during abstinence. Raw Fos+ cell counts for each region are z-score normalized to the homecage group distribution for statistical analysis. Mean z-scored cell counts for 10 major anatomical sub-divisions showing time-dependent changes in activation pattern in multiple brain regions induced by the relapse tests (left panel). * Significant differences (p<0.05) 1-way ANOVA. Heatmap of individual z-scored Fos+ cell counts, 1-way ANOVA p-value and Tukey HSD pairwise group comparison p-values for 56 subregions across the analyzed brain volume (right panel). Individual data is sorted by group and ranked in descending order of activation level within each group. Subregions are organized by 10 parent anatomical sub-divisions and ranked in descending order of mean day 60 group activation level. See Supplementary Tables S4-S5 for a detailed listing of all statistical results associated with this figure.

#### Main regions

For Level 1, the mixed ANOVA, using the between-subjects factor of Group (homecage, day 1, day 15, day 60) and the within-subjects factor of Region, showed significant main effects of Group (F3,25=4.4, p=0.013) and Region (F1.9,47.7=18.2, p<0.001), and an interaction between the two factors (F5.7,47.7=6.1, p<0.001). Follow up one-way ANOVAs (between-subjects factor Group) were significant for 6 out of 10 tested regions (denoted by * in Figure 2B, left panel). These data indicate increased brain-wide activation after the relapse test after 60 abstinence days, but not 1 or 15 days. Detailed statistical reporting of the ClearMap analysis at Level 1 are provided in Supplementary Table S4.

#### Sub-regions

For Level 2, the mixed ANOVA showed significant main effects of Group (F3,25=6.0, p=0.003), and Sub-region (F1.8,45.3=18.7, p<0.001) and an interaction between the two factors (F5.4,45.3=8.0, p<0.001). Follow up one-way ANOVAs (between-subjects factor Group) were significant for 33 out of 56 tested subregions, reflecting selective increased activation following relapse test after 60 but not 1 or 15 days of abstinence. The most activated areas were olfactory, cortical, cortical subplate, and striatal subregions - collectively designated as ‘corticostriatal’, with sparse subregional activation in the retrohippocampal region of the hippocampus and medulla of the hindbrain. Increased Fos expression in the day 60 group was not observed in subregions of thalamus, hypothalamus, and midbrain structures. Detailed statistical reporting of ClearMap analysis at Level 2 are provided in Supplementary Table S5.

### Targeted analysis of the corticostriatal coronal subvolume using SMART2

We processed a 200 μm thick coronal subvolume spanning AP +1.55 to AP +1.75 relative to Bregma using the SMART2 pipeline. Spatial maps of group-wise Fos+ cell densities within the subvolume are shown in Figure 3A to aid visualization of differences in activation patterns between groups. We extracted Fos+ cell counts within regions of interest (ROIs) at three levels of the atlas hierarchy - (1) 5 major anatomical regions (Fig. 3B), (2) 18 main subregions (Fig. 3C) and (3) 43 internal subdivisions within the subregions (Fig. 3D). We performed z-score normalization (relative to homecage mean) for each ROI prior to statistical analysis. One-way ANOVA and Tukey HSD p-values for all Level 2 ROIs and a subset of Level 3 ROIs are shown as heatmaps in Figure 3C and Figure 3D.

**Figure 3.**
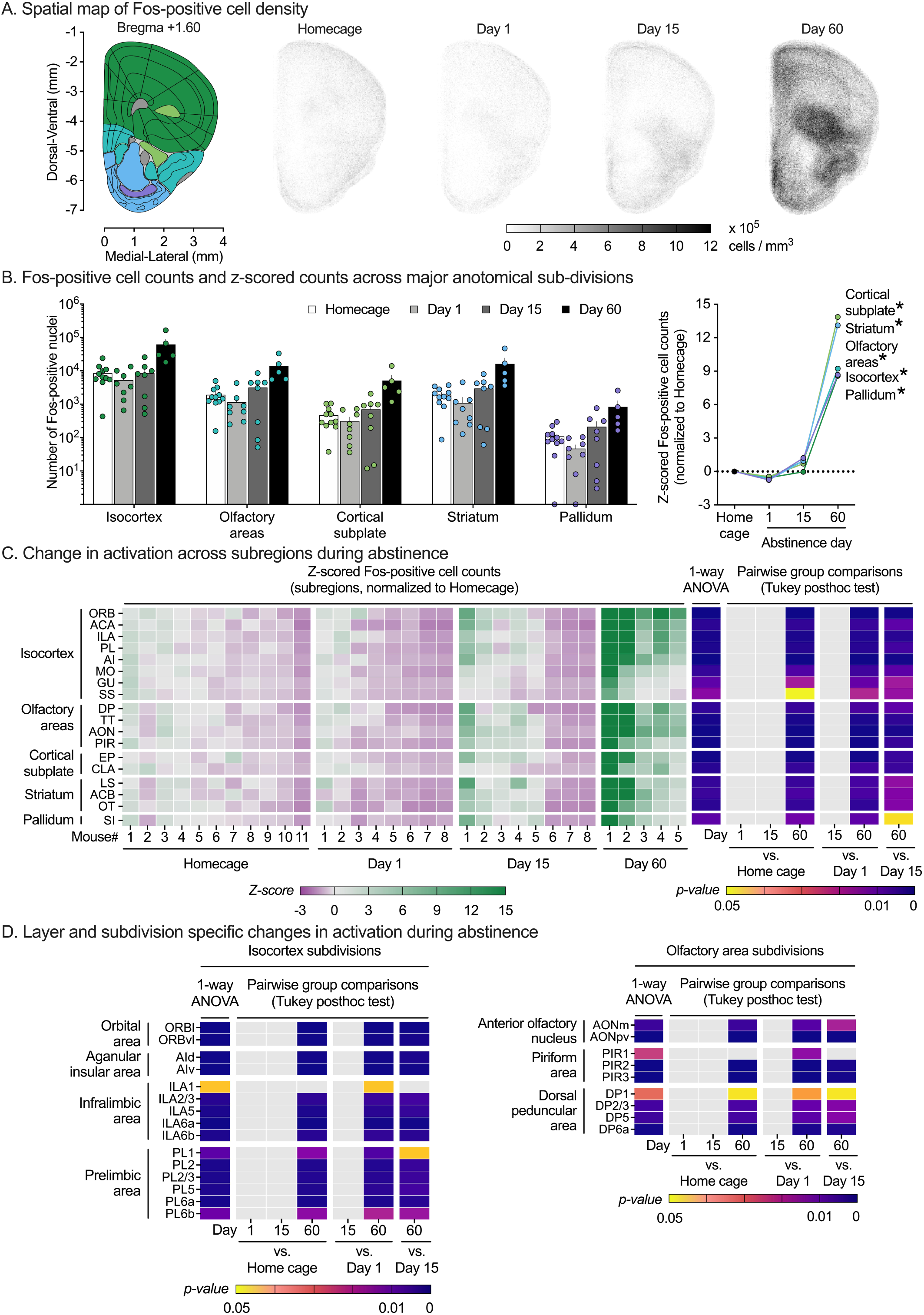
Targeted analysis of corticostriatal coronal subvolume (AP +1.55 to AP +1.75 relative to Bregma) using SMART2. (**A**) Spatial map of Fos+ cell density. Grayscale intensity of individual points (20 μm x 20 μm x 200 μm voxel) represents mean cell density in cells/mm^3^ within AP +1.55 to AP +1.75 coronal subvolume. (**B, left**) Fos+ cell counts for 5 major anatomical regions within the subvolume. (**B, right**) Z-score normalized counts (normalized to home cage group distribution) for 5 major anatomical regions within the subvolume. * Significant differences (p<0.05) 1-way ANOVAs. (**C**) Changes in activation across the subvolume during abstinence. Fos+ cell counts for each region are z-score normalized to homecage group distribution for statistical analysis. Heatmap of individual z-scored Fos+ cell counts (**left**), 1-way ANOVA p-value (**middle**) and Tukey HSD pairwise group comparisons p-values (**right**) for 18 sub-regions within the analyzed coronal subvolume. Subregions are organized by 5 parent anatomical sub-divisions and ranked in descending order of mean day 60 group activation level. (**D**) Layer and subdivision specific changes in activation. Heatmap of 1-way ANOVA p-value and Tukey HSD pairwise group comparison p-values for selected sub-region layers or subdivisions within Isocortex (**left**) or Olfactory areas (**right**). See Supplementary Tables S6-S8 for a detailed listing of all statistical results associated with this figure.

#### Main regions

For Level 1, the mixed ANOVA showed significant main effects of Region (F1.6,4.7=6.5, p=0.006) and Group (F3,28=6.9, p=0.001), and an interaction between the two factors (F4.7,44.0=4.7, p=0.002). Follow-up one-way ANOVAs were significant for all 5 tested regions, reflecting increased activation following the relapse test after 60 but not 1 or 15 days of abstinence. Detailed statistical reporting for SMART2 analysis at Level 1 are provided in Supplementary Table S6.

#### Sub-regions

For Level 2, the mixed ANOVA showed significant main effects of Sub-region (F1.2,32.7=9.5, p=0.003) and Group (F3,28=7.2, p=0.001), and an interaction between the two factors (F3.5,32.7=6.3, p=0.001). Follow-up one-way ANOVAs were significant for all 18 tested subregions, reflecting increased activation following the relapse test after 60 but not 1 or 15 days of abstinence. Detailed statistical reporting for SMART2 analysis at Level 2 are provided in Supplementary Table S7.

#### Sub-divisions

For Level 3, the mixed ANOVA showed significant main effects of Sub-division (F1.2,32.2=10.3, p=0.002) and Group (F3,28=7.1, p=0.001), and an interaction between the two factors (F3.5,32.2=6.5, p=0.001). Follow-up one-way ANOVAs were significant for all 43 tested subdivisions, reflecting increased activation following the relapse test after 60 but not 1 or 15 days of abstinence. Detailed statistical reporting for SMART2 analysis at Level 3 are provided in Supplementary Table S8.

### Targeted analysis of the thalamocortical coronal subvolume using SMART2

We processed a 200 μm thick coronal subvolume spanning AP -1.08 to AP -1.28 relative to Bregma using the SMART2 pipeline. Spatial maps of group-wise Fos+ cell densities within the subvolume are shown in Figure 4A to aid visualization of differences in activation patterns between groups. We extracted Fos+ cell counts within ROIs at three levels of the atlas hierarchy - (1) 8 major anatomical regions (Fig. 4B), (2) 28 main subregions (Fig. 4C), and (3) 64 internal subdivisions within the subregions (Fig. 4D). One-way ANOVAs and Tukey HSD p-values for all Level 2 ROIs and a subset of Level 3 ROIs are shown as heatmaps in Figure 4C and Figure 4D.

**Figure 4.**
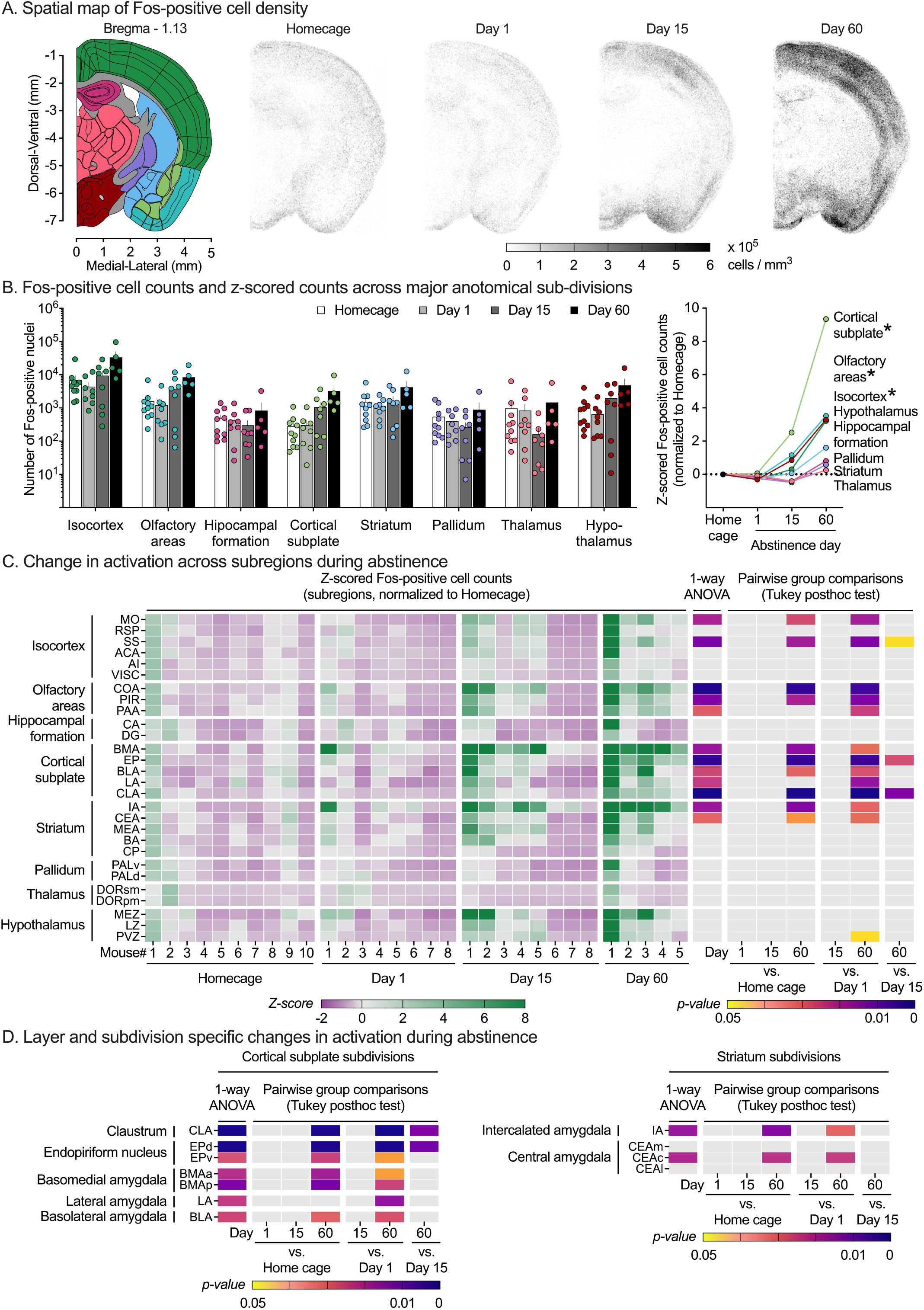
Targeted analysis of thalamocortical coronal subvolume (AP -1.08 to AP -1.28 relative to Bregma) using SMART2. (**A**) Spatial map of Fos+ cell density. Grayscale intensity of individual points (20 μm x 20 μm x 200 μm voxel) represents mean cell density in cells/mm^3^ within AP -1.08 to AP -1.28 coronal subvolume. (**B, left**) Fos+ cell counts for 8 major anatomical regions within the subvolume. (**B, right**) Z-score normalized counts (normalized to home cage group distribution) for 5 major anatomical regions within the subvolume. * Significant differences (p<0.05) 1-way ANOVAs. (**C**) Changes in activation across the subvolume. Fos+ cell counts for each region are z-score normalized to home cage group distribution for statistical analysis. Heatmap of individual z-scored Fos+ cell counts (left), 1-way ANOVAs p-values (middle) and Tukey HSD pairwise group comparisons p-values (**right**) for 28 sub-regions within the analyzed coronal subvolume. Subregions are organized by 8 parent anatomical subdivisions and ranked in descending order of mean day 60 group activation level. (**D**) Layer and subdivision specific changes in activation. Heatmap of 1-way ANOVAs p-values and Tukey HSD pairwise group comparison p-values for selected subregion layers or subdivisions within Cortical subplate (**left**) or Striatum (**right**). See Supplementary Tables S9-S11 for a detailed listing of all statistical results related to this figure.

#### Main regions

For Level 1, the mixed ANOVA showed significant main effects of Region (F1.2,32.3=14.0, p<0.001) and Group (F3,27=3.2, p=0.040), and an interaction between the two factors (F3.6,32.3=5.5, p=0.002). Follow-up one-way ANOVAs were significant for 3 of 8 tested regions, reflecting increased activation following the relapse test after 60 but not 1 or 15 days of abstinence. Detailed statistical reporting for SMART2 analysis at Level 1 are provided in Supplementary Table S9.

#### Sub-regions

For Level 2, two-way ANOVA showed significant main effects of Sub-region (F1.1,30.8=9.5, p=0.003) and Group (F3,27=3.4, p=0.033), and an interaction between the two factors (F3.4,30.8=3.5, p=0.024). Follow-up one-way ANOVAs were significant for 12 of 18 tested subregions, reflecting increased activation following relapse test after 60 but not 1 or 15 days of abstinence. Detailed statistical reporting for SMART2 analysis at Level 2 are provided in Supplementary Table S10.

#### Sub-divisions

For Level 3, the mixed ANOVA showed significant main effects of Region (F1.2,31.2=7.8, p=0.007) and Group (F3,27=3.1, p=0.044), and an interaction between the two factors (F3.5,31.2=3.0, p=0.038). Follow-up one-way ANOVAs were significant for 20 of 64 tested subdivisions, reflecting increased activation following the relapse test after 60 but not 1 or 15 days of abstinence. Detailed statistical reporting for SMART2 analysis at Level 3 are provided in Supplementary Table S11.

## Discussion

We used unbiased intact-brain mapping of Fos expression to investigate brain-wide activation patterns during incubation of palatable food seeking in mice. Relapse to food seeking was higher after 60 abstinence days than after 1 or 15 days, indicating incubation of food seeking. More importantly, unbiased whole-brain analysis of Fos expression using ClearMap showed a strong induction of neural activity across multiple brain regions that mirrored incubation of food seeking after 60, but not 15, days of abstinence. Targeted coronal slice analysis of Fos expression using SMART2 replicated and validated the time-dependent increases in activation patterns within corticostriatal and thalamocortical subvolumes and enabled detailed analysis of subdivision and layer-specific changes during abstinence. Overall, our data indicate that incubation of palatable food craving in male mice correlates with widespread activation of many brain regions beyond those previously implicated in incubation of food or drug craving (see Supplementary Table S1).

### Outbred mice show incubation of palatable food seeking similar to outbred rat models

Food-seeking in the male mice increased following 60, but not 15, days of abstinence from food self-administration training. Previous studies in rats reported incubation of food seeking but with a different time course; robust incubation of sucrose craving was observed in rats after 7, 15, and 30 abstinence days, but not 60 days (5). Incubation of food craving has also been reported in rats trained to self-administer standard chow pellets after 30 abstinence days (14), and high-carbohydrate pellets after 21 abstinence days (15). By extending the model to mice we were able to leverage mouse-optimized procedures to investigate brain-wide activation patterns of incubation of food craving. A question for future research is whether a similar timecourse of incubation of food craving (and pattern of brain activation) will be observed in CD-1 female mice.

Notably, we used outbred CD-1 mice in these experiments rather than more commonly used inbred C57BL/6J. In our experience, outbred and hybrid mice exhibit a more complex spectrum of behavior and acquire learned behavior more robustly than their inbred counterparts(51–54).

Additionally, an inbred genetic background limits the generalizability of genotype-phenotype relationships(55) and display highly heritable stain-specific phenotypes in brain volume, scalar diffusion tensor imaging metrics, and quantitative connectomes(56). Similarly, contrary to the assumed relationship, meta-analysis of coefficients of variation do not find evidence of greater trait stability in inbred mice than outbred mice, and hybrid mice show enhanced properties desired for neurobiological research such as reduced anxiety-like behavior, improved learning, and enhanced long-term spatial memory(57). Taken within the context of developing a resource whole brain atlas for incubation of food craving in mice, outbred populations likely provide a more generalizable and robust platform.

### Brain-wide patterns of increased neural activation following incubation of food seeking during abstinence

Previous targeted region-by-region analyses in rats have identified several brain regions that are activated during incubation of food and drug craving during abstinence (1, 3, 21, 28, 29).

However, an unbiased brain-wide interrogation of regions engaged during ‘incubated’ reward seeking has not previously been performed. We used a modified version of the iDISCO+ procedure to label active (Fos+) neurons in intact mouse brains and employed light-sheet fluorescence microscopy to image these potentially behaviorally relevant neurons across the entire brain volume at single cell resolution. Our unbiased ClearMap analysis revealed a time-dependent increase in activation of multiple brain regions (e.g., prelimbic, infralimbic, orbitofrontal, and insular cortices, central amygdala and basolateral amygdala, ventral and dorsal striatum) similar to that shown in rats following incubation of food and drug seeking (58–64) (see Supplementary Table S1). However, this time-dependent increase of neural activity was not restricted to these previously identified incubation-related regions. Indeed, over half of the tested main regions (6 out of 10 main anatomical divisions, and 33 of 56 subregions), including most regions within the isocortex, olfactory areas, cortical subplate, and striatum, showed statistically significant increased activation after 60, but not 15 abstinence days. Additionally, compared to ’resting state’ activity in the homecage, most regions showed an initial dip in activation following food seeking tests on days 1 and 15, which preceded strong induction on day 60.

Of note, it is possible that a small number of the statistically significant activated regions reflect false positive results, because we did not correct the statistical analyses for multiple one-way ANOVA comparisons. On the other hand, the fact that statistically significant increases in Fos expression in multiple brain regions were detected using a relatively small n per group (n=5 to 8 for abstinence days 1, 15, and 60) speaks to the robustness of our results.

Two recent studies used a similar unbiased approach to investigate brain-wide activation (assessed by Fos) patterns of short-term alcohol abstinence after two-bottle choice and chronic intermittent ethanol vapor exposure (65) and acute withdrawal following experimenter-administered psychostimulants (66). They used hierarchical clustering techniques to demonstrate a strong decrease in modularity after abstinence/withdrawal compared to drug-naïve controls and employed graph theory approaches to identify hub regions that might drive this functional restructuring. It is possible that this observed decrease in modularity might be a result of the recruitment of networks of regions like that seen in our study following food seeking in mice.

However, it is important to note that in these studies, brains were collected directly from homecage and not after behavioral testing. Thus, the observed changes likely reflect shifts in ’resting-state’ functional brain architecture and not behaviorally evoked differential network engagement during reward seeking.

### Targeted subvolume analysis reveals variations in time-dependent brain activation patterns

We followed-up on our global volumetric ClearMap findings by analyzing activation patterns within two coronal volumes using an updated version of the SMART pipeline (41). This approach allowed us to use expert-guided registration and Fos-segmentation to (1) isolate ClearMap effects to specific coronal plates along the anterior-posterior axis, (2) validate our data against previous incubation studies that used selected coronal slices for Fos-mapping, (3) include brains that failed ClearMap due to physical damage during processing, and (4) extend our analysis to subdivisions and layers within these subvolumes for future mechanistic investigation. In agreement with the ClearMap analyses, SMART2 analysis identified several subregions with increased activation (following the relapse tests on abstinence day 60) within the isocortex, olfactory areas, cortical subplate, and striatum across the two selected subvolumes, several of which have been previously identified after incubation of food and drug seeking (e.g., prelimbic cortex, infralimbic cortex, orbitofrontal cortex, basolateral amygdala, central amygdala nucleus, nucleus accumbens, somatosensory cortex) and drug seeking (e.g., prelimbic cortex, infralimbic cortex, agranular insular area, basolateral amygdala, central amygdalar nucleus, nucleus accumbens) in rat models (58-64, 67-69) (see Supplementary Table S1). However, while some identified subregions showed similar increases in activation across both subvolumes (e.g., somatomotor and somatosensory areas, claustrum, piriform area), others were only present in one subvolume (e.g., prelimbic and infralimbic cortex, basolateral and central amygdala) or showed differential engagement along anterior-posterior axis (e.g., anterior cingulate and agranular insular cortices). Even within a subvolume and subregion, activation patterns were not uniform but sometimes graded across layers (e.g., layers of piriform area, dorsal peduncular area, infralimbic and prelimbic areas) or isolated to specific subdivisions (e.g., central but not medial or lateral subdivisions of central amygdala), suggesting different degrees of engagement across multiple brain regions and likely circuits after 60 abstinence days.

## Conclusions

We demonstrated that incubation of palatable food reward seeking is accompanied by an induction of neural engagement in multiple brain regions, many of them extend beyond the traditional brain areas and circuits involved in incubation of food and drug craving. We extend the rat incubation of food seeking model to male CD1 mice and leverage mouse-specific unbiased whole-brain staining, clearing, and analysis pipelines to generate a single-cell resolution whole-brain atlas of incubation food seeking during prolonged abstinence. The results of our study suggest that the overarching neural mechanism underlying incubation of reward seeking is more anatomically widespread than suggested by the published literature and likely not localized to a particular brain area or circuit. Whatever neural mechanisms mediate incubation of reward seeking, our findings suggest that these mechanisms affect acute neural responses throughout the brain, either through widespread alterations in all these brain areas or in key brain areas that regulate brain-wide circuitry. And finally, the ‘incubation atlas’ here provides a mineable dataset for better understanding system-level alterations related to incubation of food craving and relapse.

## Acknowledgments

We want to thank members of the Hope, Shaham, and Golden labs for their support and insight during all stages of this study. We also thank Dr. Jennifer M. Bossert and Dr. Ida Fredriksson for assistance with whole-brain immunohistochemical assays, and Dr. Carlos A. Mejias-Aponte and Vadim Kashtelyan for technical assistance with sample processing and light sheet fluorescence microscopy.

## Supplementary Information

**Fig. S1.**
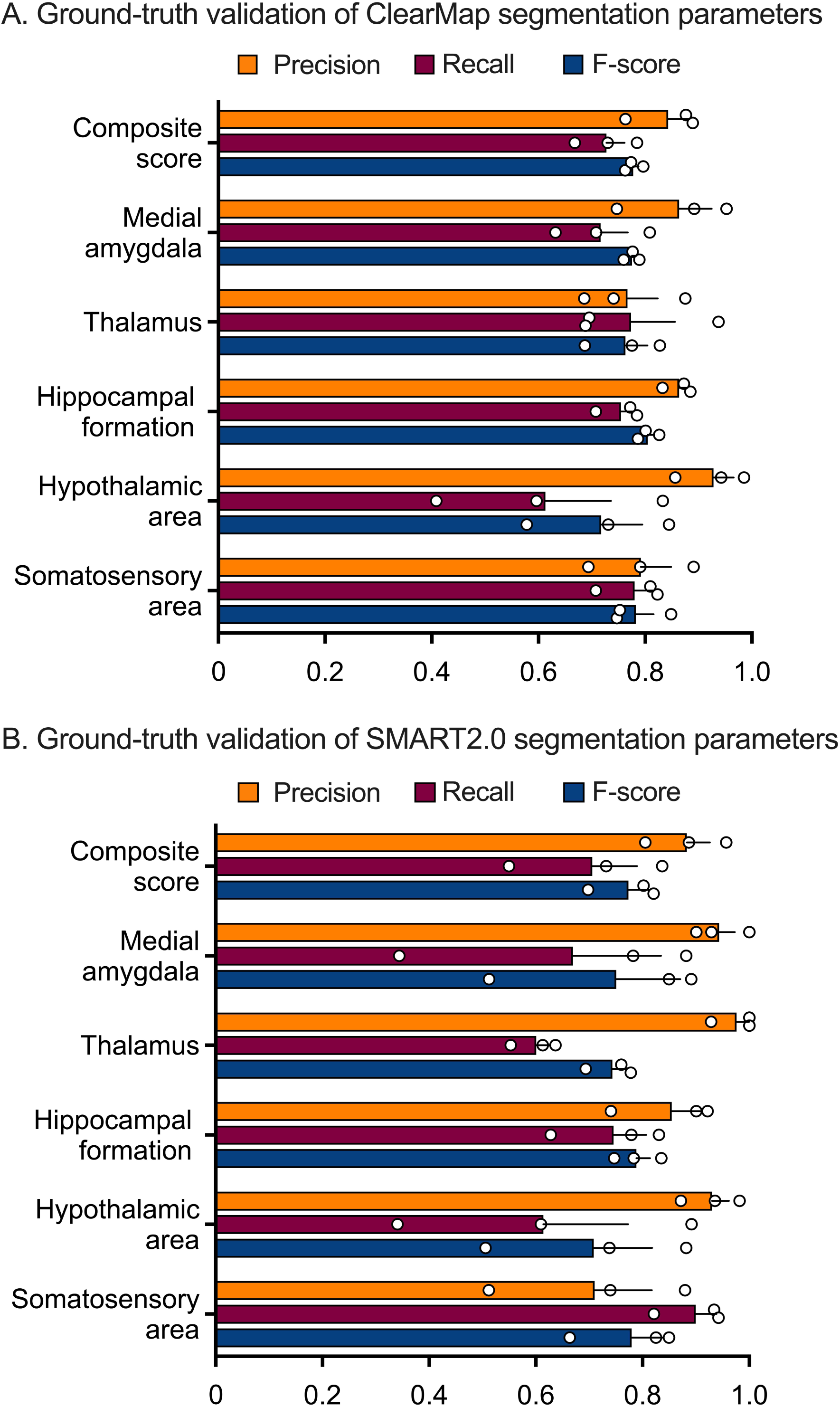
Validation of ClearMap (A) and SMART2 (B) segmentation parameters against expert Fos+ cell annotation

**Table S1.**
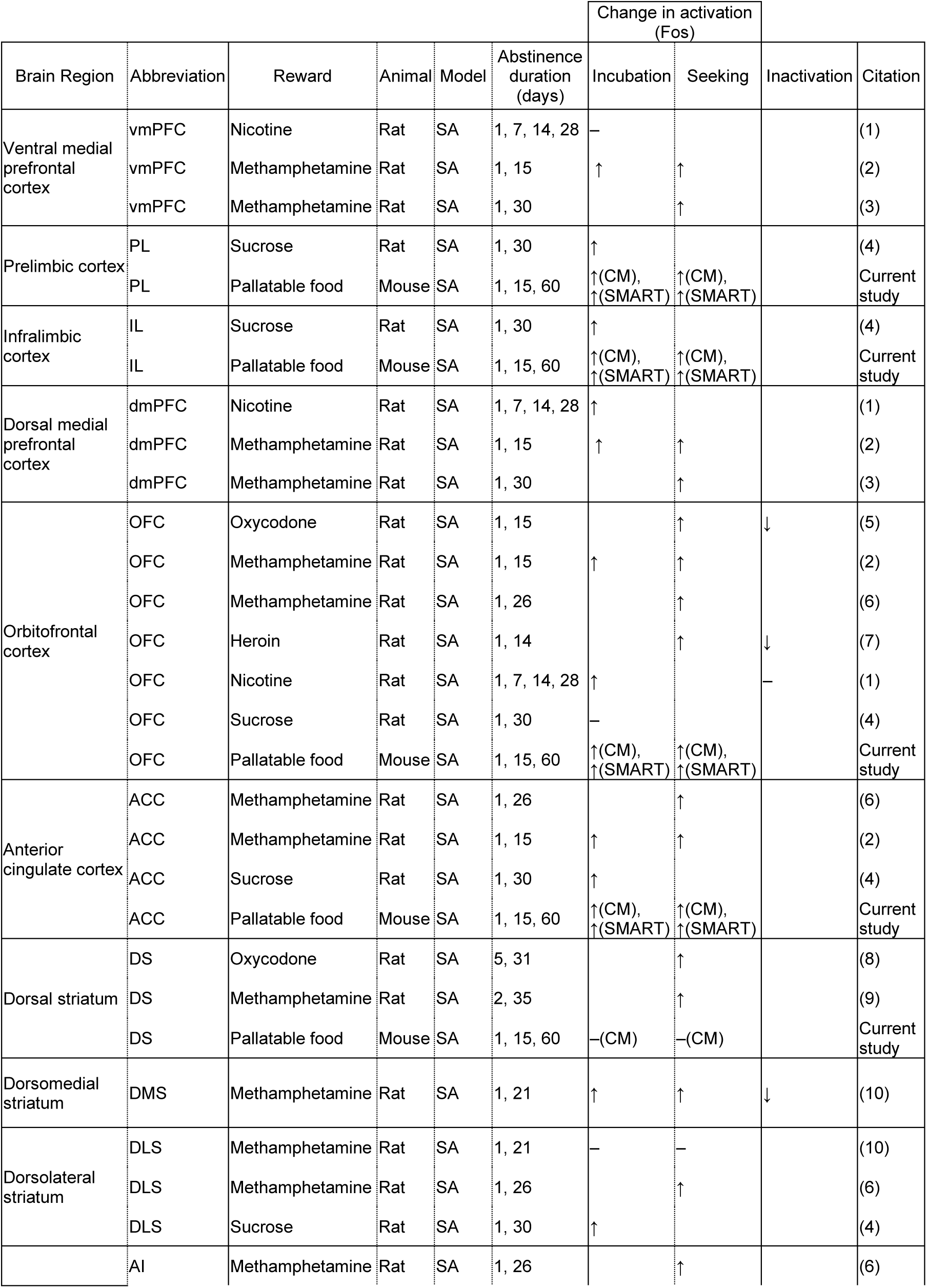
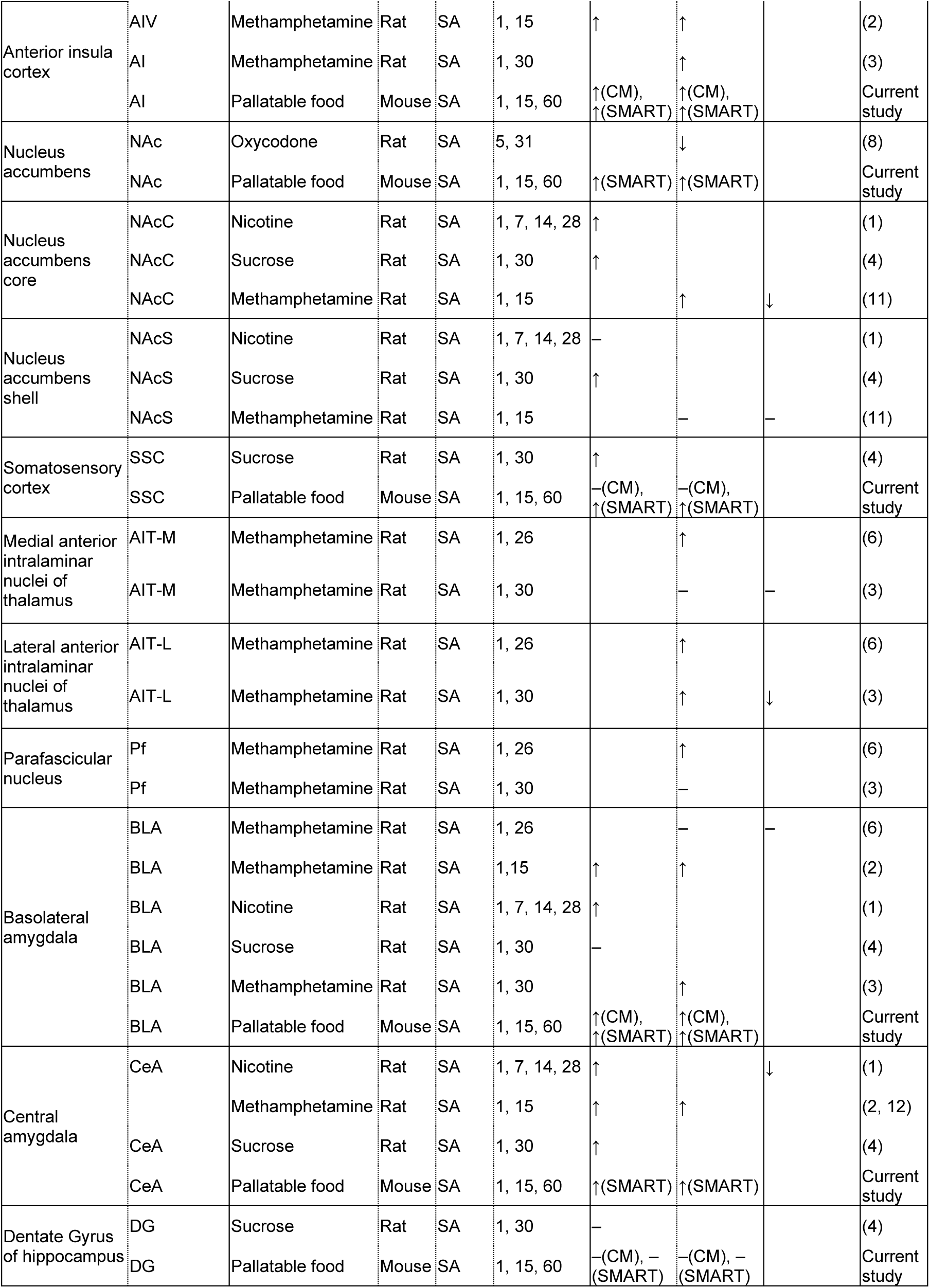
Incubation-relevant regions identified using targeted section-based activity mapping approaches.

**Table S2.**
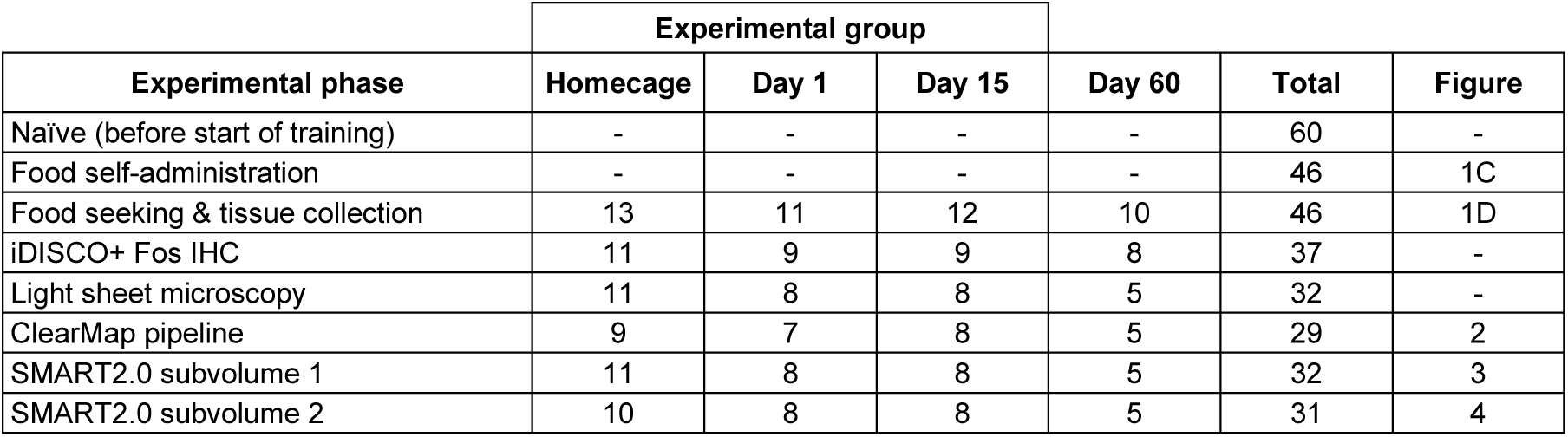
Number of subjects that completed each phase of the study.

| Experimental phase | Experimental group |  |  |  | Total | Figure |
| --- | --- | --- | --- | --- | --- | --- |
|  | Homecage | Day 1 | Day 15 | Day 60 |  |  |
| Naïve (before start of training) | - | - | - | - | 60 | - |
| Food self-administration | - | - | - | - | 46 | 1C |
| Food seeking & tissue collection | 13 | 11 | 12 | 10 | 46 | 1D |
| iDISCO+ Fos IHC | 11 | 9 | 9 | 8 | 37 | - |
| Light sheet microscopy | 11 | 8 | 8 | 5 | 32 | - |
| ClearMap pipeline | 9 | 7 | 8 | 5 | 29 | 2 |
| SMART2.0 subvolume 1 | 11 | 8 | 8 | 5 | 32 | 3 |
| SMART2.0 subvolume 2 | 10 | 8 | 8 | 5 | 31 | 4 |

**Table S3.**
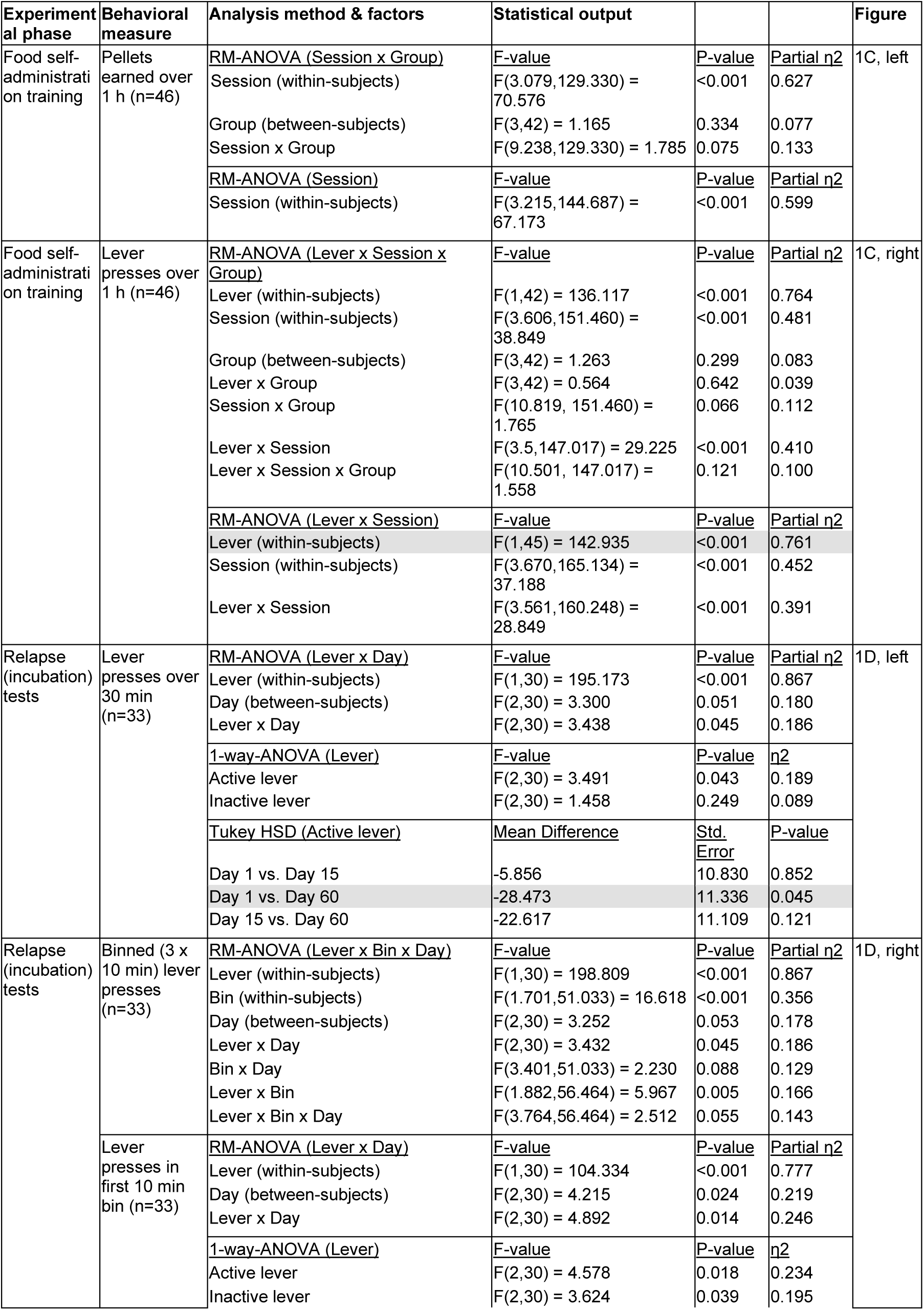
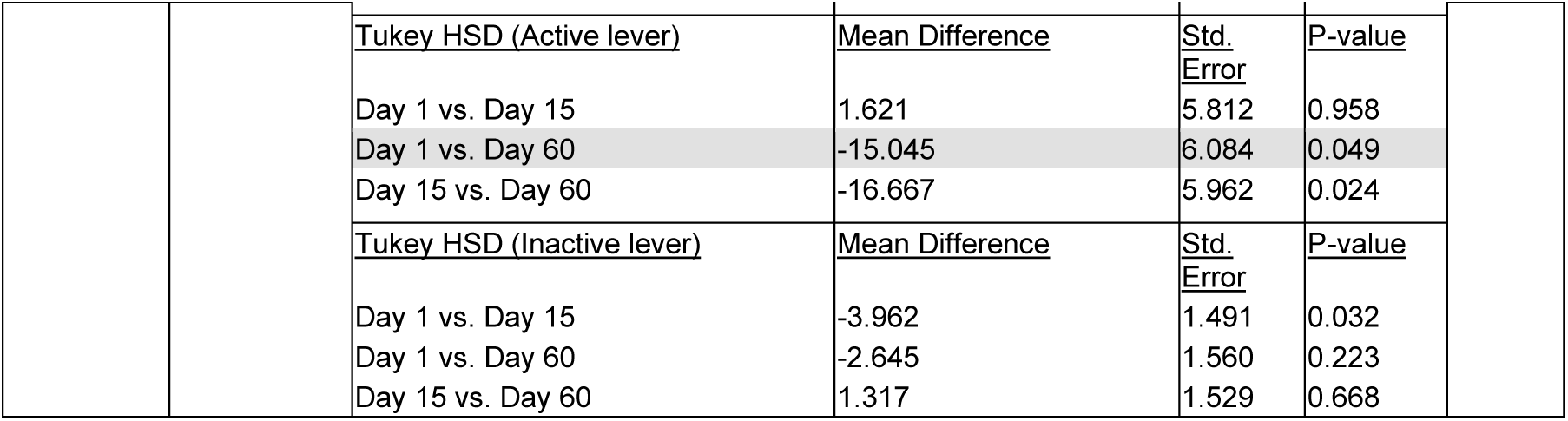
Statistical output for Figure 1: (analyses shown in figure 1 are highlighted in grey)

**Table S4.**
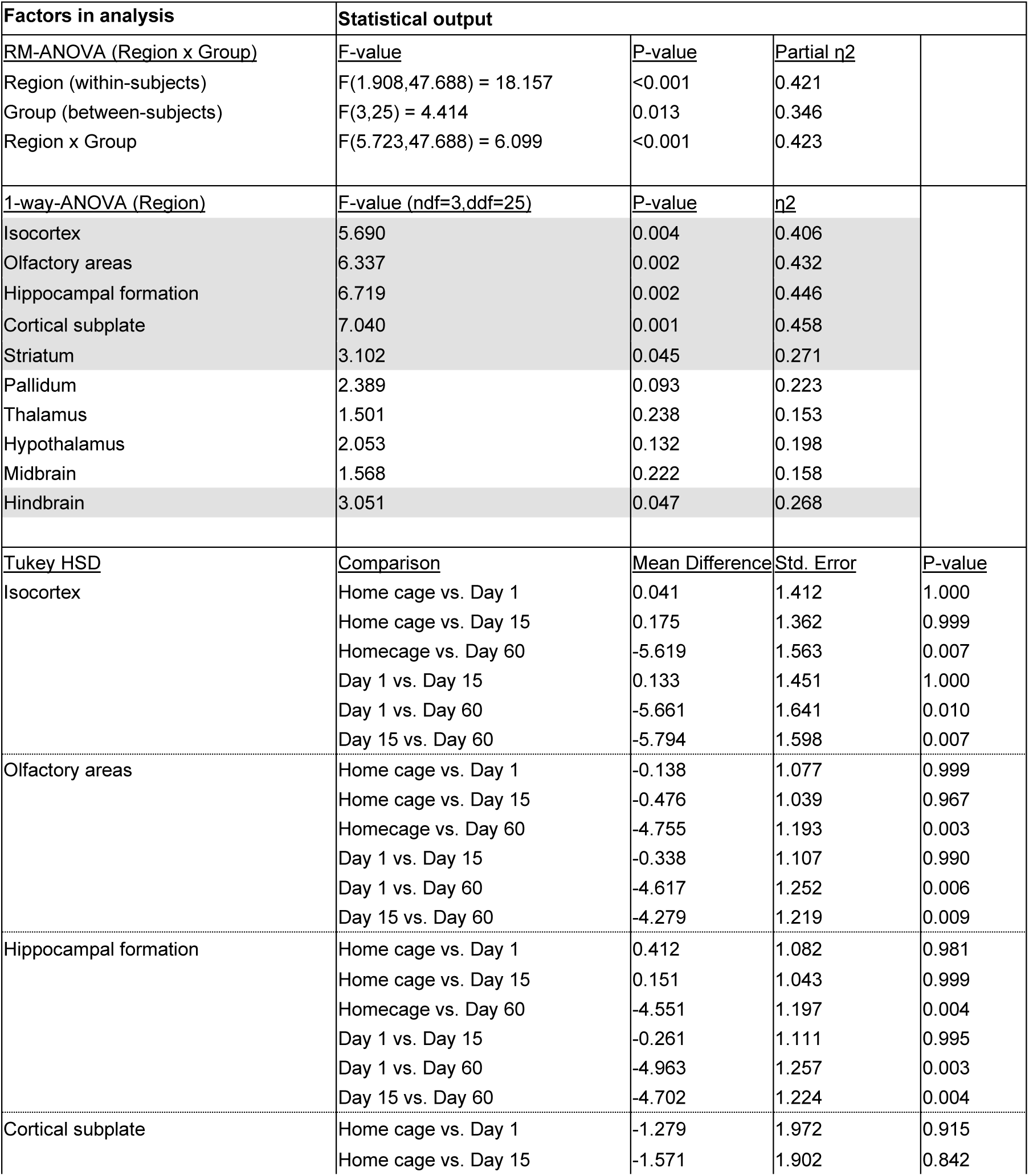
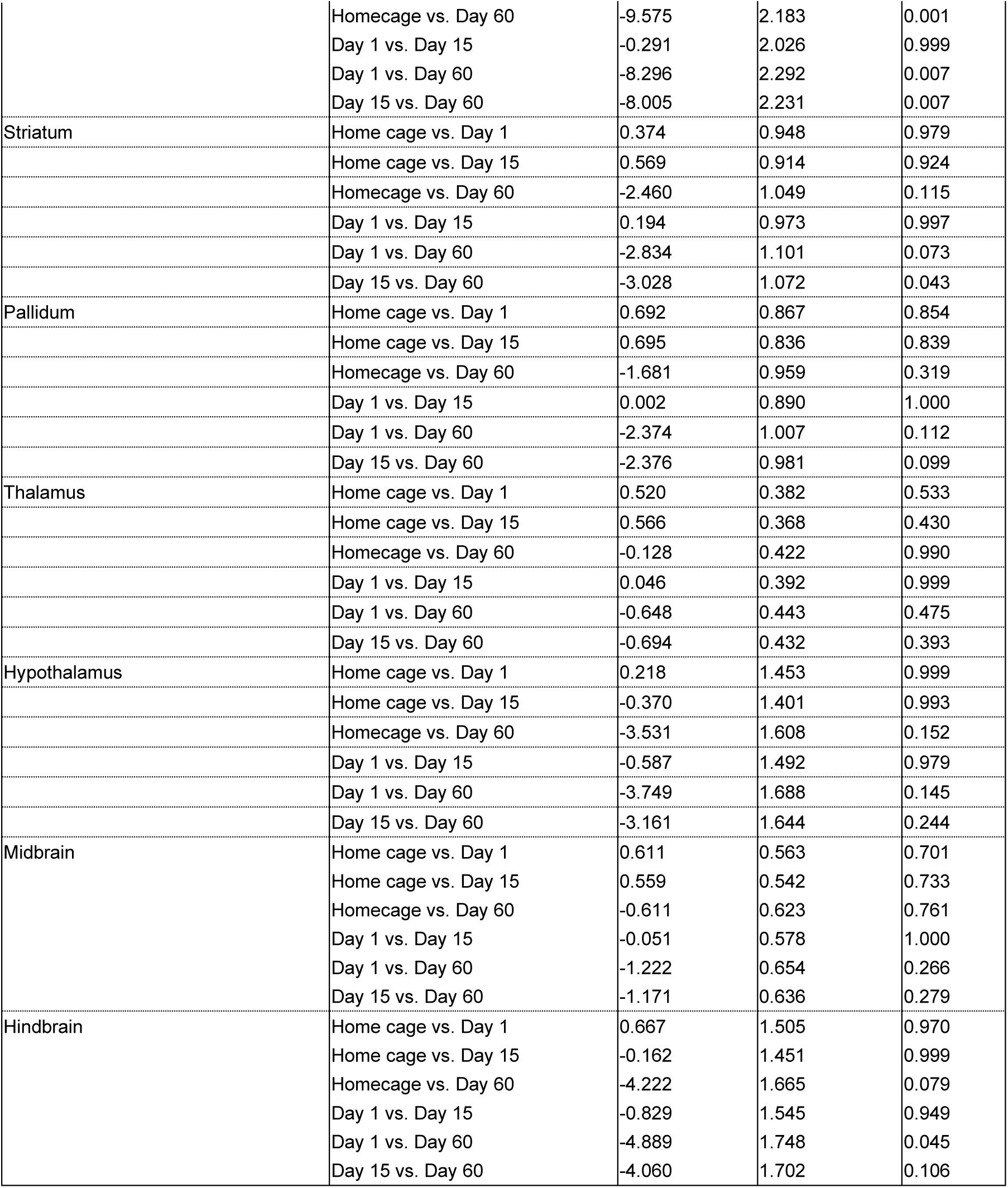
Statistical output for Figure 2B, left panel : Analysis of Z-scored counts from 10 major anatomical regions (n=29, analyses shown in figure 2B are highlighted in grey)

**Table S5.**
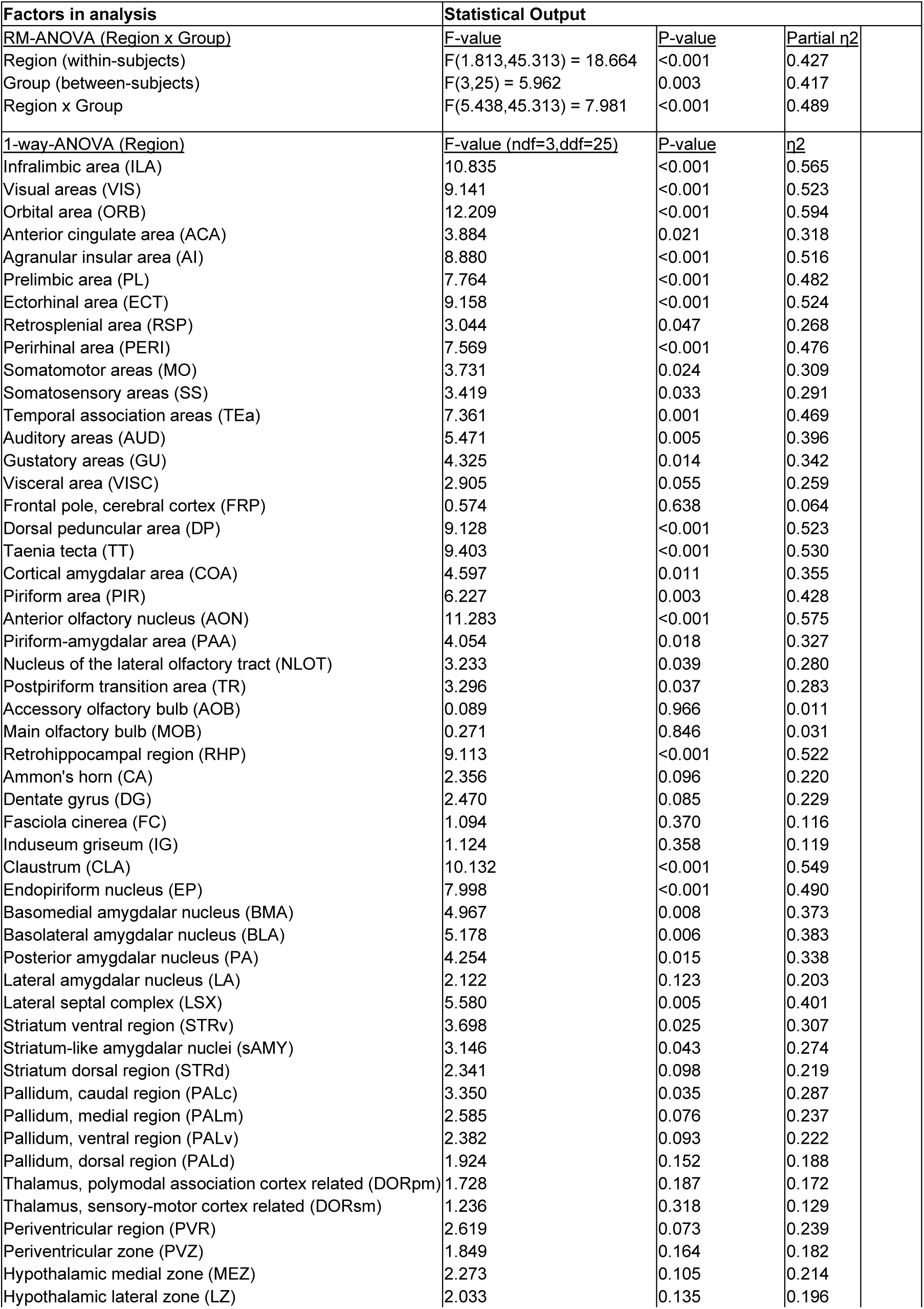
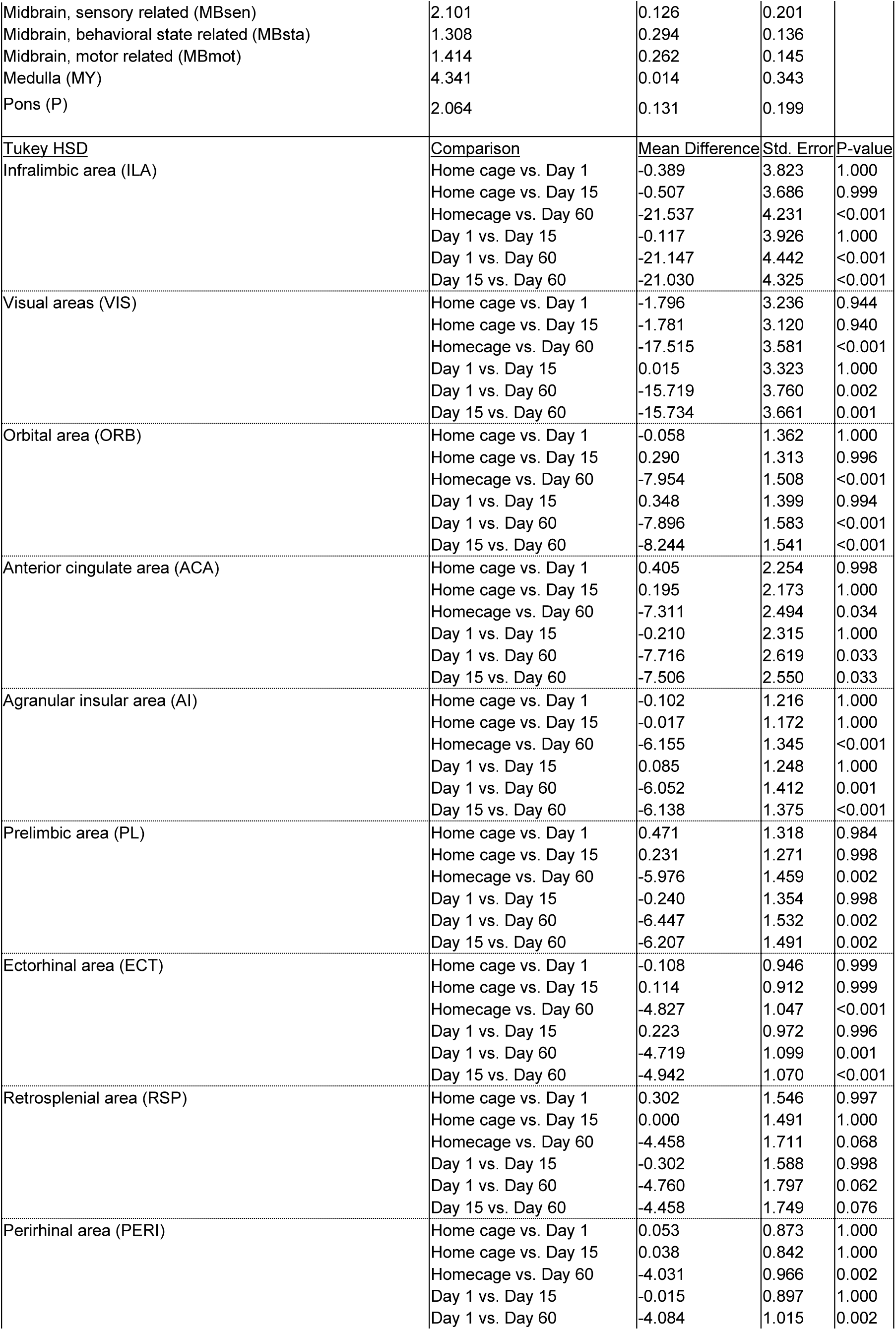
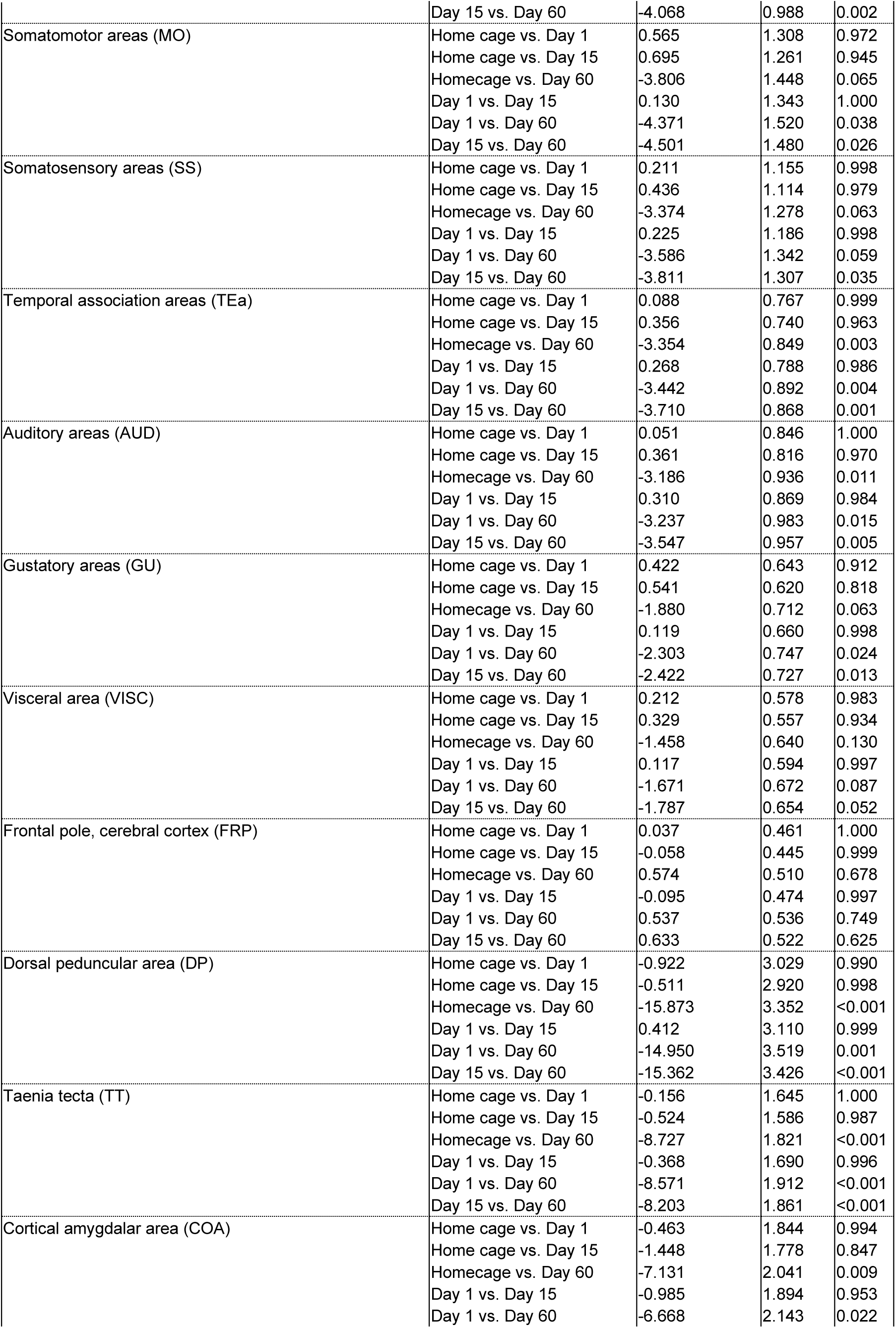
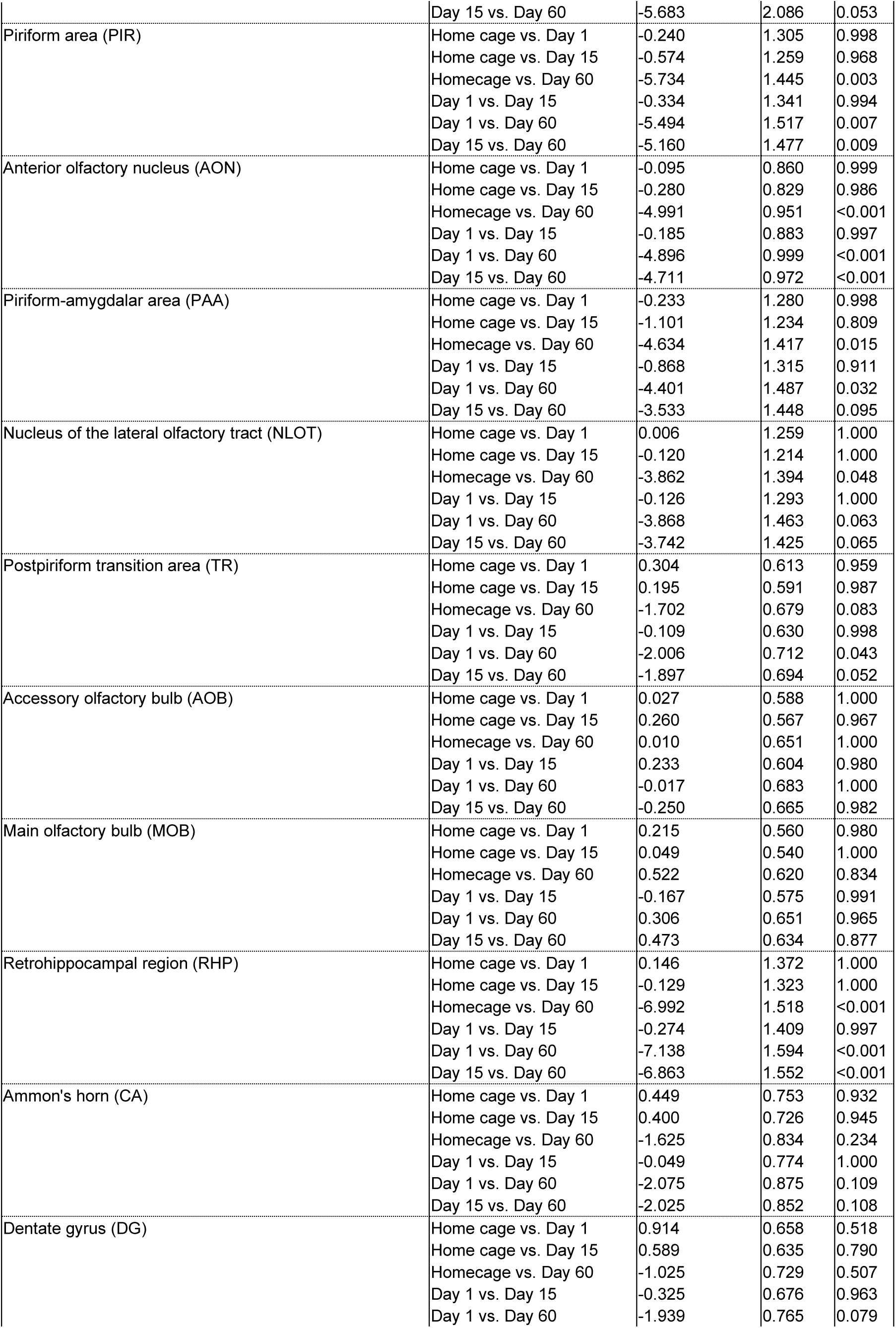
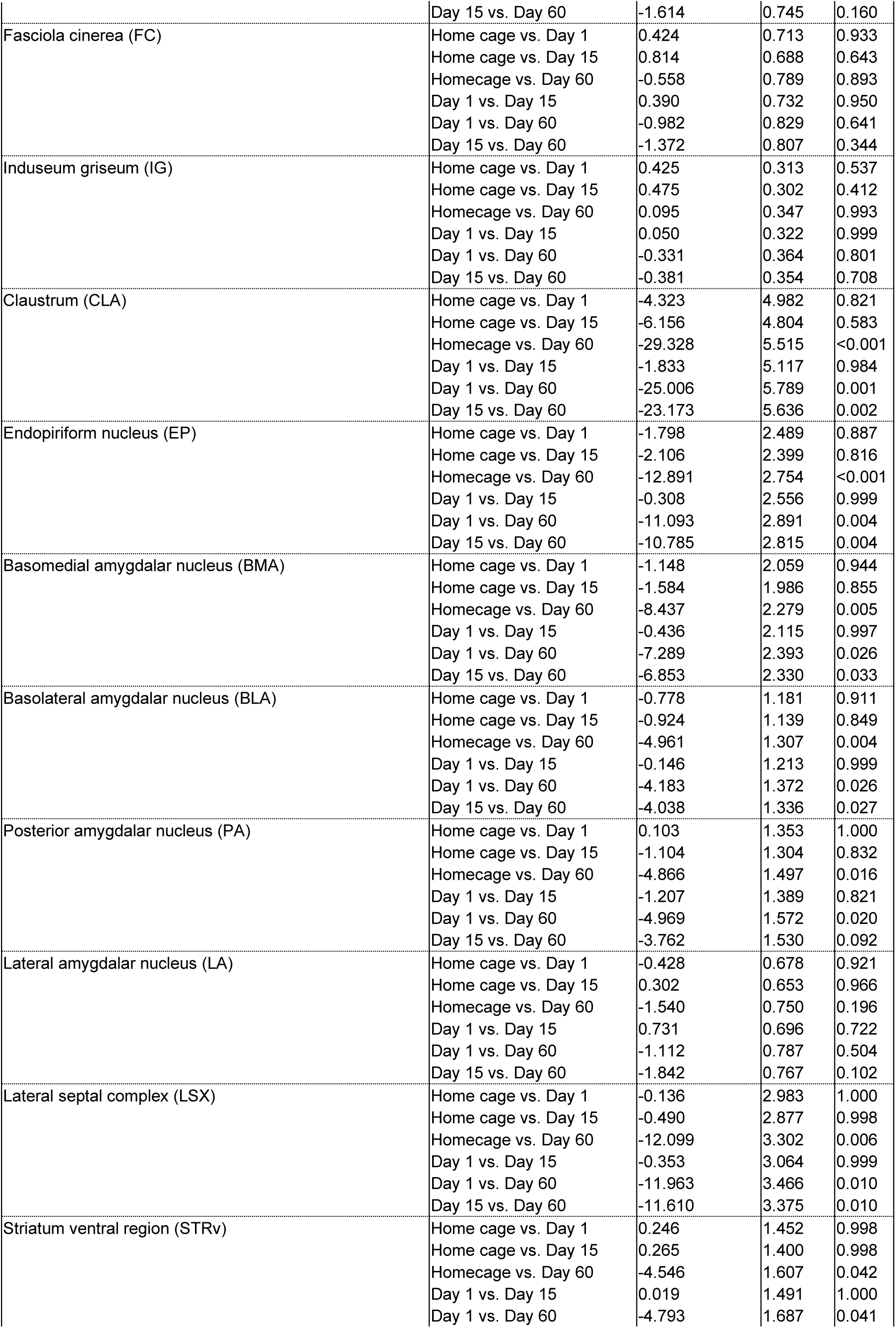
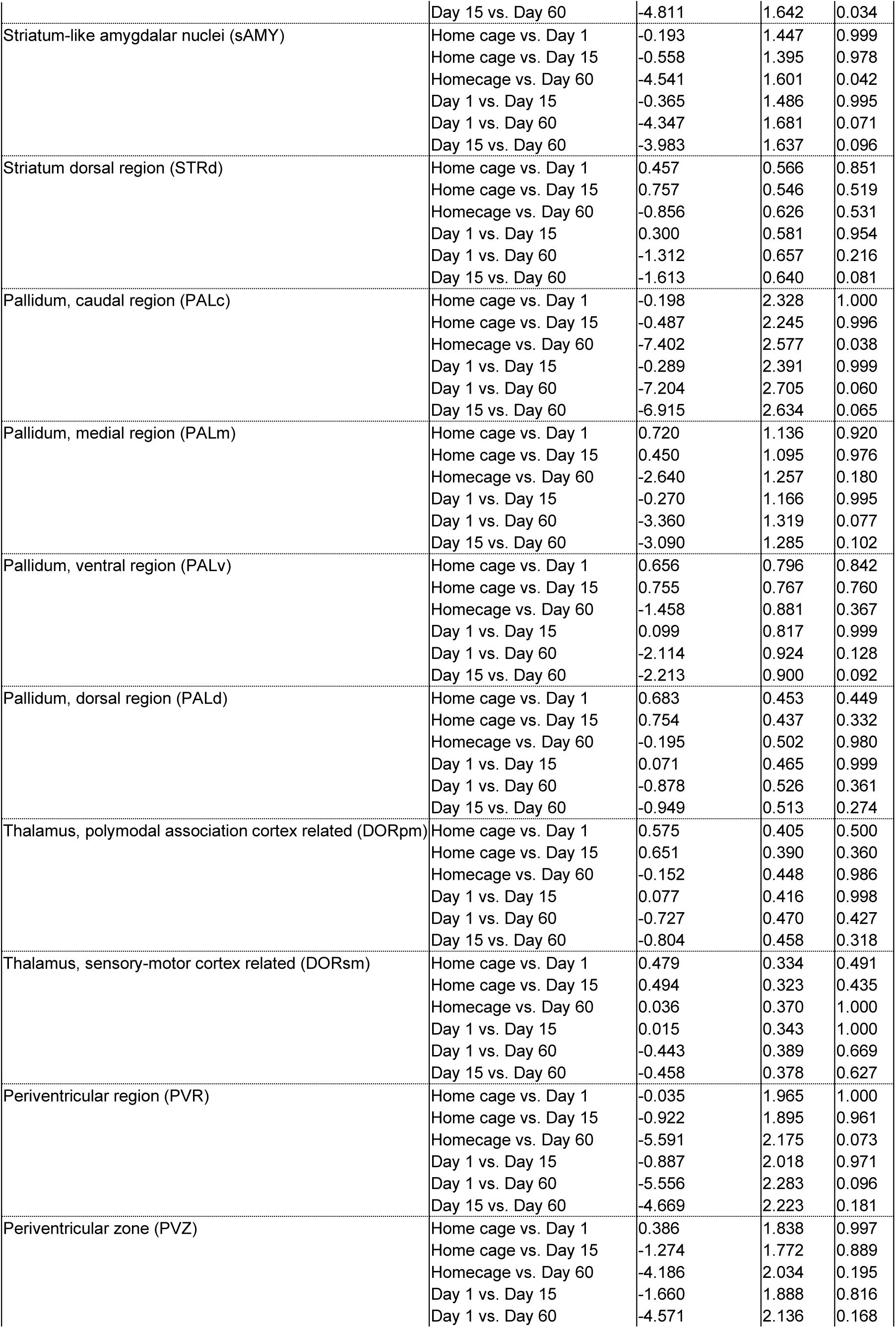
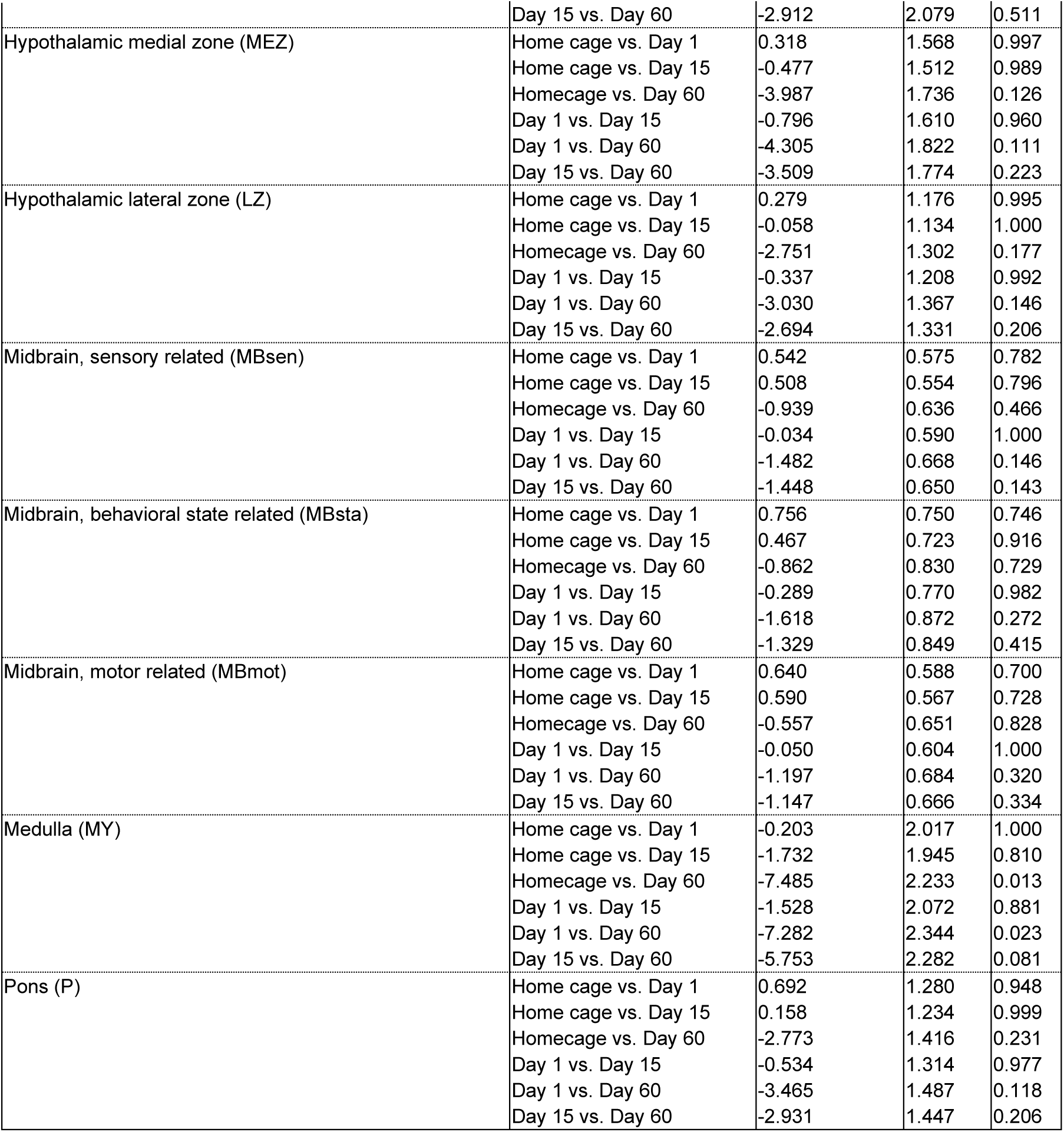
Statistical output for Figure 2B, right panel: Analysis of Z-scored counts from 56 subdivisions of 10 major anatomical regions (n=29)

| Factors in analysis | Statistical Output |  |  |
| --- | --- | --- | --- |
| <u>RM-ANOVA (Region x Group)</u> | <u>F-value</u> | <u>P-value</u> | <u>Partial $\eta^2$</u> |
| Region (within-subjects) | F(1.813,45.313) = 18.664 | <0.001 | 0.427 |
| Group (between-subjects) | F(3,25) = 5.962 | 0.003 | 0.417 |
| Region x Group | F(5.438,45.313) = 7.981 | <0.001 | 0.489 |
| <u>1-way-ANOVA (Region)</u> | <u>F-value (ndf=3,ddf=25)</u> | <u>P-value</u> | <u>$\eta^2$</u> |
| Infralimbic area (ILA) | 10.835 | <0.001 | 0.565 |
| Visual areas (VIS) | 9.141 | <0.001 | 0.523 |
| Orbital area (ORB) | 12.209 | <0.001 | 0.594 |
| Anterior cingulate area (ACA) | 3.884 | 0.021 | 0.318 |
| Agranular insular area (AI) | 8.880 | <0.001 | 0.516 |
| Prelimbic area (PL) | 7.764 | <0.001 | 0.482 |
| Ectorhinal area (ECT) | 9.158 | <0.001 | 0.524 |
| Retrosplenial area (RSP) | 3.044 | 0.047 | 0.268 |
| Perirhinal area (PERI) | 7.569 | <0.001 | 0.476 |
| Somatomotor areas (MO) | 3.731 | 0.024 | 0.309 |
| Somatosensory areas (SS) | 3.419 | 0.033 | 0.291 |
| Temporal association areas (TEa) | 7.361 | 0.001 | 0.469 |
| Auditory areas (AUD) | 5.471 | 0.005 | 0.396 |
| Gustatory areas (GU) | 4.325 | 0.014 | 0.342 |
| Visceral area (VISC) | 2.905 | 0.055 | 0.259 |
| Frontal pole, cerebral cortex (FRP) | 0.574 | 0.638 | 0.064 |
| Dorsal peduncular area (DP) | 9.128 | <0.001 | 0.523 |
| Taenia tecta (TT) | 9.403 | <0.001 | 0.530 |
| Cortical amygdalar area (COA) | 4.597 | 0.011 | 0.355 |
| Piriform area (PIR) | 6.227 | 0.003 | 0.428 |
| Anterior olfactory nucleus (AON) | 11.283 | <0.001 | 0.575 |
| Piriform-amygdalar area (PAA) | 4.054 | 0.018 | 0.327 |
| Nucleus of the lateral olfactory tract (NLOT) | 3.233 | 0.039 | 0.280 |
| Postpiriform transition area (TR) | 3.296 | 0.037 | 0.283 |
| Accessory olfactory bulb (AOB) | 0.089 | 0.966 | 0.011 |
| Main olfactory bulb (MOB) | 0.271 | 0.846 | 0.031 |
| Retrohippocampal region (RHP) | 9.113 | <0.001 | 0.522 |
| Ammon's horn (CA) | 2.356 | 0.096 | 0.220 |
| Dentate gyrus (DG) | 2.470 | 0.085 | 0.229 |
| Fasciola cinerea (FC) | 1.094 | 0.370 | 0.116 |
| Induseum griseum (IG) | 1.124 | 0.358 | 0.119 |
| Clastrum (CLA) | 10.132 | <0.001 | 0.549 |
| Endopiriform nucleus (EP) | 7.998 | <0.001 | 0.490 |
| Basomedial amygdalar nucleus (BMA) | 4.967 | 0.008 | 0.373 |
| Basolateral amygdalar nucleus (BLA) | 5.178 | 0.006 | 0.383 |
| Posterior amygdalar nucleus (PA) | 4.254 | 0.015 | 0.338 |
| Lateral amygdalar nucleus (LA) | 2.122 | 0.123 | 0.203 |
| Lateral septal complex (LSX) | 5.580 | 0.005 | 0.401 |
| Striatum ventral region (STRv) | 3.698 | 0.025 | 0.307 |
| Striatum-like amygdalar nuclei (sAMY) | 3.146 | 0.043 | 0.274 |
| Striatum dorsal region (STRd) | 2.341 | 0.098 | 0.219 |
| Pallidum, caudal region (PALc) | 3.350 | 0.035 | 0.287 |
| Pallidum, medial region (PALm) | 2.585 | 0.076 | 0.237 |
| Pallidum, ventral region (PALv) | 2.382 | 0.093 | 0.222 |
| Pallidum, dorsal region (PALd) | 1.924 | 0.152 | 0.188 |
| Thalamus, polymodal association cortex related (DORpm) | 1.728 | 0.187 | 0.172 |
| Thalamus, sensory-motor cortex related (DORsm) | 1.236 | 0.318 | 0.129 |
| Periventricular region (PVR) | 2.619 | 0.073 | 0.239 |
| Periventricular zone (PVZ) | 1.849 | 0.164 | 0.182 |
| Hypothalamic medial zone (MEZ) | 2.273 | 0.105 | 0.214 |
| Hypothalamic lateral zone (LZ) | 2.033 | 0.135 | 0.196 |

|  |  |  |  |  |
| --- | --- | --- | --- | --- |
| Midbrain, sensory related (MBsen) | 2.101 | 0.126 | 0.201 |  |
| Midbrain, behavioral state related (MBsta) | 1.308 | 0.294 | 0.136 |  |
| Midbrain, motor related (MBmot) | 1.414 | 0.262 | 0.145 |  |
| Medulla (MY) | 4.341 | 0.014 | 0.343 |  |
| Pons (P) | 2.064 | 0.131 | 0.199 |  |
| <u>Tukey HSD</u> | <u>Comparison</u> | <u>Mean Difference</u> | <u>Std. Error</u> | <u>P-value</u> |
| Infralimbic area (ILA) | Home cage vs. Day 1 | -0.389 | 3.823 | 1.000 |
|  | Home cage vs. Day 15 | -0.507 | 3.686 | 0.999 |
|  | Homecage vs. Day 60 | -21.537 | 4.231 | <0.001 |
|  | Day 1 vs. Day 15 | -0.117 | 3.926 | 1.000 |
|  | Day 1 vs. Day 60 | -21.147 | 4.442 | <0.001 |
|  | Day 15 vs. Day 60 | -21.030 | 4.325 | <0.001 |
| Visual areas (VIS) | Home cage vs. Day 1 | -1.796 | 3.236 | 0.944 |
|  | Home cage vs. Day 15 | -1.781 | 3.120 | 0.940 |
|  | Homecage vs. Day 60 | -17.515 | 3.581 | <0.001 |
|  | Day 1 vs. Day 15 | 0.015 | 3.323 | 1.000 |
|  | Day 1 vs. Day 60 | -15.719 | 3.760 | 0.002 |
|  | Day 15 vs. Day 60 | -15.734 | 3.661 | 0.001 |
| Orbital area (ORB) | Home cage vs. Day 1 | -0.058 | 1.362 | 1.000 |
|  | Home cage vs. Day 15 | 0.290 | 1.313 | 0.996 |
|  | Homecage vs. Day 60 | -7.954 | 1.508 | <0.001 |
|  | Day 1 vs. Day 15 | 0.348 | 1.399 | 0.994 |
|  | Day 1 vs. Day 60 | -7.896 | 1.583 | <0.001 |
|  | Day 15 vs. Day 60 | -8.244 | 1.541 | <0.001 |
| Anterior cingulate area (ACA) | Home cage vs. Day 1 | 0.405 | 2.254 | 0.998 |
|  | Home cage vs. Day 15 | 0.195 | 2.173 | 1.000 |
|  | Homecage vs. Day 60 | -7.311 | 2.494 | 0.034 |
|  | Day 1 vs. Day 15 | -0.210 | 2.315 | 1.000 |
|  | Day 1 vs. Day 60 | -7.716 | 2.619 | 0.033 |
|  | Day 15 vs. Day 60 | -7.506 | 2.550 | 0.033 |
| Agranular insular area (AI) | Home cage vs. Day 1 | -0.102 | 1.216 | 1.000 |
|  | Home cage vs. Day 15 | -0.017 | 1.172 | 1.000 |
|  | Homecage vs. Day 60 | -6.155 | 1.345 | <0.001 |
|  | Day 1 vs. Day 15 | 0.085 | 1.248 | 1.000 |
|  | Day 1 vs. Day 60 | -6.052 | 1.412 | 0.001 |
|  | Day 15 vs. Day 60 | -6.138 | 1.375 | <0.001 |
| Prelimbic area (PL) | Home cage vs. Day 1 | 0.471 | 1.318 | 0.984 |
|  | Home cage vs. Day 15 | 0.231 | 1.271 | 0.998 |
|  | Homecage vs. Day 60 | -5.976 | 1.459 | 0.002 |
|  | Day 1 vs. Day 15 | -0.240 | 1.354 | 0.998 |
|  | Day 1 vs. Day 60 | -6.447 | 1.532 | 0.002 |
|  | Day 15 vs. Day 60 | -6.207 | 1.491 | 0.002 |
| Ectorhinal area (ECT) | Home cage vs. Day 1 | -0.108 | 0.946 | 0.999 |
|  | Home cage vs. Day 15 | 0.114 | 0.912 | 0.999 |
|  | Homecage vs. Day 60 | -4.827 | 1.047 | <0.001 |
|  | Day 1 vs. Day 15 | 0.223 | 0.972 | 0.996 |
|  | Day 1 vs. Day 60 | -4.719 | 1.099 | 0.001 |
|  | Day 15 vs. Day 60 | -4.942 | 1.070 | <0.001 |
| Retrosplenial area (RSP) | Home cage vs. Day 1 | 0.302 | 1.546 | 0.997 |
|  | Home cage vs. Day 15 | 0.000 | 1.491 | 1.000 |
|  | Homecage vs. Day 60 | -4.458 | 1.711 | 0.068 |
|  | Day 1 vs. Day 15 | -0.302 | 1.588 | 0.998 |
|  | Day 1 vs. Day 60 | -4.760 | 1.797 | 0.062 |
|  | Day 15 vs. Day 60 | -4.458 | 1.749 | 0.076 |
| Perirhinal area (PERI) | Home cage vs. Day 1 | 0.053 | 0.873 | 1.000 |
|  | Home cage vs. Day 15 | 0.038 | 0.842 | 1.000 |
|  | Homecage vs. Day 60 | -4.031 | 0.966 | 0.002 |
|  | Day 1 vs. Day 15 | -0.015 | 0.897 | 1.000 |
|  | Day 1 vs. Day 60 | -4.084 | 1.015 | 0.002 |
|  | Day 15 vs. Day 60 | -4.068 | 0.988 | 0.002 |
| Somatomotor areas (MO) | Home cage vs. Day 1 | 0.565 | 1.308 | 0.972 |
|  | Home cage vs. Day 15 | 0.695 | 1.261 | 0.945 |
|  | Homecage vs. Day 60 | -3.806 | 1.448 | 0.065 |
|  | Day 1 vs. Day 15 | 0.130 | 1.343 | 1.000 |
|  | Day 1 vs. Day 60 | -4.371 | 1.520 | 0.038 |
|  | Day 15 vs. Day 60 | -4.501 | 1.480 | 0.026 |
| Somatosensory areas (SS) | Home cage vs. Day 1 | 0.211 | 1.155 | 0.998 |
|  | Home cage vs. Day 15 | 0.436 | 1.114 | 0.979 |
|  | Homecage vs. Day 60 | -3.374 | 1.278 | 0.063 |
|  | Day 1 vs. Day 15 | 0.225 | 1.186 | 0.998 |
|  | Day 1 vs. Day 60 | -3.586 | 1.342 | 0.059 |
|  | Day 15 vs. Day 60 | -3.811 | 1.307 | 0.035 |
| Temporal association areas (TEa) | Home cage vs. Day 1 | 0.088 | 0.767 | 0.999 |
|  | Home cage vs. Day 15 | 0.356 | 0.740 | 0.963 |
|  | Homecage vs. Day 60 | -3.354 | 0.849 | 0.003 |
|  | Day 1 vs. Day 15 | 0.268 | 0.788 | 0.986 |
|  | Day 1 vs. Day 60 | -3.442 | 0.892 | 0.004 |
|  | Day 15 vs. Day 60 | -3.710 | 0.868 | 0.001 |
| Auditory areas (AUD) | Home cage vs. Day 1 | 0.051 | 0.846 | 1.000 |
|  | Home cage vs. Day 15 | 0.361 | 0.816 | 0.970 |
|  | Homecage vs. Day 60 | -3.186 | 0.936 | 0.011 |
|  | Day 1 vs. Day 15 | 0.310 | 0.869 | 0.984 |
|  | Day 1 vs. Day 60 | -3.237 | 0.983 | 0.015 |
|  | Day 15 vs. Day 60 | -3.547 | 0.957 | 0.005 |
| Gustatory areas (GU) | Home cage vs. Day 1 | 0.422 | 0.643 | 0.912 |
|  | Home cage vs. Day 15 | 0.541 | 0.620 | 0.818 |
|  | Homecage vs. Day 60 | -1.880 | 0.712 | 0.063 |
|  | Day 1 vs. Day 15 | 0.119 | 0.660 | 0.998 |
|  | Day 1 vs. Day 60 | -2.303 | 0.747 | 0.024 |
|  | Day 15 vs. Day 60 | -2.422 | 0.727 | 0.013 |
| Visceral area (VISC) | Home cage vs. Day 1 | 0.212 | 0.578 | 0.983 |
|  | Home cage vs. Day 15 | 0.329 | 0.557 | 0.934 |
|  | Homecage vs. Day 60 | -1.458 | 0.640 | 0.130 |
|  | Day 1 vs. Day 15 | 0.117 | 0.594 | 0.997 |
|  | Day 1 vs. Day 60 | -1.671 | 0.672 | 0.087 |
|  | Day 15 vs. Day 60 | -1.787 | 0.654 | 0.052 |
| Frontal pole, cerebral cortex (FRP) | Home cage vs. Day 1 | 0.037 | 0.461 | 1.000 |
|  | Home cage vs. Day 15 | -0.058 | 0.445 | 0.999 |
|  | Homecage vs. Day 60 | 0.574 | 0.510 | 0.678 |
|  | Day 1 vs. Day 15 | -0.095 | 0.474 | 0.997 |
|  | Day 1 vs. Day 60 | 0.537 | 0.536 | 0.749 |
|  | Day 15 vs. Day 60 | 0.633 | 0.522 | 0.625 |
| Dorsal peduncular area (DP) | Home cage vs. Day 1 | -0.922 | 3.029 | 0.990 |
|  | Home cage vs. Day 15 | -0.511 | 2.920 | 0.998 |
|  | Homecage vs. Day 60 | -15.873 | 3.352 | <0.001 |
|  | Day 1 vs. Day 15 | 0.412 | 3.110 | 0.999 |
|  | Day 1 vs. Day 60 | -14.950 | 3.519 | 0.001 |
|  | Day 15 vs. Day 60 | -15.362 | 3.426 | <0.001 |
| Taenia tecta (TT) | Home cage vs. Day 1 | -0.156 | 1.645 | 1.000 |
|  | Home cage vs. Day 15 | -0.524 | 1.586 | 0.987 |
|  | Homecage vs. Day 60 | -8.727 | 1.821 | <0.001 |
|  | Day 1 vs. Day 15 | -0.368 | 1.690 | 0.996 |
|  | Day 1 vs. Day 60 | -8.571 | 1.912 | <0.001 |
|  | Day 15 vs. Day 60 | -8.203 | 1.861 | <0.001 |
| Cortical amygdalar area (COA) | Home cage vs. Day 1 | -0.463 | 1.844 | 0.994 |
|  | Home cage vs. Day 15 | -1.448 | 1.778 | 0.847 |
|  | Homecage vs. Day 60 | -7.131 | 2.041 | 0.009 |
|  | Day 1 vs. Day 15 | -0.985 | 1.894 | 0.953 |
|  | Day 1 vs. Day 60 | -6.668 | 2.143 | 0.022 |
|  | Day 15 vs. Day 60 | -5.683 | 2.086 | 0.053 |
| Piriform area (PIR) | Home cage vs. Day 1 | -0.240 | 1.305 | 0.998 |
|  | Home cage vs. Day 15 | -0.574 | 1.259 | 0.968 |
|  | Homecage vs. Day 60 | -5.734 | 1.445 | 0.003 |
|  | Day 1 vs. Day 15 | -0.334 | 1.341 | 0.994 |
|  | Day 1 vs. Day 60 | -5.494 | 1.517 | 0.007 |
|  | Day 15 vs. Day 60 | -5.160 | 1.477 | 0.009 |
| Anterior olfactory nucleus (AON) | Home cage vs. Day 1 | -0.095 | 0.860 | 0.999 |
|  | Home cage vs. Day 15 | -0.280 | 0.829 | 0.986 |
|  | Homecage vs. Day 60 | -4.991 | 0.951 | <0.001 |
|  | Day 1 vs. Day 15 | -0.185 | 0.883 | 0.997 |
|  | Day 1 vs. Day 60 | -4.896 | 0.999 | <0.001 |
|  | Day 15 vs. Day 60 | -4.711 | 0.972 | <0.001 |
| Piriform-amygdalar area (PAA) | Home cage vs. Day 1 | -0.233 | 1.280 | 0.998 |
|  | Home cage vs. Day 15 | -1.101 | 1.234 | 0.809 |
|  | Homecage vs. Day 60 | -4.634 | 1.417 | 0.015 |
|  | Day 1 vs. Day 15 | -0.868 | 1.315 | 0.911 |
|  | Day 1 vs. Day 60 | -4.401 | 1.487 | 0.032 |
|  | Day 15 vs. Day 60 | -3.533 | 1.448 | 0.095 |
| Nucleus of the lateral olfactory tract (NLOT) | Home cage vs. Day 1 | 0.006 | 1.259 | 1.000 |
|  | Home cage vs. Day 15 | -0.120 | 1.214 | 1.000 |
|  | Homecage vs. Day 60 | -3.862 | 1.394 | 0.048 |
|  | Day 1 vs. Day 15 | -0.126 | 1.293 | 1.000 |
|  | Day 1 vs. Day 60 | -3.868 | 1.463 | 0.063 |
|  | Day 15 vs. Day 60 | -3.742 | 1.425 | 0.065 |
| Postpiriform transition area (TR) | Home cage vs. Day 1 | 0.304 | 0.613 | 0.959 |
|  | Home cage vs. Day 15 | 0.195 | 0.591 | 0.987 |
|  | Homecage vs. Day 60 | -1.702 | 0.679 | 0.083 |
|  | Day 1 vs. Day 15 | -0.109 | 0.630 | 0.998 |
|  | Day 1 vs. Day 60 | -2.006 | 0.712 | 0.043 |
|  | Day 15 vs. Day 60 | -1.897 | 0.694 | 0.052 |
| Accessory olfactory bulb (AOB) | Home cage vs. Day 1 | 0.027 | 0.588 | 1.000 |
|  | Home cage vs. Day 15 | 0.260 | 0.567 | 0.967 |
|  | Homecage vs. Day 60 | 0.010 | 0.651 | 1.000 |
|  | Day 1 vs. Day 15 | 0.233 | 0.604 | 0.980 |
|  | Day 1 vs. Day 60 | -0.017 | 0.683 | 1.000 |
|  | Day 15 vs. Day 60 | -0.250 | 0.665 | 0.982 |
| Main olfactory bulb (MOB) | Home cage vs. Day 1 | 0.215 | 0.560 | 0.980 |
|  | Home cage vs. Day 15 | 0.049 | 0.540 | 1.000 |
|  | Homecage vs. Day 60 | 0.522 | 0.620 | 0.834 |
|  | Day 1 vs. Day 15 | -0.167 | 0.575 | 0.991 |
|  | Day 1 vs. Day 60 | 0.306 | 0.651 | 0.965 |
|  | Day 15 vs. Day 60 | 0.473 | 0.634 | 0.877 |
| Retrohippocampal region (RHP) | Home cage vs. Day 1 | 0.146 | 1.372 | 1.000 |
|  | Home cage vs. Day 15 | -0.129 | 1.323 | 1.000 |
|  | Homecage vs. Day 60 | -6.992 | 1.518 | <0.001 |
|  | Day 1 vs. Day 15 | -0.274 | 1.409 | 0.997 |
|  | Day 1 vs. Day 60 | -7.138 | 1.594 | <0.001 |
|  | Day 15 vs. Day 60 | -6.863 | 1.552 | <0.001 |
| Ammon's horn (CA) | Home cage vs. Day 1 | 0.449 | 0.753 | 0.932 |
|  | Home cage vs. Day 15 | 0.400 | 0.726 | 0.945 |
|  | Homecage vs. Day 60 | -1.625 | 0.834 | 0.234 |
|  | Day 1 vs. Day 15 | -0.049 | 0.774 | 1.000 |
|  | Day 1 vs. Day 60 | -2.075 | 0.875 | 0.109 |
|  | Day 15 vs. Day 60 | -2.025 | 0.852 | 0.108 |
| Dentate gyrus (DG) | Home cage vs. Day 1 | 0.914 | 0.658 | 0.518 |
|  | Home cage vs. Day 15 | 0.589 | 0.635 | 0.790 |
|  | Homecage vs. Day 60 | -1.025 | 0.729 | 0.507 |
|  | Day 1 vs. Day 15 | -0.325 | 0.676 | 0.963 |
|  | Day 1 vs. Day 60 | -1.939 | 0.765 | 0.079 |
|  | Day 15 vs. Day 60 | -1.614 | 0.745 | 0.160 |
| Fasciola cinerea (FC) | Home cage vs. Day 1 | 0.424 | 0.713 | 0.933 |
|  | Home cage vs. Day 15 | 0.814 | 0.688 | 0.643 |
|  | Homecage vs. Day 60 | -0.558 | 0.789 | 0.893 |
|  | Day 1 vs. Day 15 | 0.390 | 0.732 | 0.950 |
|  | Day 1 vs. Day 60 | -0.982 | 0.829 | 0.641 |
|  | Day 15 vs. Day 60 | -1.372 | 0.807 | 0.344 |
| Induseum griseum (IG) | Home cage vs. Day 1 | 0.425 | 0.313 | 0.537 |
|  | Home cage vs. Day 15 | 0.475 | 0.302 | 0.412 |
|  | Homecage vs. Day 60 | 0.095 | 0.347 | 0.993 |
|  | Day 1 vs. Day 15 | 0.050 | 0.322 | 0.999 |
|  | Day 1 vs. Day 60 | -0.331 | 0.364 | 0.801 |
|  | Day 15 vs. Day 60 | -0.381 | 0.354 | 0.708 |
| Clastrum (CLA) | Home cage vs. Day 1 | -4.323 | 4.982 | 0.821 |
|  | Home cage vs. Day 15 | -6.156 | 4.804 | 0.583 |
|  | Homecage vs. Day 60 | -29.328 | 5.515 | <0.001 |
|  | Day 1 vs. Day 15 | -1.833 | 5.117 | 0.984 |
|  | Day 1 vs. Day 60 | -25.006 | 5.789 | 0.001 |
|  | Day 15 vs. Day 60 | -23.173 | 5.636 | 0.002 |
| Endopiriform nucleus (EP) | Home cage vs. Day 1 | -1.798 | 2.489 | 0.887 |
|  | Home cage vs. Day 15 | -2.106 | 2.399 | 0.816 |
|  | Homecage vs. Day 60 | -12.891 | 2.754 | <0.001 |
|  | Day 1 vs. Day 15 | -0.308 | 2.556 | 0.999 |
|  | Day 1 vs. Day 60 | -11.093 | 2.891 | 0.004 |
|  | Day 15 vs. Day 60 | -10.785 | 2.815 | 0.004 |
| Basomedial amygdalar nucleus (BMA) | Home cage vs. Day 1 | -1.148 | 2.059 | 0.944 |
|  | Home cage vs. Day 15 | -1.584 | 1.986 | 0.855 |
|  | Homecage vs. Day 60 | -8.437 | 2.279 | 0.005 |
|  | Day 1 vs. Day 15 | -0.436 | 2.115 | 0.997 |
|  | Day 1 vs. Day 60 | -7.289 | 2.393 | 0.026 |
|  | Day 15 vs. Day 60 | -6.853 | 2.330 | 0.033 |
| Basolateral amygdalar nucleus (BLA) | Home cage vs. Day 1 | -0.778 | 1.181 | 0.911 |
|  | Home cage vs. Day 15 | -0.924 | 1.139 | 0.849 |
|  | Homecage vs. Day 60 | -4.961 | 1.307 | 0.004 |
|  | Day 1 vs. Day 15 | -0.146 | 1.213 | 0.999 |
|  | Day 1 vs. Day 60 | -4.183 | 1.372 | 0.026 |
|  | Day 15 vs. Day 60 | -4.038 | 1.336 | 0.027 |
| Posterior amygdalar nucleus (PA) | Home cage vs. Day 1 | 0.103 | 1.353 | 1.000 |
|  | Home cage vs. Day 15 | -1.104 | 1.304 | 0.832 |
|  | Homecage vs. Day 60 | -4.866 | 1.497 | 0.016 |
|  | Day 1 vs. Day 15 | -1.207 | 1.389 | 0.821 |
|  | Day 1 vs. Day 60 | -4.969 | 1.572 | 0.020 |
|  | Day 15 vs. Day 60 | -3.762 | 1.530 | 0.092 |
| Lateral amygdalar nucleus (LA) | Home cage vs. Day 1 | -0.428 | 0.678 | 0.921 |
|  | Home cage vs. Day 15 | 0.302 | 0.653 | 0.966 |
|  | Homecage vs. Day 60 | -1.540 | 0.750 | 0.196 |
|  | Day 1 vs. Day 15 | 0.731 | 0.696 | 0.722 |
|  | Day 1 vs. Day 60 | -1.112 | 0.787 | 0.504 |
|  | Day 15 vs. Day 60 | -1.842 | 0.767 | 0.102 |
| Lateral septal complex (LSX) | Home cage vs. Day 1 | -0.136 | 2.983 | 1.000 |
|  | Home cage vs. Day 15 | -0.490 | 2.877 | 0.998 |
|  | Homecage vs. Day 60 | -12.099 | 3.302 | 0.006 |
|  | Day 1 vs. Day 15 | -0.353 | 3.064 | 0.999 |
|  | Day 1 vs. Day 60 | -11.963 | 3.466 | 0.010 |
|  | Day 15 vs. Day 60 | -11.610 | 3.375 | 0.010 |
| Striatum ventral region (STRv) | Home cage vs. Day 1 | 0.246 | 1.452 | 0.998 |
|  | Home cage vs. Day 15 | 0.265 | 1.400 | 0.998 |
|  | Homecage vs. Day 60 | -4.546 | 1.607 | 0.042 |
|  | Day 1 vs. Day 15 | 0.019 | 1.491 | 1.000 |
|  | Day 1 vs. Day 60 | -4.793 | 1.687 | 0.041 |
|  | Day 15 vs. Day 60 | -4.811 | 1.642 | 0.034 |
| Striatum-like amygdalar nuclei (sAMY) | Home cage vs. Day 1 | -0.193 | 1.447 | 0.999 |
|  | Home cage vs. Day 15 | -0.558 | 1.395 | 0.978 |
|  | Homecage vs. Day 60 | -4.541 | 1.601 | 0.042 |
|  | Day 1 vs. Day 15 | -0.365 | 1.486 | 0.995 |
|  | Day 1 vs. Day 60 | -4.347 | 1.681 | 0.071 |
|  | Day 15 vs. Day 60 | -3.983 | 1.637 | 0.096 |
| Striatum dorsal region (STRd) | Home cage vs. Day 1 | 0.457 | 0.566 | 0.851 |
|  | Home cage vs. Day 15 | 0.757 | 0.546 | 0.519 |
|  | Homecage vs. Day 60 | -0.856 | 0.626 | 0.531 |
|  | Day 1 vs. Day 15 | 0.300 | 0.581 | 0.954 |
|  | Day 1 vs. Day 60 | -1.312 | 0.657 | 0.216 |
|  | Day 15 vs. Day 60 | -1.613 | 0.640 | 0.081 |
| Pallidum, caudal region (PALc) | Home cage vs. Day 1 | -0.198 | 2.328 | 1.000 |
|  | Home cage vs. Day 15 | -0.487 | 2.245 | 0.996 |
|  | Homecage vs. Day 60 | -7.402 | 2.577 | 0.038 |
|  | Day 1 vs. Day 15 | -0.289 | 2.391 | 0.999 |
|  | Day 1 vs. Day 60 | -7.204 | 2.705 | 0.060 |
|  | Day 15 vs. Day 60 | -6.915 | 2.634 | 0.065 |
| Pallidum, medial region (PALm) | Home cage vs. Day 1 | 0.720 | 1.136 | 0.920 |
|  | Home cage vs. Day 15 | 0.450 | 1.095 | 0.976 |
|  | Homecage vs. Day 60 | -2.640 | 1.257 | 0.180 |
|  | Day 1 vs. Day 15 | -0.270 | 1.166 | 0.995 |
|  | Day 1 vs. Day 60 | -3.360 | 1.319 | 0.077 |
|  | Day 15 vs. Day 60 | -3.090 | 1.285 | 0.102 |
| Pallidum, ventral region (PALv) | Home cage vs. Day 1 | 0.656 | 0.796 | 0.842 |
|  | Home cage vs. Day 15 | 0.755 | 0.767 | 0.760 |
|  | Homecage vs. Day 60 | -1.458 | 0.881 | 0.367 |
|  | Day 1 vs. Day 15 | 0.099 | 0.817 | 0.999 |
|  | Day 1 vs. Day 60 | -2.114 | 0.924 | 0.128 |
|  | Day 15 vs. Day 60 | -2.213 | 0.900 | 0.092 |
| Pallidum, dorsal region (PALd) | Home cage vs. Day 1 | 0.683 | 0.453 | 0.449 |
|  | Home cage vs. Day 15 | 0.754 | 0.437 | 0.332 |
|  | Homecage vs. Day 60 | -0.195 | 0.502 | 0.980 |
|  | Day 1 vs. Day 15 | 0.071 | 0.465 | 0.999 |
|  | Day 1 vs. Day 60 | -0.878 | 0.526 | 0.361 |
|  | Day 15 vs. Day 60 | -0.949 | 0.513 | 0.274 |
| Thalamus, polymodal association cortex related (DORpm) | Home cage vs. Day 1 | 0.575 | 0.405 | 0.500 |
|  | Home cage vs. Day 15 | 0.651 | 0.390 | 0.360 |
|  | Homecage vs. Day 60 | -0.152 | 0.448 | 0.986 |
|  | Day 1 vs. Day 15 | 0.077 | 0.416 | 0.998 |
|  | Day 1 vs. Day 60 | -0.727 | 0.470 | 0.427 |
|  | Day 15 vs. Day 60 | -0.804 | 0.458 | 0.318 |
| Thalamus, sensory-motor cortex related (DORsm) | Home cage vs. Day 1 | 0.479 | 0.334 | 0.491 |
|  | Home cage vs. Day 15 | 0.494 | 0.323 | 0.435 |
|  | Homecage vs. Day 60 | 0.036 | 0.370 | 1.000 |
|  | Day 1 vs. Day 15 | 0.015 | 0.343 | 1.000 |
|  | Day 1 vs. Day 60 | -0.443 | 0.389 | 0.669 |
|  | Day 15 vs. Day 60 | -0.458 | 0.378 | 0.627 |
| Periventricular region (PVR) | Home cage vs. Day 1 | -0.035 | 1.965 | 1.000 |
|  | Home cage vs. Day 15 | -0.922 | 1.895 | 0.961 |
|  | Homecage vs. Day 60 | -5.591 | 2.175 | 0.073 |
|  | Day 1 vs. Day 15 | -0.887 | 2.018 | 0.971 |
|  | Day 1 vs. Day 60 | -5.556 | 2.283 | 0.096 |
|  | Day 15 vs. Day 60 | -4.669 | 2.223 | 0.181 |
| Periventricular zone (PVZ) | Home cage vs. Day 1 | 0.386 | 1.838 | 0.997 |
|  | Home cage vs. Day 15 | -1.274 | 1.772 | 0.889 |
|  | Homecage vs. Day 60 | -4.186 | 2.034 | 0.195 |
|  | Day 1 vs. Day 15 | -1.660 | 1.888 | 0.816 |
|  | Day 1 vs. Day 60 | -4.571 | 2.136 | 0.168 |
|  | Day 15 vs. Day 60 | -2.912 | 2.079 | 0.511 |
| Hypothalamic medial zone (MEZ) | Home cage vs. Day 1 | 0.318 | 1.568 | 0.997 |
|  | Home cage vs. Day 15 | -0.477 | 1.512 | 0.989 |
|  | Homecage vs. Day 60 | -3.987 | 1.736 | 0.126 |
|  | Day 1 vs. Day 15 | -0.796 | 1.610 | 0.960 |
|  | Day 1 vs. Day 60 | -4.305 | 1.822 | 0.111 |
|  | Day 15 vs. Day 60 | -3.509 | 1.774 | 0.223 |
| Hypothalamic lateral zone (LZ) | Home cage vs. Day 1 | 0.279 | 1.176 | 0.995 |
|  | Home cage vs. Day 15 | -0.058 | 1.134 | 1.000 |
|  | Homecage vs. Day 60 | -2.751 | 1.302 | 0.177 |
|  | Day 1 vs. Day 15 | -0.337 | 1.208 | 0.992 |
|  | Day 1 vs. Day 60 | -3.030 | 1.367 | 0.146 |
|  | Day 15 vs. Day 60 | -2.694 | 1.331 | 0.206 |
| Midbrain, sensory related (MBsen) | Home cage vs. Day 1 | 0.542 | 0.575 | 0.782 |
|  | Home cage vs. Day 15 | 0.508 | 0.554 | 0.796 |
|  | Homecage vs. Day 60 | -0.939 | 0.636 | 0.466 |
|  | Day 1 vs. Day 15 | -0.034 | 0.590 | 1.000 |
|  | Day 1 vs. Day 60 | -1.482 | 0.668 | 0.146 |
|  | Day 15 vs. Day 60 | -1.448 | 0.650 | 0.143 |
| Midbrain, behavioral state related (MBsta) | Home cage vs. Day 1 | 0.756 | 0.750 | 0.746 |
|  | Home cage vs. Day 15 | 0.467 | 0.723 | 0.916 |
|  | Homecage vs. Day 60 | -0.862 | 0.830 | 0.729 |
|  | Day 1 vs. Day 15 | -0.289 | 0.770 | 0.982 |
|  | Day 1 vs. Day 60 | -1.618 | 0.872 | 0.272 |
|  | Day 15 vs. Day 60 | -1.329 | 0.849 | 0.415 |
| Midbrain, motor related (MBmot) | Home cage vs. Day 1 | 0.640 | 0.588 | 0.700 |
|  | Home cage vs. Day 15 | 0.590 | 0.567 | 0.728 |
|  | Homecage vs. Day 60 | -0.557 | 0.651 | 0.828 |
|  | Day 1 vs. Day 15 | -0.050 | 0.604 | 1.000 |
|  | Day 1 vs. Day 60 | -1.197 | 0.684 | 0.320 |
|  | Day 15 vs. Day 60 | -1.147 | 0.666 | 0.334 |
| Medulla (MY) | Home cage vs. Day 1 | -0.203 | 2.017 | 1.000 |
|  | Home cage vs. Day 15 | -1.732 | 1.945 | 0.810 |
|  | Homecage vs. Day 60 | -7.485 | 2.233 | 0.013 |
|  | Day 1 vs. Day 15 | -1.528 | 2.072 | 0.881 |
|  | Day 1 vs. Day 60 | -7.282 | 2.344 | 0.023 |
|  | Day 15 vs. Day 60 | -5.753 | 2.282 | 0.081 |
| Pons (P) | Home cage vs. Day 1 | 0.692 | 1.280 | 0.948 |
|  | Home cage vs. Day 15 | 0.158 | 1.234 | 0.999 |
|  | Homecage vs. Day 60 | -2.773 | 1.416 | 0.231 |
|  | Day 1 vs. Day 15 | -0.534 | 1.314 | 0.977 |
|  | Day 1 vs. Day 60 | -3.465 | 1.487 | 0.118 |
|  | Day 15 vs. Day 60 | -2.931 | 1.447 | 0.206 |

**Table S6.** Statistical output for Figure 3B, right panel : Analysis of Z-scored counts from 5 major anatomical regions within AP +1.55 to AP +1.75 coronal subvolume (n=31, analyses shown in figure 3B are highlighted in grey)

| Factors in analysis | Statistical output |  |  |  |
| --- | --- | --- | --- | --- |
| <u>RM-ANOVA (Region x Group)</u> | <u>F-value</u> | <u>P-value</u> | <u>Partial $\eta^2$</u> |  |
| Region (within-subjects) | F(1.573,4.720) = 6.533 | 0.006 | 0.189 |  |
| Group (between-subjects) | F(3,28) = 6.853 | 0.001 | 0.423 |  |
| Region x Group | F(4.720,44.049) = 4.689 | 0.002 | 0.334 |  |
| <u>1-way-ANOVA (Region)</u> | <u>F-value (ndf=3,ddf=28)</u> | <u>P-value</u> | <u>$\eta^2$</u> |  |
| Isocortex | 7.388 | 0.001 | 0.442 |  |
| Olfactory areas | 8.949 | <0.001 | 0.489 |  |
| Cortical subplate | 7.689 | <0.001 | 0.452 |  |
| Striatum | 5.953 | 0.003 | 0.389 |  |
| Pallidum | 4.790 | 0.008 | 0.339 |  |
| <u>Tukey HSD</u> | <u>Comparison</u> | <u>Mean Difference</u> | <u>Std. Error</u> | <u>P-value</u> |
| Isocortex | Home cage vs. Day 1 | 0.545 | 1.778 | 0.990 |
|  | Home cage vs. Day 15 | 0.031 | 1.778 | 1.000 |
|  | Homecage vs. Day 60 | -8.578 | 2.064 | 0.001 |
|  | Day 1 vs. Day 15 | -0.514 | 1.913 | 0.993 |
|  | Day 1 vs. Day 60 | -9.123 | 2.181 | 0.001 |
|  | Day 15 vs. Day 60 | -8.609 | 2.181 | 0.003 |
| Olfactory areas | Home cage vs. Day 1 | 0.595 | 1.703 | 0.985 |
|  | Home cage vs. Day 15 | -0.957 | 1.703 | 0.942 |
|  | Homecage vs. Day 60 | -9.228 | 1.977 | 0.000 |
|  | Day 1 vs. Day 15 | -1.552 | 1.833 | 0.832 |
|  | Day 1 vs. Day 60 | -9.822 | 2.090 | 0.000 |
|  | Day 15 vs. Day 60 | -8.271 | 2.090 | 0.003 |
| Cortical subplate | Home cage vs. Day 1 | 0.459 | 2.752 | 0.998 |
|  | Home cage vs. Day 15 | -0.700 | 2.752 | 0.994 |
|  | Homecage vs. Day 60 | -13.872 | 3.194 | 0.001 |
|  | Day 1 vs. Day 15 | -1.159 | 2.961 | 0.979 |
|  | Day 1 vs. Day 60 | -14.331 | 3.376 | 0.001 |
|  | Day 15 vs. Day 60 | -13.172 | 3.376 | 0.003 |
| Striatum | Home cage vs. Day 1 | 0.754 | 2.972 | 0.994 |
|  | Home cage vs. Day 15 | -0.941 | 2.972 | 0.989 |
|  | Homecage vs. Day 60 | -13.111 | 3.450 | 0.004 |
|  | Day 1 vs. Day 15 | -1.696 | 3.198 | 0.951 |
|  | Day 1 vs. Day 60 | -13.866 | 3.646 | 0.004 |
|  | Day 15 vs. Day 60 | -12.170 | 3.646 | 0.012 |
| Pallidum | Home cage vs. Day 1 | 0.750 | 2.204 | 0.986 |
|  | Home cage vs. Day 15 | -1.217 | 2.204 | 0.945 |
|  | Homecage vs. Day 60 | -8.677 | 2.559 | 0.011 |
|  | Day 1 vs. Day 15 | -1.966 | 2.372 | 0.840 |
|  | Day 1 vs. Day 60 | -9.427 | 2.704 | 0.008 |
|  | Day 15 vs. Day 60 | -7.460 | 2.704 | 0.047 |

**Table S7.** Statistical output for Figure 3C: Analysis of Z-scored counts from 18 subregions within AP +1.55 to AP +1.75 coronal subvolume (n=31)

| Factors in analysis | Statistical Output |  |  |  |
| --- | --- | --- | --- | --- |
| <u>RM-ANOVA (Region x Group)</u> | <u>F-value</u> | <u>P-value</u> | <u>Partial $\eta^2$</u> |  |
| Region (within-subjects) | F(1.168,32.698) = 9.502 | 0.003 | 0.253 |  |
| Group (between-subjects) | F(3,28) = 7.240 | 0.001 | 0.437 |  |
| Region x Group | F(3.503,32.698) = 6.321 | 0.001 | 0.404 |  |
| <u>1-way-ANOVA (Region)</u> | <u>F-value (ndf=3,ddf=28)</u> | <u>P-value</u> | <u>$\eta^2$</u> |  |
| Orbital area (ORB) | 11.349 | <0.001 | 0.549 |  |
| Anterior cingulate area (ACA) | 6.074 | 0.003 | 0.394 |  |
| Infralimbic area (ILA) | 7.624 | <0.001 | 0.450 |  |
| Prelimbic area (PL) | 6.868 | 0.001 | 0.424 |  |
| Agranular insular area (AI) | 11.030 | <0.001 | 0.542 |  |
| Somatomotor areas (MO) | 5.737 | 0.003 | 0.381 |  |
| Gustatory areas (GU) | 4.764 | 0.008 | 0.338 |  |
| Somatosensory areas (SS) | 4.215 | 0.014 | 0.311 |  |
| Dorsal peduncular area (DP) | 5.876 | 0.003 | 0.386 |  |
| Taenia tecta (TT) | 7.782 | <0.001 | 0.455 |  |
| Anterior olfactory nucleus (AON) | 10.154 | <0.001 | 0.521 |  |
| Piriform area (PIR) | 8.909 | <0.001 | 0.488 |  |
| Endopiriform nucleus (EP) | 8.681 | <0.001 | 0.482 |  |
| Clastrum (CLA) | 6.220 | 0.002 | 0.400 |  |
| Lateral septum (LS) | 5.714 | 0.004 | 0.380 |  |
| Nucleus accumbens (ACB) | 5.509 | 0.004 | 0.371 |  |
| Olfactory tubercle (OT) | 6.070 | 0.003 | 0.394 |  |
| Substantia innominata (SI) | 4.790 | 0.008 | 0.339 |  |
| <u>Tukey HSD</u> | <u>Comparison</u> | <u>Mean Difference</u> | <u>Std. Error</u> | <u>P-value</u> |
| Orbital area (ORB) | Home cage vs. Day 1 | 0.018 | 4.269 | 1.000 |
|  | Home cage vs. Day 15 | -1.379 | 4.269 | 0.988 |
|  | Homecage vs. Day 60 | -26.455 | 4.956 | <0.001 |
|  | Day 1 vs. Day 15 | -1.397 | 4.594 | 0.990 |
|  | Day 1 vs. Day 60 | -26.473 | 5.238 | <0.001 |
|  | Day 15 vs. Day 60 | -25.076 | 5.238 | <0.001 |
| Anterior cingulate area (ACA) | Home cage vs. Day 1 | 0.885 | 5.763 | 0.999 |
|  | Home cage vs. Day 15 | -1.098 | 5.763 | 0.997 |
|  | Homecage vs. Day 60 | -25.764 | 6.689 | 0.003 |
|  | Day 1 vs. Day 15 | -1.982 | 6.201 | 0.988 |
|  | Day 1 vs. Day 60 | -26.648 | 7.070 | 0.004 |
|  | Day 15 vs. Day 60 | -24.666 | 7.070 | 0.008 |
| Infralimbic area (ILA) | Home cage vs. Day 1 | 0.147 | 4.793 | 1.000 |
|  | Home cage vs. Day 15 | -0.367 | 4.793 | 1.000 |
|  | Homecage vs. Day 60 | -24.079 | 5.564 | <0.001 |
|  | Day 1 vs. Day 15 | -0.514 | 5.158 | 1.000 |
|  | Day 1 vs. Day 60 | -24.226 | 5.881 | 0.002 |
|  | Day 15 vs. Day 60 | -23.713 | 5.881 | 0.002 |
| Prelimbic area (PL) | Home cage vs. Day 1 | 0.578 | 4.348 | 0.999 |
|  | Home cage vs. Day 15 | -1.044 | 4.348 | 0.995 |
|  | Homecage vs. Day 60 | -20.753 | 5.047 | 0.002 |
|  | Day 1 vs. Day 15 | -1.623 | 4.678 | 0.985 |
|  | Day 1 vs. Day 60 | -21.332 | 5.334 | 0.002 |
|  | Day 15 vs. Day 60 | -19.709 | 5.334 | 0.005 |
| Agranular insular area (AI) | Home cage vs. Day 1 | 0.349 | 1.931 | 0.998 |
|  | Home cage vs. Day 15 | -0.879 | 1.931 | 0.968 |
|  | Homecage vs. Day 60 | -11.727 | 2.241 | <0.001 |
|  | Day 1 vs. Day 15 | -1.228 | 2.077 | 0.934 |
|  | Day 1 vs. Day 60 | -12.076 | 2.369 | <0.001 |
|  | Day 15 vs. Day 60 | -10.848 | 2.369 | <0.001 |
| Somatomotor areas (MO) | Home cage vs. Day 1 | 0.662 | 1.516 | 0.972 |
|  | Home cage vs. Day 15 | 0.255 | 1.516 | 0.998 |
|  | Homecage vs. Day 60 | -6.283 | 1.760 | 0.007 |
|  | Day 1 vs. Day 15 | -0.406 | 1.632 | 0.994 |
|  | Day 1 vs. Day 60 | -6.945 | 1.860 | 0.004 |
|  | Day 15 vs. Day 60 | -6.539 | 1.860 | 0.008 |
| Gustatory areas (GU) | Home cage vs. Day 1 | 0.481 | 0.908 | 0.951 |
|  | Home cage vs. Day 15 | 0.201 | 0.908 | 0.996 |
|  | Homecage vs. Day 60 | -3.361 | 1.054 | 0.017 |
|  | Day 1 vs. Day 15 | -0.281 | 0.977 | 0.992 |
|  | Day 1 vs. Day 60 | -3.842 | 1.115 | 0.009 |
|  | Day 15 vs. Day 60 | -3.561 | 1.115 | 0.017 |
| Somatosensory areas (SS) | Home cage vs. Day 1 | 0.503 | 0.744 | 0.905 |
|  | Home cage vs. Day 15 | 0.587 | 0.744 | 0.859 |
|  | Homecage vs. Day 60 | -2.363 | 0.863 | 0.049 |
|  | Day 1 vs. Day 15 | 0.084 | 0.800 | 1.000 |
|  | Day 1 vs. Day 60 | -2.866 | 0.912 | 0.019 |
|  | Day 15 vs. Day 60 | -2.950 | 0.912 | 0.016 |
| Dorsal peduncular area (DP) | Home cage vs. Day 1 | 0.309 | 6.916 | 1.000 |
|  | Home cage vs. Day 15 | -1.005 | 6.916 | 0.999 |
|  | Homecage vs. Day 60 | -30.601 | 8.027 | 0.004 |
|  | Day 1 vs. Day 15 | -1.314 | 7.442 | 0.998 |
|  | Day 1 vs. Day 60 | -30.910 | 8.485 | 0.006 |
|  | Day 15 vs. Day 60 | -29.596 | 8.485 | 0.008 |
| Taenia tecta (TT) | Home cage vs. Day 1 | 0.297 | 4.496 | 1.000 |
|  | Home cage vs. Day 15 | -1.455 | 4.496 | 0.988 |
|  | Homecage vs. Day 60 | -23.030 | 5.219 | <0.001 |
|  | Day 1 vs. Day 15 | -1.752 | 4.838 | 0.983 |
|  | Day 1 vs. Day 60 | -23.327 | 5.516 | 0.001 |
|  | Day 15 vs. Day 60 | -21.575 | 5.516 | 0.003 |
| Anterior olfactory nucleus (AON) | Home cage vs. Day 1 | 0.447 | 2.981 | 0.999 |
|  | Home cage vs. Day 15 | -1.445 | 2.981 | 0.962 |
|  | Homecage vs. Day 60 | -17.425 | 3.460 | <0.001 |
|  | Day 1 vs. Day 15 | -1.892 | 3.208 | 0.934 |
|  | Day 1 vs. Day 60 | -17.872 | 3.657 | <0.001 |
|  | Day 15 vs. Day 60 | -15.980 | 3.657 | <0.001 |
| Piriform area (PIR) | Home cage vs. Day 1 | 0.608 | 1.229 | 0.960 |
|  | Home cage vs. Day 15 | -0.835 | 1.229 | 0.904 |
|  | Homecage vs. Day 60 | -6.574 | 1.427 | <0.001 |
|  | Day 1 vs. Day 15 | -1.442 | 1.323 | 0.698 |
|  | Day 1 vs. Day 60 | -7.182 | 1.508 | <0.001 |
|  | Day 15 vs. Day 60 | -5.739 | 1.508 | 0.004 |
| Endopiriform nucleus (EP) | Home cage vs. Day 1 | 0.502 | 1.817 | 0.992 |
|  | Home cage vs. Day 15 | -1.348 | 1.817 | 0.879 |
|  | Homecage vs. Day 60 | -9.782 | 2.109 | <0.001 |
|  | Day 1 vs. Day 15 | -1.850 | 1.955 | 0.780 |
|  | Day 1 vs. Day 60 | -10.284 | 2.229 | <0.001 |
|  | Day 15 vs. Day 60 | -8.434 | 2.229 | 0.004 |
| Clastrum (CLA) | Home cage vs. Day 1 | 0.145 | 2.131 | 1.000 |
|  | Home cage vs. Day 15 | 0.332 | 2.131 | 0.999 |
|  | Homecage vs. Day 60 | -9.499 | 2.474 | 0.003 |
|  | Day 1 vs. Day 15 | 0.187 | 2.293 | 1.000 |
|  | Day 1 vs. Day 60 | -9.644 | 2.615 | 0.005 |
|  | Day 15 vs. Day 60 | -9.831 | 2.615 | 0.004 |
| Lateral septum (LS) | Home cage vs. Day 1 | 0.373 | 4.426 | 1.000 |
|  | Home cage vs. Day 15 | -1.852 | 4.426 | 0.975 |
|  | Homecage vs. Day 60 | -19.497 | 5.138 | 0.004 |
|  | Day 1 vs. Day 15 | -2.225 | 4.763 | 0.966 |
|  | Day 1 vs. Day 60 | -19.869 | 5.431 | 0.005 |
|  | Day 15 vs. Day 60 | -17.644 | 5.431 | 0.015 |
| Nucleus accumbens (ACB) | Home cage vs. Day 1 | 0.920 | 4.006 | 0.996 |
|  | Home cage vs. Day 15 | -0.662 | 4.006 | 0.998 |
|  | Homecage vs. Day 60 | -16.919 | 4.650 | 0.006 |
|  | Day 1 vs. Day 15 | -1.582 | 4.311 | 0.983 |
|  | Day 1 vs. Day 60 | -17.839 | 4.915 | 0.006 |
|  | Day 15 vs. Day 60 | -16.257 | 4.915 | 0.013 |
| Olfactory tubercle (OT) | Home cage vs. Day 1 | 0.638 | 2.081 | 0.990 |
|  | Home cage vs. Day 15 | -0.817 | 2.081 | 0.979 |
|  | Homecage vs. Day 60 | -9.247 | 2.415 | 0.004 |
|  | Day 1 vs. Day 15 | -1.455 | 2.239 | 0.915 |
|  | Day 1 vs. Day 60 | -9.885 | 2.553 | 0.003 |
|  | Day 15 vs. Day 60 | -8.430 | 2.553 | 0.013 |
| Substantia innominata (SI) | Home cage vs. Day 1 | 0.750 | 2.204 | 0.986 |
|  | Home cage vs. Day 15 | -1.217 | 2.204 | 0.945 |
|  | Homecage vs. Day 60 | -8.677 | 2.559 | 0.011 |
|  | Day 1 vs. Day 15 | -1.966 | 2.372 | 0.840 |
|  | Day 1 vs. Day 60 | -9.427 | 2.704 | 0.008 |
|  | Day 15 vs. Day 60 | -7.460 | 2.704 | 0.047 |

**Table S8.**
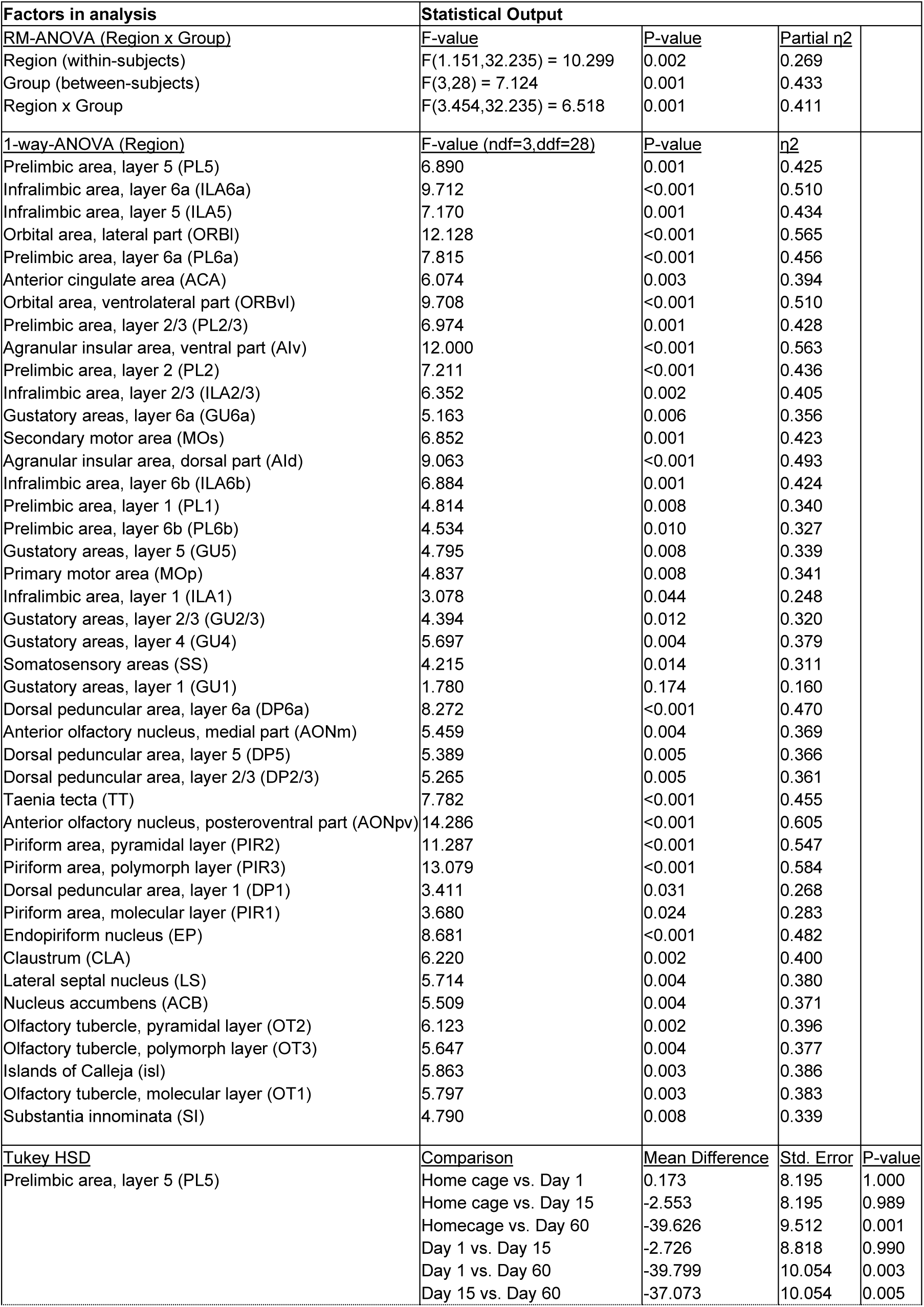
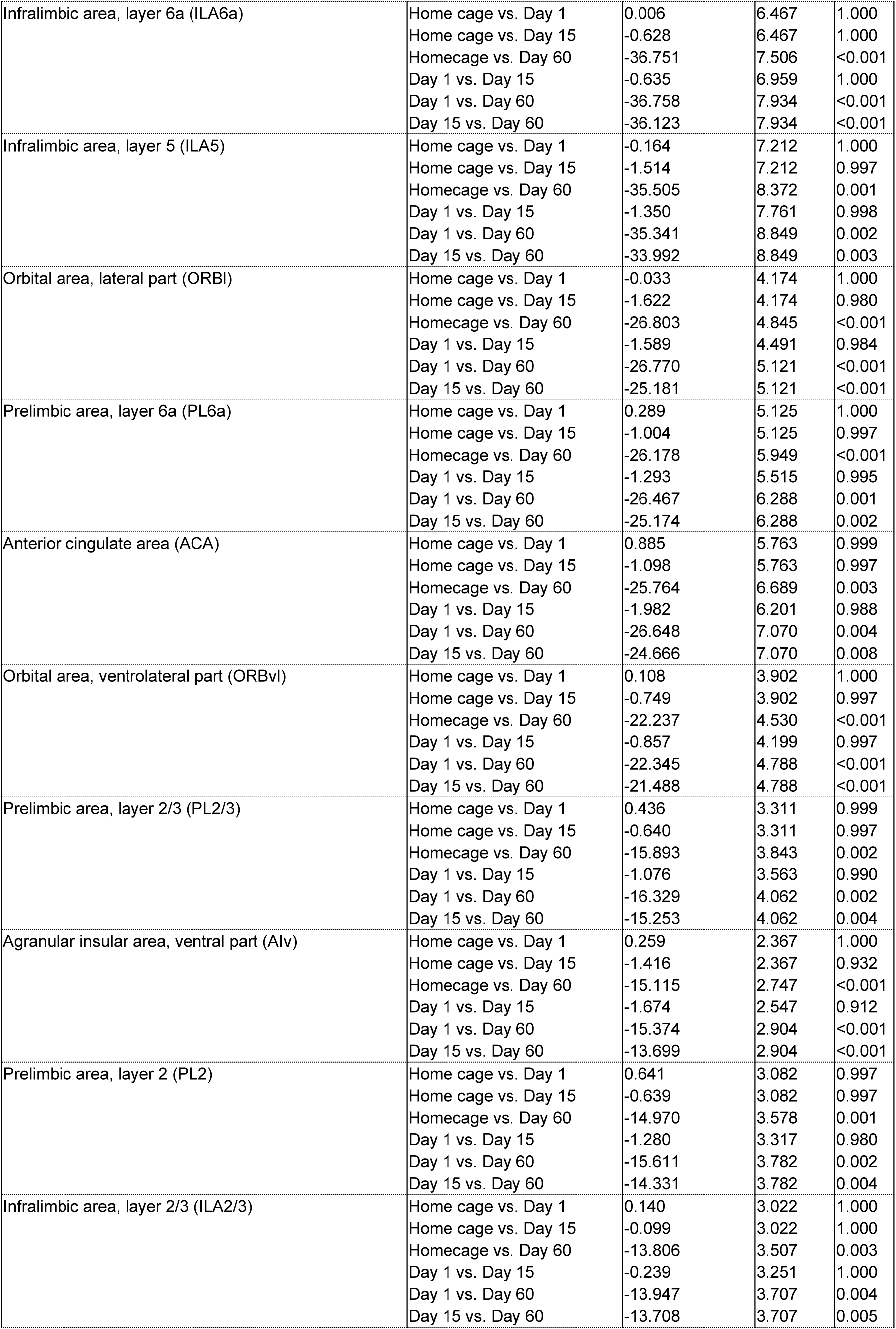
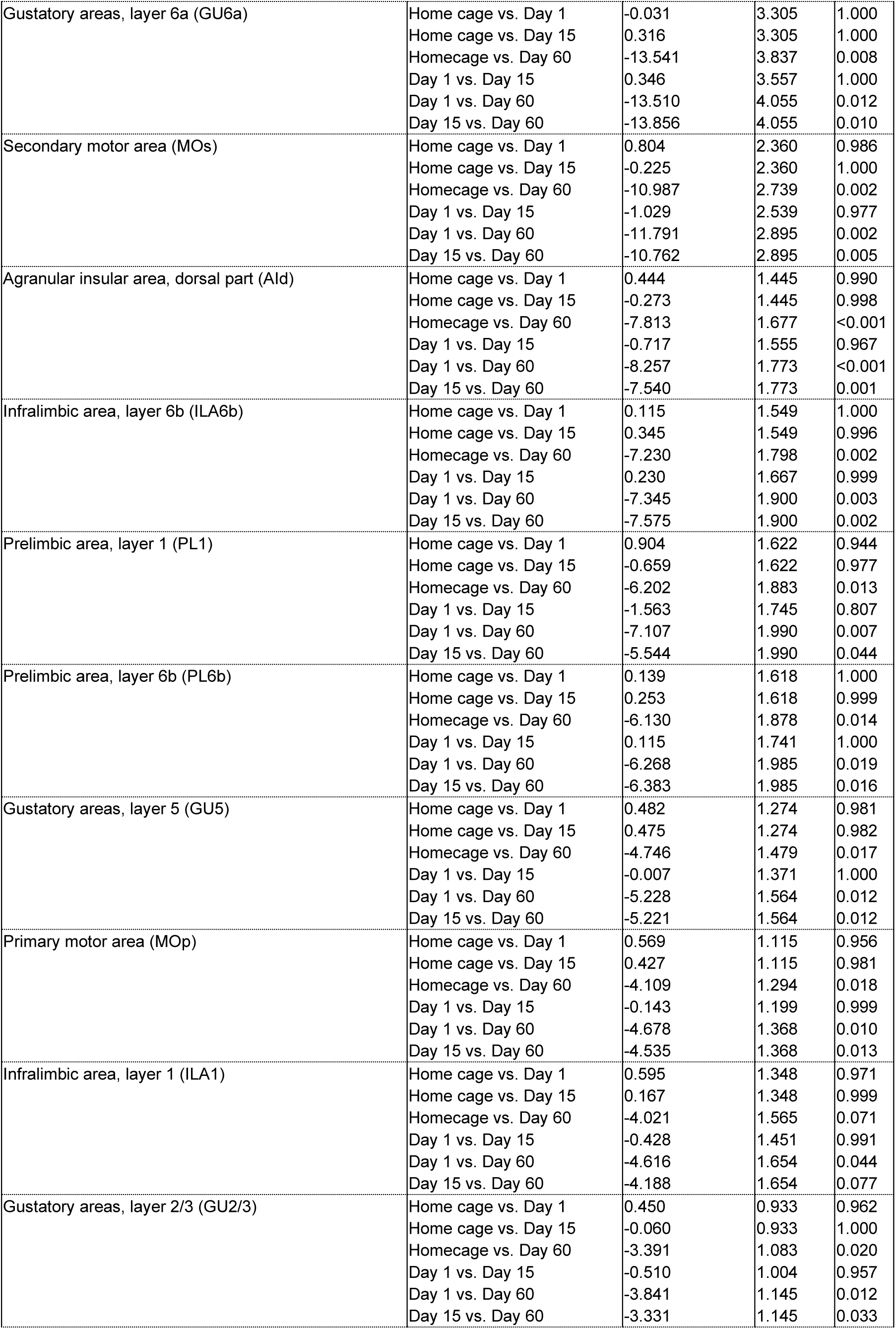
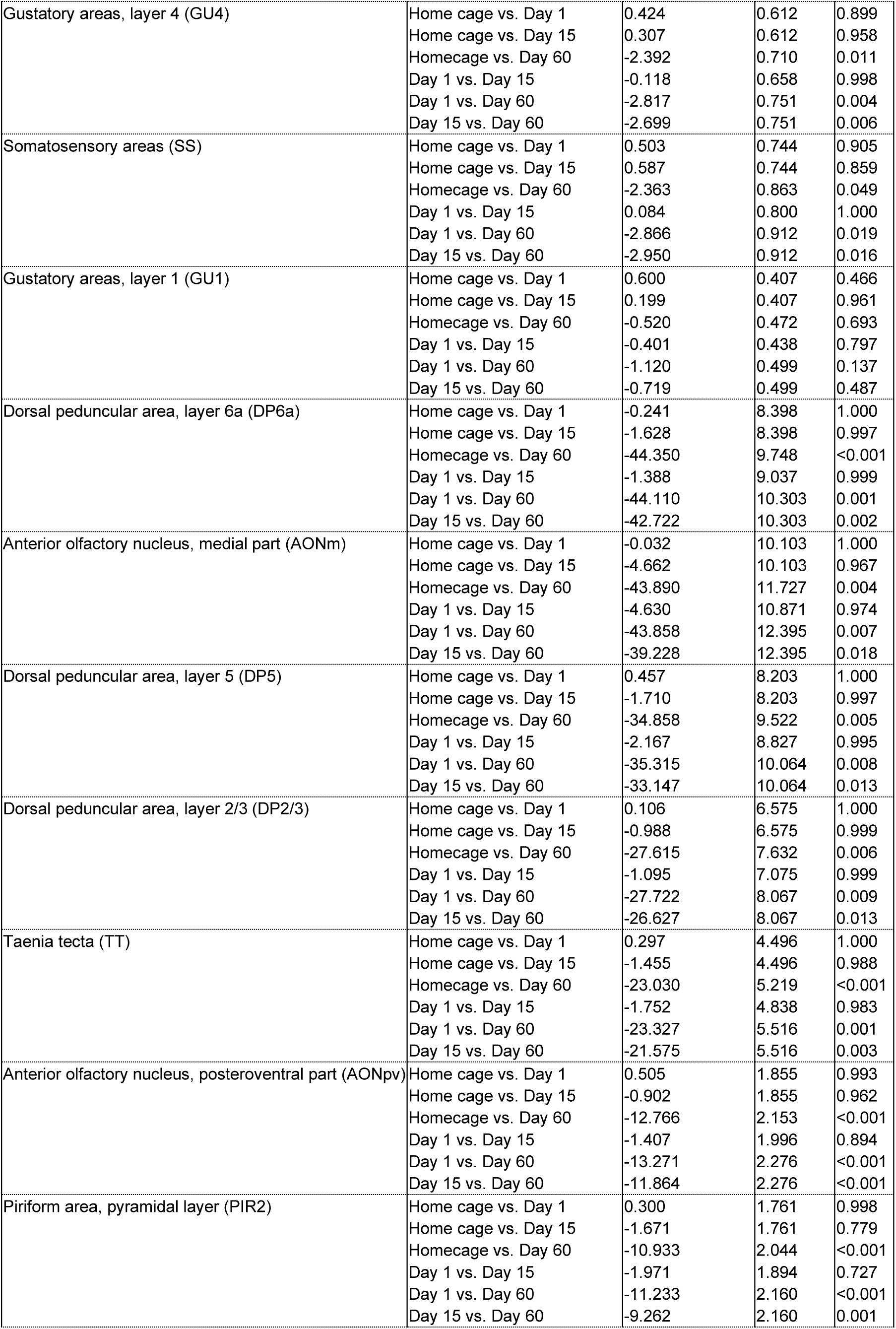
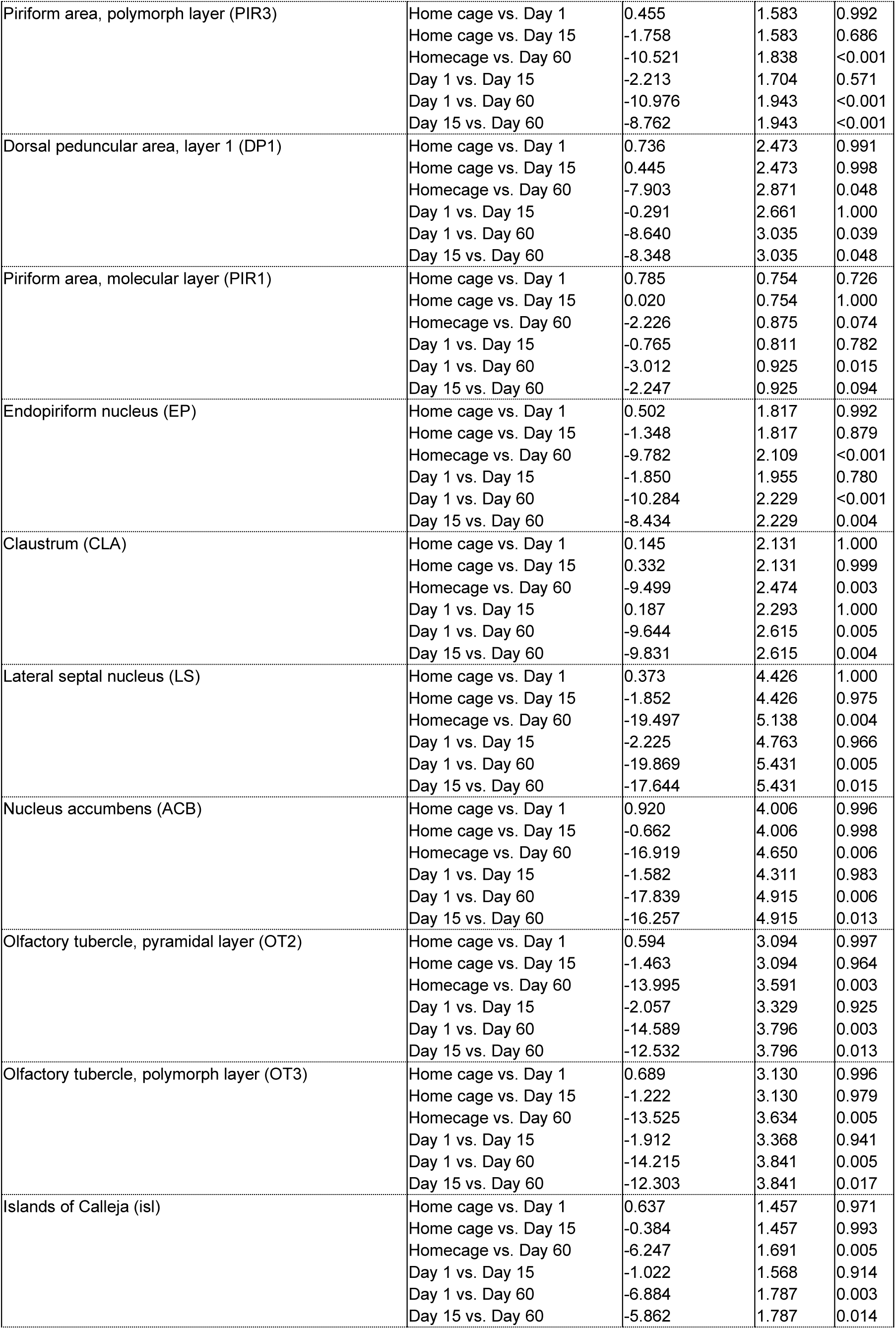
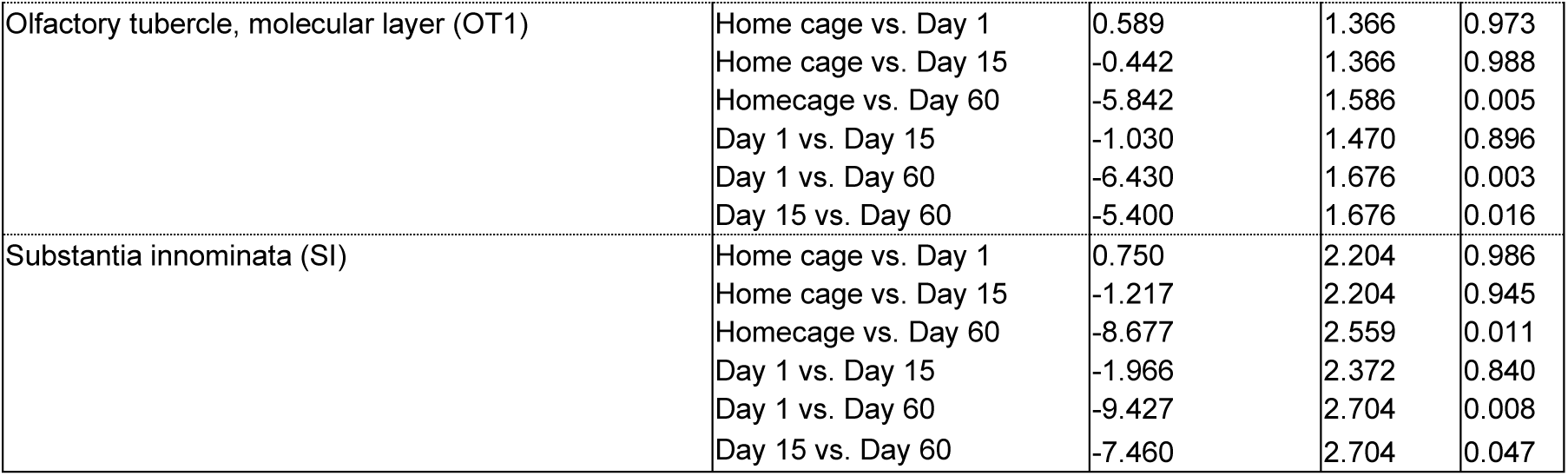
Statistical output for Figure 3D: Analysis of Z-scored counts from 43 subdivisions within AP +1.55 to AP +1.75 coronal subvolume (n=31)

**Table S9.** Statistical output for Figure 4B, right panel : Analysis of Z-scored counts from 8 major anatomical regions within AP -1.08 to AP -1.28 coronal subvolume (n=30, analyses shown in figure 3B are highlighted in grey)

| Factors in analysis | Statistical output |  |  |  |
| --- | --- | --- | --- | --- |
| <u>RM-ANOVA (Region x Group)</u> | <u>F-value</u> | <u>P-value</u> | <u>Partial $\eta^2$</u> |  |
| Region (within-subjects) | F(1.198,32.349) = 14.028 | <0.001 | 0.342 |  |
| Group (between-subjects) | F(3,27) = 3.189 | 0.040 | 0.262 |  |
| Region x Group | F(3.594,32.349) = 5.476 | 0.002 | 0.378 |  |
| <u>1-way-ANOVA (Region)</u> | <u>F-value (ndf=3,ddf=27)</u> | <u>P-value</u> | <u>$\eta^2$</u> |  |
| Isocortex | 4.078 | 0.016 | 0.312 |  |
| Olfactory areas | 5.049 | 0.007 | 0.359 |  |
| Hippocampal formation | 0.754 | 0.530 | 0.077 |  |
| Cortical subplate | 4.772 | 0.009 | 0.346 |  |
| Striatum | 2.085 | 0.126 | 0.188 |  |
| Pallidum | 1.098 | 0.367 | 0.109 |  |
| Thalamus | 0.761 | 0.526 | 0.078 |  |
| Hypothalamus | 2.723 | 0.064 | 0.232 |  |
| <u>Tukey HSD</u> | <u>Comparison</u> | <u>Mean Difference</u> | <u>Std. Error</u> | <u>P-value</u> |
| Isocortex | Home cage vs. Day 1 | 0.309 | 0.910 | 0.986 |
|  | Home cage vs. Day 15 | -0.304 | 0.910 | 0.987 |
|  | Homecage vs. Day 60 | -3.219 | 1.051 | 0.024 |
|  | Day 1 vs. Day 15 | -0.613 | 0.959 | 0.918 |
|  | Day 1 vs. Day 60 | -3.528 | 1.093 | 0.016 |
|  | Day 15 vs. Day 60 | -2.915 | 1.093 | 0.058 |
| Olfactory areas | Home cage vs. Day 1 | 0.215 | 0.883 | 0.995 |
|  | Home cage vs. Day 15 | -1.152 | 0.883 | 0.568 |
|  | Homecage vs. Day 60 | -3.513 | 1.019 | 0.010 |
|  | Day 1 vs. Day 15 | -1.367 | 0.930 | 0.469 |
|  | Day 1 vs. Day 60 | -3.728 | 1.061 | 0.008 |
|  | Day 15 vs. Day 60 | -2.361 | 1.061 | 0.142 |
| Hippocampal formation | Home cage vs. Day 1 | 0.143 | 0.692 | 0.997 |
|  | Home cage vs. Day 15 | 0.417 | 0.692 | 0.930 |
|  | Homecage vs. Day 60 | -0.808 | 0.799 | 0.744 |
|  | Day 1 vs. Day 15 | 0.273 | 0.729 | 0.982 |
|  | Day 1 vs. Day 60 | -0.951 | 0.831 | 0.666 |
|  | Day 15 vs. Day 60 | -1.224 | 0.831 | 0.467 |
| Cortical subplate | Home cage vs. Day 1 | -0.071 | 2.315 | 1.000 |
|  | Home cage vs. Day 15 | -2.517 | 2.315 | 0.700 |
|  | Homecage vs. Day 60 | -9.349 | 2.673 | 0.008 |
|  | Day 1 vs. Day 15 | -2.446 | 2.440 | 0.749 |
|  | Day 1 vs. Day 60 | -9.278 | 2.782 | 0.013 |
|  | Day 15 vs. Day 60 | -6.832 | 2.782 | 0.090 |
| Striatum | Home cage vs. Day 1 | 0.245 | 0.655 | 0.982 |
|  | Home cage vs. Day 15 | -0.075 | 0.655 | 0.999 |
|  | Homecage vs. Day 60 | -1.602 | 0.757 | 0.173 |
|  | Day 1 vs. Day 15 | -0.320 | 0.691 | 0.966 |
|  | Day 1 vs. Day 60 | -1.847 | 0.787 | 0.113 |
|  | Day 15 vs. Day 60 | -1.527 | 0.787 | 0.236 |
| Pallidum | Home cage vs. Day 1 | 0.248 | 0.507 | 0.961 |
|  | Home cage vs. Day 15 | 0.478 | 0.507 | 0.782 |
|  | Homecage vs. Day 60 | -0.585 | 0.586 | 0.751 |
|  | Day 1 vs. Day 15 | 0.231 | 0.535 | 0.973 |
|  | Day 1 vs. Day 60 | -0.833 | 0.610 | 0.531 |
|  | Day 15 vs. Day 60 | -1.064 | 0.610 | 0.321 |
| Thalamus | Home cage vs. Day 1 | 0.071 | 0.385 | 0.998 |
|  | Home cage vs. Day 15 | 0.412 | 0.385 | 0.710 |
|  | Homecage vs. Day 60 | -0.253 | 0.444 | 0.940 |
|  | Day 1 vs. Day 15 | 0.341 | 0.405 | 0.834 |
|  | Day 1 vs. Day 60 | -0.324 | 0.462 | 0.896 |
|  | Day 15 vs. Day 60 | -0.665 | 0.462 | 0.487 |
| Hypothalamus | Home cage vs. Day 1 | 0.301 | 1.133 | 0.993 |
|  | Home cage vs. Day 15 | -0.862 | 1.133 | 0.871 |
|  | Homecage vs. Day 60 | -3.289 | 1.308 | 0.080 |
|  | Day 1 vs. Day 15 | -1.162 | 1.194 | 0.766 |
|  | Day 1 vs. Day 60 | -3.589 | 1.362 | 0.062 |
|  | Day 15 vs. Day 60 | -2.427 | 1.362 | 0.303 |

**Table S10.** Statistical output for Figure 4C: Analysis of Z-scored counts from 28 subregions within AP -1.08 to AP -1.28 coronal subvolume (n=30)

| Factors in analysis | Statistical Output |  |  |  |
| --- | --- | --- | --- | --- |
| **RM-ANOVA (Region x Group)** | **F-value** | **P-value** | **Partial $\eta^2$** |  |
| Region (within-subjects) | F(1.140,30.793) = 9.528 | 0.003 | 0.261 |  |
| Group (between-subjects) | F(3,27) = 3.373 | 0.033 | 0.273 |  |
| Region x Group | F(3.421,30.793) = 3.453 | 0.024 | 0.277 |  |
| **1-way-ANOVA (Region)** | **F-value (ndf=3,ddf=27)** | **P-value** | **$\eta^2$** |  |
| Somatomotor areas (MO) | 3.936 | 0.019 | 0.304 |  |
| Retrosplenial area (RSP) | 1.904 | 0.153 | 0.175 |  |
| Somatosensory areas (SS) | 4.377 | 0.012 | 0.327 |  |
| Anterior cingulate area (ACA) | 1.580 | 0.217 | 0.149 |  |
| Agranular insular area (AI) | 2.042 | 0.132 | 0.185 |  |
| Visceral area (VISC) | 1.943 | 0.146 | 0.178 |  |
| Cortical amygdalar area (COA) | 6.456 | 0.002 | 0.418 |  |
| Piriform area (PIR) | 4.306 | 0.013 | 0.324 |  |
| Piriform-amygdalar area (PAA) | 3.508 | 0.029 | 0.280 |  |
| Ammon's horn (CA) | 0.832 | 0.488 | 0.085 |  |
| Dentate gyrus (DG) | 0.683 | 0.570 | 0.071 |  |
| Basomedial amygdalar nucleus (BMA) | 4.061 | 0.017 | 0.311 |  |
| Endopiriform nucleus (EP) | 5.828 | 0.003 | 0.393 |  |
| Basolateral amygdalar nucleus (BLA) | 3.638 | 0.025 | 0.288 |  |
| Lateral amygdalar nucleus (LA) | 3.728 | 0.023 | 0.293 |  |
| Clastrum (CLA) | 7.956 | <0.001 | 0.469 |  |
| Intercalated amygdalar nucleus (IA) | 4.051 | 0.017 | 0.310 |  |
| Central amygdalar nucleus (CEA) | 3.465 | 0.030 | 0.278 |  |
| Medial amygdalar nucleus (MEA) | 2.867 | 0.055 | 0.242 |  |
| Bed nucleus of the accessory olfactory tract (BA) | 1.560 | 0.222 | 0.148 |  |
| Caudoputamen (CP) | 1.126 | 0.356 | 0.111 |  |
| Pallidum, ventral region (PALv) | 1.535 | 0.228 | 0.146 |  |
| Pallidum, dorsal region (PALd) | 0.835 | 0.487 | 0.085 |  |
| Thalamus, sensory-motor cortex related (DORsm) | 0.726 | 0.545 | 0.075 |  |
| Thalamus, polymodal association cortex related (DORpm) | 0.786 | 0.512 | 0.080 |  |
| Hypothalamic medial zone (MEZ) | 2.509 | 0.080 | 0.218 |  |
| Hypothalamic lateral zone (LZ) | 2.677 | 0.067 | 0.229 |  |
| Periventricular zone (PVZ) | 2.933 | 0.051 | 0.246 |  |
| **Tukey HSD** | **Comparison** | **Mean Difference** | **Std. Error** | **P-value** |
| Somatomotor areas (MO) | Home cage vs. Day 1 | 0.523 | 1.555 | 0.987 |
|  | Home cage vs. Day 15 | -0.803 | 1.555 | 0.954 |
|  | Homecage vs. Day 60 | -5.428 | 1.795 | 0.026 |
|  | Day 1 vs. Day 15 | -1.326 | 1.639 | 0.849 |
|  | Day 1 vs. Day 60 | -5.951 | 1.868 | 0.018 |
|  | Day 15 vs. Day 60 | -4.625 | 1.868 | 0.087 |
| Retrosplenial area (RSP) | Home cage vs. Day 1 | 0.264 | 1.817 | 0.999 |
|  | Home cage vs. Day 15 | -0.350 | 1.817 | 0.997 |
|  | Homecage vs. Day 60 | -4.456 | 2.098 | 0.171 |
|  | Day 1 vs. Day 15 | -0.614 | 1.915 | 0.988 |
|  | Day 1 vs. Day 60 | -4.720 | 2.184 | 0.160 |
|  | Day 15 vs. Day 60 | -4.106 | 2.184 | 0.260 |
| Somatosensory areas (SS) | Home cage vs. Day 1 | 0.274 | 0.928 | 0.991 |
|  | Home cage vs. Day 15 | -0.343 | 0.928 | 0.982 |
|  | Homecage vs. Day 60 | -3.431 | 1.072 | 0.017 |
|  | Day 1 vs. Day 15 | -0.617 | 0.978 | 0.921 |
|  | Day 1 vs. Day 60 | -3.705 | 1.116 | 0.013 |
|  | Day 15 vs. Day 60 | -3.088 | 1.116 | 0.047 |
| Anterior cingulate area (ACA) | Home cage vs. Day 1 | 0.238 | 1.115 | 0.996 |
|  | Home cage vs. Day 15 | -0.347 | 1.115 | 0.989 |
|  | Homecage vs. Day 60 | -2.465 | 1.287 | 0.246 |
|  | Day 1 vs. Day 15 | -0.585 | 1.175 | 0.959 |
|  | Day 1 vs. Day 60 | -2.703 | 1.340 | 0.207 |
|  | Day 15 vs. Day 60 | -2.118 | 1.340 | 0.406 |
| Agranular insular area (AI) | Home cage vs. Day 1 | 0.354 | 0.425 | 0.838 |
|  | Home cage vs. Day 15 | 0.148 | 0.425 | 0.985 |
|  | Homecage vs. Day 60 | -0.864 | 0.490 | 0.313 |
|  | Day 1 vs. Day 15 | -0.206 | 0.448 | 0.967 |
|  | Day 1 vs. Day 60 | -1.218 | 0.510 | 0.104 |
|  | Day 15 vs. Day 60 | -1.012 | 0.510 | 0.219 |
| Visceral area (VISC) | Home cage vs. Day 1 | 0.342 | 0.400 | 0.828 |
|  | Home cage vs. Day 15 | 0.162 | 0.400 | 0.977 |
|  | Homecage vs. Day 60 | -0.776 | 0.462 | 0.355 |
|  | Day 1 vs. Day 15 | -0.181 | 0.422 | 0.973 |
|  | Day 1 vs. Day 60 | -1.118 | 0.481 | 0.118 |
|  | Day 15 vs. Day 60 | -0.937 | 0.481 | 0.233 |
| Cortical amygdalar area (COA) | Home cage vs. Day 1 | -0.030 | 1.080 | 1.000 |
|  | Home cage vs. Day 15 | -2.262 | 1.080 | 0.180 |
|  | Homecage vs. Day 60 | -4.858 | 1.247 | 0.003 |
|  | Day 1 vs. Day 15 | -2.232 | 1.138 | 0.227 |
|  | Day 1 vs. Day 60 | -4.828 | 1.297 | 0.005 |
|  | Day 15 vs. Day 60 | -2.596 | 1.297 | 0.213 |
| Piriform area (PIR) | Home cage vs. Day 1 | 0.278 | 0.852 | 0.988 |
|  | Home cage vs. Day 15 | -0.735 | 0.852 | 0.824 |
|  | Homecage vs. Day 60 | -3.122 | 0.984 | 0.018 |
|  | Day 1 vs. Day 15 | -1.013 | 0.898 | 0.676 |
|  | Day 1 vs. Day 60 | -3.400 | 1.024 | 0.013 |
|  | Day 15 vs. Day 60 | -2.387 | 1.024 | 0.116 |
| Piriform-amygdalar area (PAA) | Home cage vs. Day 1 | 0.434 | 0.709 | 0.927 |
|  | Home cage vs. Day 15 | -0.660 | 0.709 | 0.789 |
|  | Homecage vs. Day 60 | -2.178 | 0.819 | 0.059 |
|  | Day 1 vs. Day 15 | -1.094 | 0.747 | 0.472 |
|  | Day 1 vs. Day 60 | -2.612 | 0.852 | 0.024 |
|  | Day 15 vs. Day 60 | -1.518 | 0.852 | 0.304 |
| Ammon's horn (CA) | Home cage vs. Day 1 | 0.068 | 0.820 | 1.000 |
|  | Home cage vs. Day 15 | 0.345 | 0.820 | 0.975 |
|  | Homecage vs. Day 60 | -1.155 | 0.947 | 0.620 |
|  | Day 1 vs. Day 15 | 0.277 | 0.865 | 0.988 |
|  | Day 1 vs. Day 60 | -1.223 | 0.986 | 0.607 |
|  | Day 15 vs. Day 60 | -1.500 | 0.986 | 0.439 |
| Dentate gyrus (DG) | Home cage vs. Day 1 | 0.288 | 0.453 | 0.920 |
|  | Home cage vs. Day 15 | 0.547 | 0.453 | 0.628 |
|  | Homecage vs. Day 60 | -0.098 | 0.523 | 0.998 |
|  | Day 1 vs. Day 15 | 0.259 | 0.478 | 0.948 |
|  | Day 1 vs. Day 60 | -0.386 | 0.545 | 0.893 |
|  | Day 15 vs. Day 60 | -0.645 | 0.545 | 0.641 |
| Basomedial amygdalar nucleus (BMA) | Home cage vs. Day 1 | -1.346 | 5.743 | 0.995 |
|  | Home cage vs. Day 15 | -7.829 | 5.743 | 0.532 |
|  | Homecage vs. Day 60 | -21.664 | 6.632 | 0.015 |
|  | Day 1 vs. Day 15 | -6.483 | 6.054 | 0.710 |
|  | Day 1 vs. Day 60 | -20.318 | 6.903 | 0.031 |
|  | Day 15 vs. Day 60 | -13.835 | 6.903 | 0.211 |
| Endopiriform nucleus (EP) | Home cage vs. Day 1 | 0.136 | 1.601 | 1.000 |
|  | Home cage vs. Day 15 | -1.252 | 1.601 | 0.862 |
|  | Homecage vs. Day 60 | -7.064 | 1.848 | 0.004 |
|  | Day 1 vs. Day 15 | -1.388 | 1.687 | 0.843 |
|  | Day 1 vs. Day 60 | -7.200 | 1.924 | 0.005 |
|  | Day 15 vs. Day 60 | -5.812 | 1.924 | 0.026 |
| Basolateral amygdalar nucleus (BLA) | Home cage vs. Day 1 | 0.386 | 2.051 | 0.998 |
|  | Home cage vs. Day 15 | -1.199 | 2.051 | 0.936 |
|  | Homecage vs. Day 60 | -7.035 | 2.369 | 0.030 |
|  | Day 1 vs. Day 15 | -1.585 | 2.162 | 0.883 |
|  | Day 1 vs. Day 60 | -7.421 | 2.465 | 0.027 |
|  | Day 15 vs. Day 60 | -5.836 | 2.465 | 0.108 |
| Lateral amygdalar nucleus (LA) | Home cage vs. Day 1 | 0.642 | 0.867 | 0.880 |
|  | Home cage vs. Day 15 | -0.399 | 0.867 | 0.967 |
|  | Homecage vs. Day 60 | -2.729 | 1.002 | 0.051 |
|  | Day 1 vs. Day 15 | -1.041 | 0.914 | 0.670 |
|  | Day 1 vs. Day 60 | -3.371 | 1.043 | 0.016 |
|  | Day 15 vs. Day 60 | -2.330 | 1.043 | 0.139 |
| Clastrum (CLA) | Home cage vs. Day 1 | 0.332 | 0.529 | 0.922 |
|  | Home cage vs. Day 15 | -0.473 | 0.529 | 0.808 |
|  | Homecage vs. Day 60 | -2.585 | 0.611 | 0.001 |
|  | Day 1 vs. Day 15 | -0.805 | 0.558 | 0.485 |
|  | Day 1 vs. Day 60 | -2.917 | 0.636 | <0.001 |
|  | Day 15 vs. Day 60 | -2.112 | 0.636 | 0.013 |
| Intercalated amygdalar nucleus (IA) | Home cage vs. Day 1 | -1.038 | 4.303 | 0.995 |
|  | Home cage vs. Day 15 | -4.797 | 4.303 | 0.684 |
|  | Homecage vs. Day 60 | -16.372 | 4.968 | 0.014 |
|  | Day 1 vs. Day 15 | -3.758 | 4.535 | 0.840 |
|  | Day 1 vs. Day 60 | -15.334 | 5.171 | 0.030 |
|  | Day 15 vs. Day 60 | -11.576 | 5.171 | 0.138 |
| Central amygdalar nucleus (CEA) | Home cage vs. Day 1 | 0.306 | 1.325 | 0.996 |
|  | Home cage vs. Day 15 | -0.553 | 1.325 | 0.975 |
|  | Homecage vs. Day 60 | -4.391 | 1.530 | 0.037 |
|  | Day 1 vs. Day 15 | -0.859 | 1.397 | 0.926 |
|  | Day 1 vs. Day 60 | -4.698 | 1.593 | 0.031 |
|  | Day 15 vs. Day 60 | -3.838 | 1.593 | 0.099 |
| Medial amygdalar nucleus (MEA) | Home cage vs. Day 1 | 0.156 | 1.389 | 0.999 |
|  | Home cage vs. Day 15 | -1.515 | 1.389 | 0.698 |
|  | Homecage vs. Day 60 | -4.193 | 1.604 | 0.065 |
|  | Day 1 vs. Day 15 | -1.671 | 1.464 | 0.668 |
|  | Day 1 vs. Day 60 | -4.349 | 1.669 | 0.066 |
|  | Day 15 vs. Day 60 | -2.677 | 1.669 | 0.393 |
| Bed nucleus of the accessory olfactory tract (BA) | Home cage vs. Day 1 | 0.077 | 0.695 | 0.999 |
|  | Home cage vs. Day 15 | -0.702 | 0.695 | 0.745 |
|  | Homecage vs. Day 60 | -1.485 | 0.802 | 0.272 |
|  | Day 1 vs. Day 15 | -0.779 | 0.732 | 0.714 |
|  | Day 1 vs. Day 60 | -1.562 | 0.835 | 0.264 |
|  | Day 15 vs. Day 60 | -0.783 | 0.835 | 0.785 |
| Caudoputamen (CP) | Home cage vs. Day 1 | 0.277 | 0.369 | 0.875 |
|  | Home cage vs. Day 15 | 0.447 | 0.369 | 0.625 |
|  | Homecage vs. Day 60 | -0.294 | 0.426 | 0.900 |
|  | Day 1 vs. Day 15 | 0.170 | 0.389 | 0.971 |
|  | Day 1 vs. Day 60 | -0.571 | 0.443 | 0.578 |
|  | Day 15 vs. Day 60 | -0.741 | 0.443 | 0.358 |
| Pallidum, ventral region (PALv) | Home cage vs. Day 1 | 0.452 | 0.706 | 0.918 |
|  | Home cage vs. Day 15 | 0.546 | 0.706 | 0.865 |
|  | Homecage vs. Day 60 | -1.127 | 0.815 | 0.520 |
|  | Day 1 vs. Day 15 | 0.095 | 0.744 | 0.999 |
|  | Day 1 vs. Day 60 | -1.579 | 0.848 | 0.268 |
|  | Day 15 vs. Day 60 | -1.673 | 0.848 | 0.223 |
| Pallidum, dorsal region (PALd) | Home cage vs. Day 1 | 0.167 | 0.443 | 0.981 |
|  | Home cage vs. Day 15 | 0.441 | 0.443 | 0.753 |
|  | Homecage vs. Day 60 | -0.373 | 0.512 | 0.885 |
|  | Day 1 vs. Day 15 | 0.275 | 0.467 | 0.935 |
|  | Day 1 vs. Day 60 | -0.540 | 0.533 | 0.743 |
|  | Day 15 vs. Day 60 | -0.814 | 0.533 | 0.435 |
| Thalamus, sensory-motor cortex related (DORsm) | Home cage vs. Day 1 | -0.044 | 0.433 | 1.000 |
|  | Home cage vs. Day 15 | 0.367 | 0.433 | 0.831 |
|  | Homecage vs. Day 60 | -0.384 | 0.500 | 0.868 |
|  | Day 1 vs. Day 15 | 0.411 | 0.456 | 0.805 |
|  | Day 1 vs. Day 60 | -0.340 | 0.520 | 0.914 |
|  | Day 15 vs. Day 60 | -0.750 | 0.520 | 0.485 |
| Thalamus, polymodal association cortex related (DORpm) | Home cage vs. Day 1 | 0.115 | 0.368 | 0.989 |
|  | Home cage vs. Day 15 | 0.429 | 0.368 | 0.654 |
|  | Homecage vs. Day 60 | -0.203 | 0.425 | 0.964 |
|  | Day 1 vs. Day 15 | 0.314 | 0.388 | 0.850 |
|  | Day 1 vs. Day 60 | -0.317 | 0.443 | 0.890 |
|  | Day 15 vs. Day 60 | -0.631 | 0.443 | 0.495 |
| Hypothalamic medial zone (MEZ) | Home cage vs. Day 1 | 0.011 | 5.940 | 1.000 |
|  | Home cage vs. Day 15 | -4.111 | 5.940 | 0.899 |
|  | Homecage vs. Day 60 | -17.359 | 6.859 | 0.078 |
|  | Day 1 vs. Day 15 | -4.122 | 6.261 | 0.912 |
|  | Day 1 vs. Day 60 | -17.369 | 7.139 | 0.095 |
|  | Day 15 vs. Day 60 | -13.248 | 7.139 | 0.271 |
| Hypothalamic lateral zone (LZ) | Home cage vs. Day 1 | 0.303 | 0.979 | 0.989 |
|  | Home cage vs. Day 15 | -0.737 | 0.979 | 0.875 |
|  | Homecage vs. Day 60 | -2.790 | 1.131 | 0.088 |
|  | Day 1 vs. Day 15 | -1.040 | 1.032 | 0.746 |
|  | Day 1 vs. Day 60 | -3.093 | 1.177 | 0.063 |
|  | Day 15 vs. Day 60 | -2.053 | 1.177 | 0.321 |
| Periventricular zone (PVZ) | Home cage vs. Day 1 | 0.329 | 0.763 | 0.973 |
|  | Home cage vs. Day 15 | -0.667 | 0.763 | 0.818 |
|  | Homecage vs. Day 60 | -2.211 | 0.881 | 0.081 |
|  | Day 1 vs. Day 15 | -0.996 | 0.804 | 0.609 |
|  | Day 1 vs. Day 60 | -2.540 | 0.917 | 0.046 |
|  | Day 15 vs. Day 60 | -1.545 | 0.917 | 0.351 |

**Table S11.**
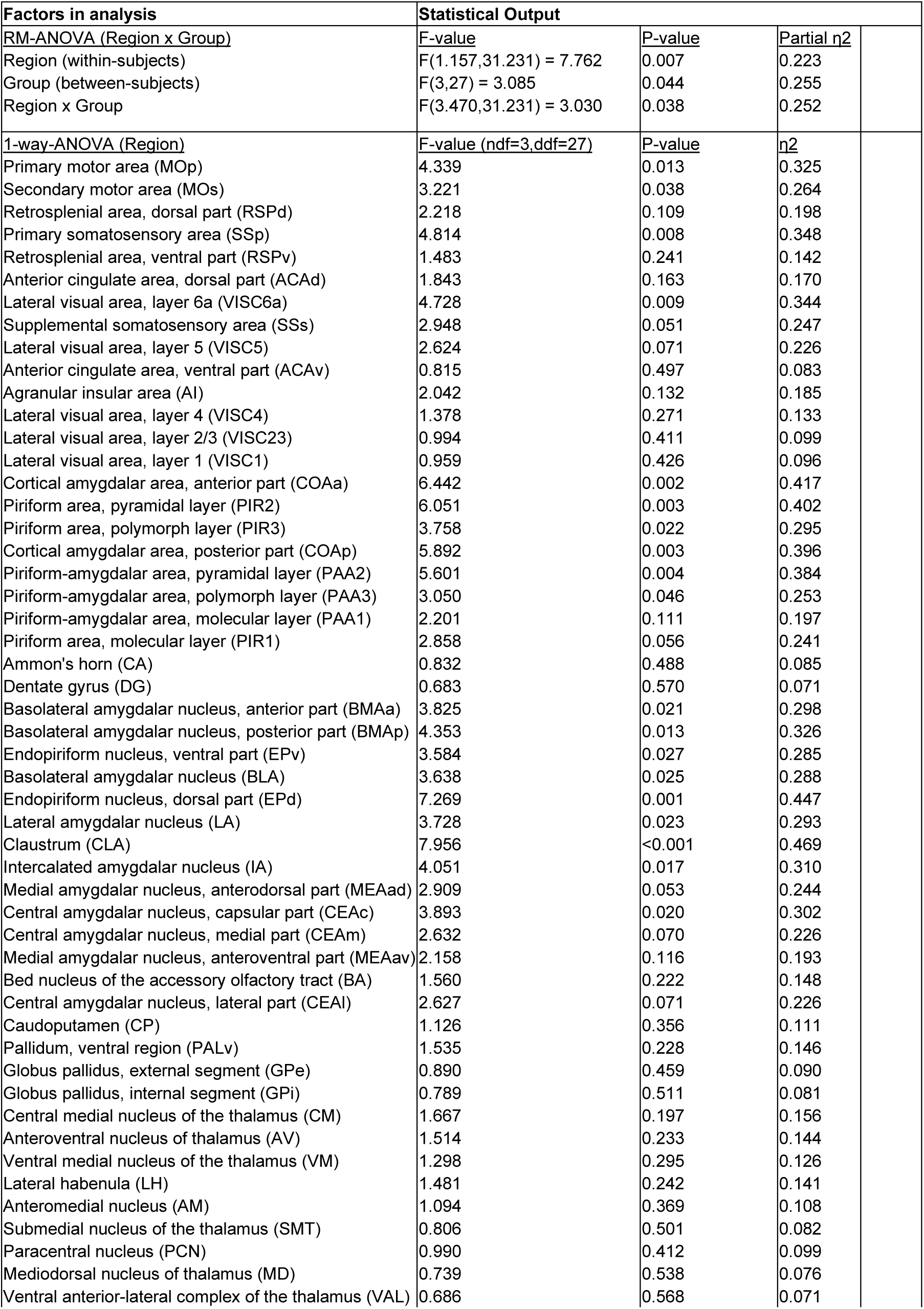
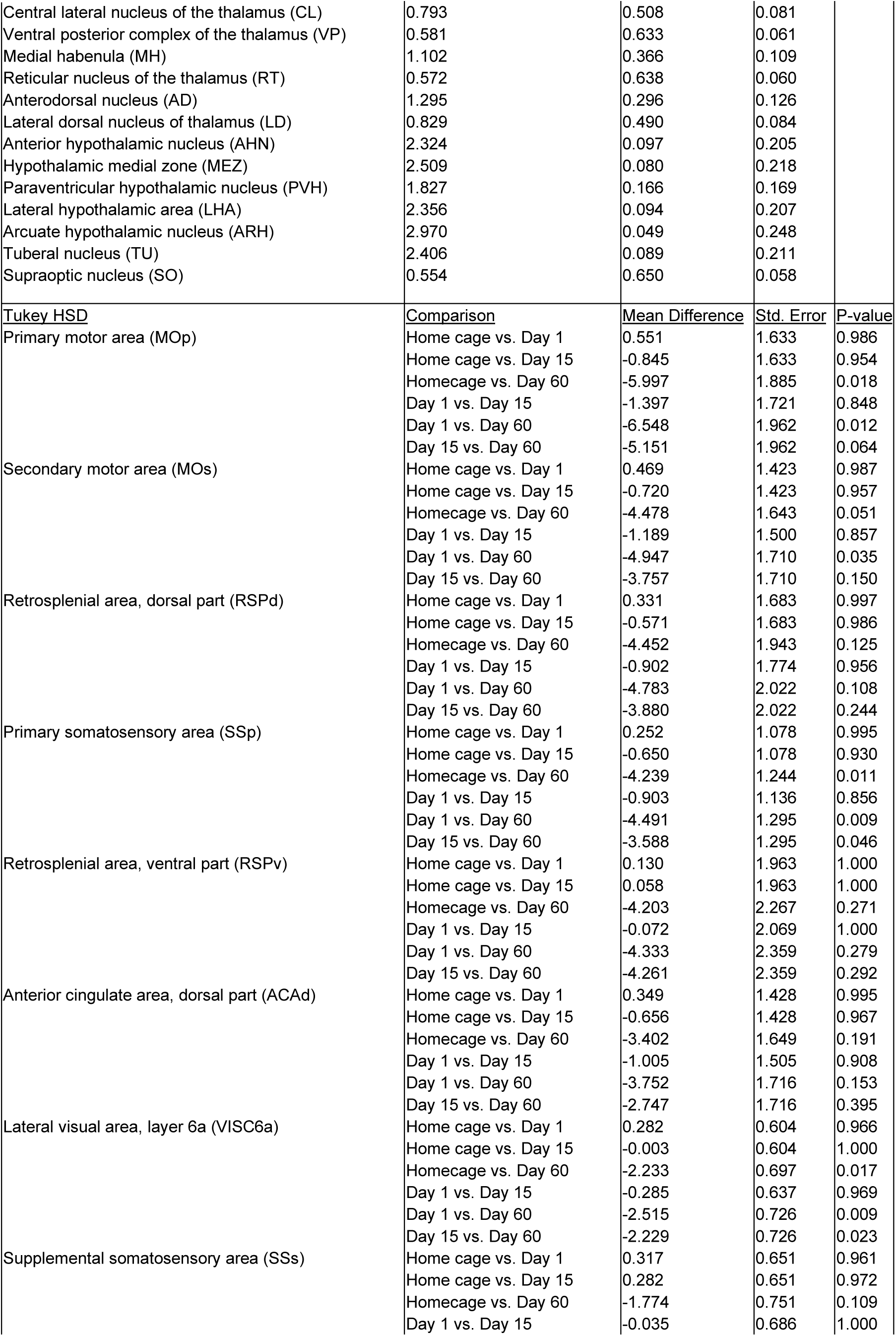
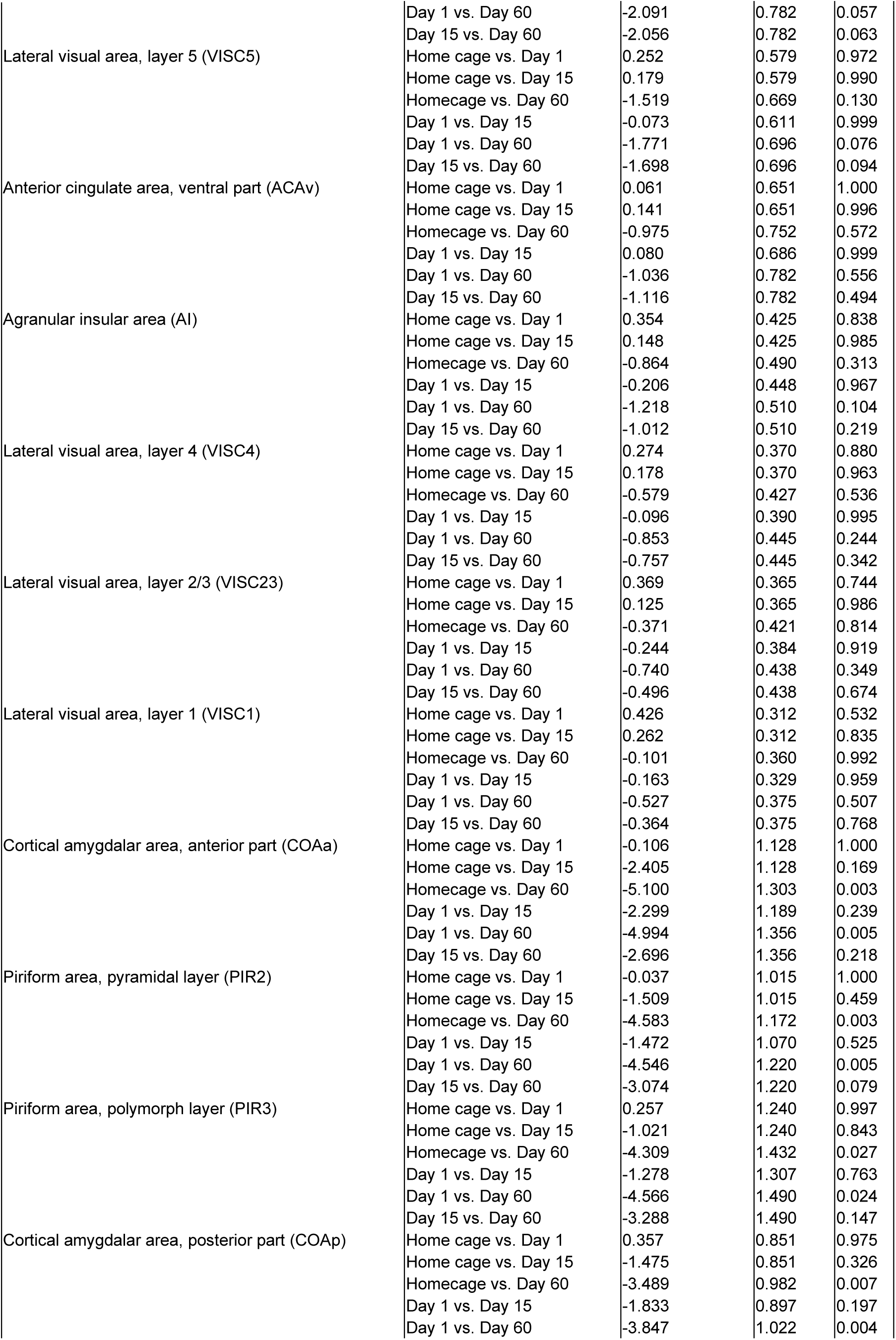
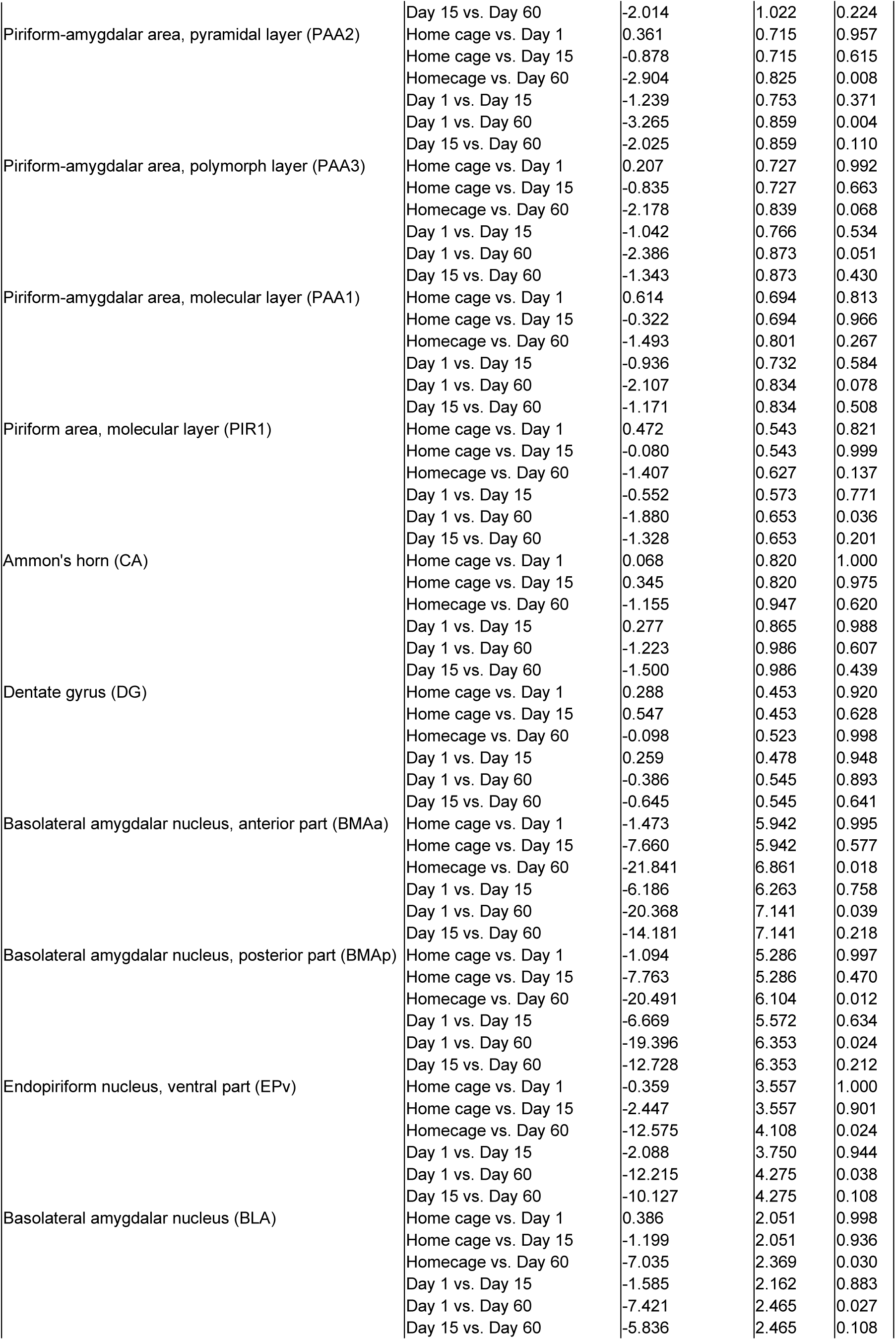
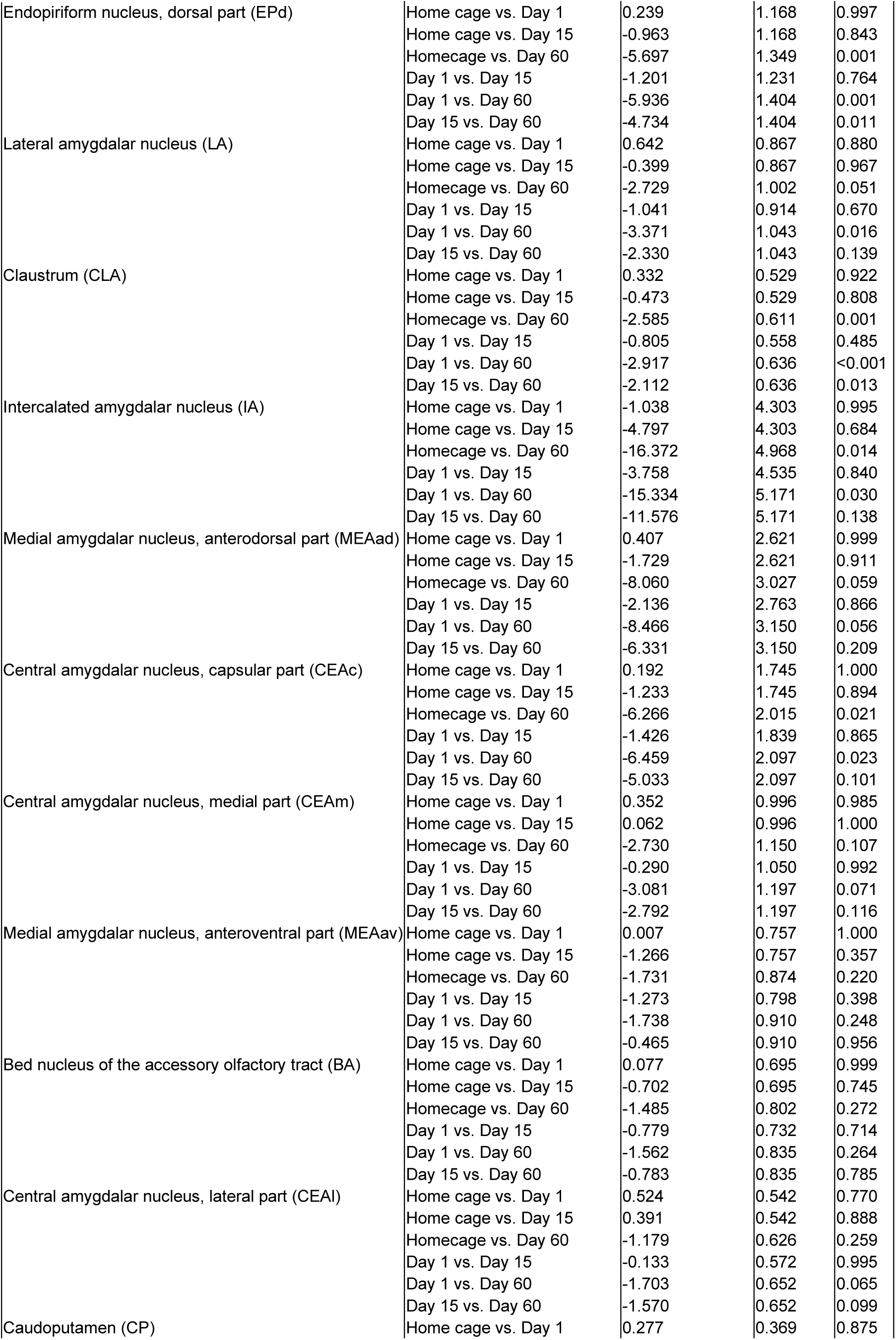
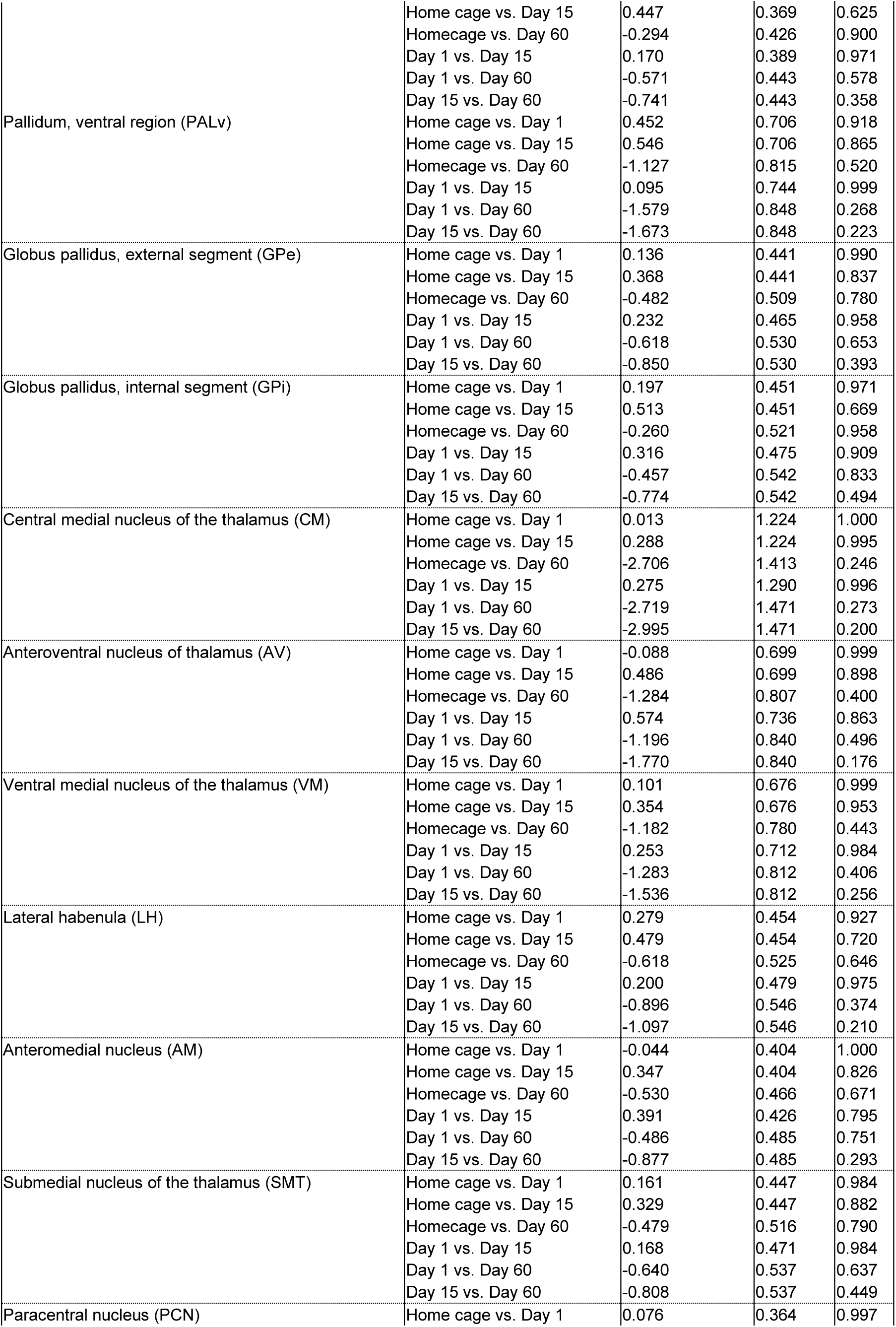
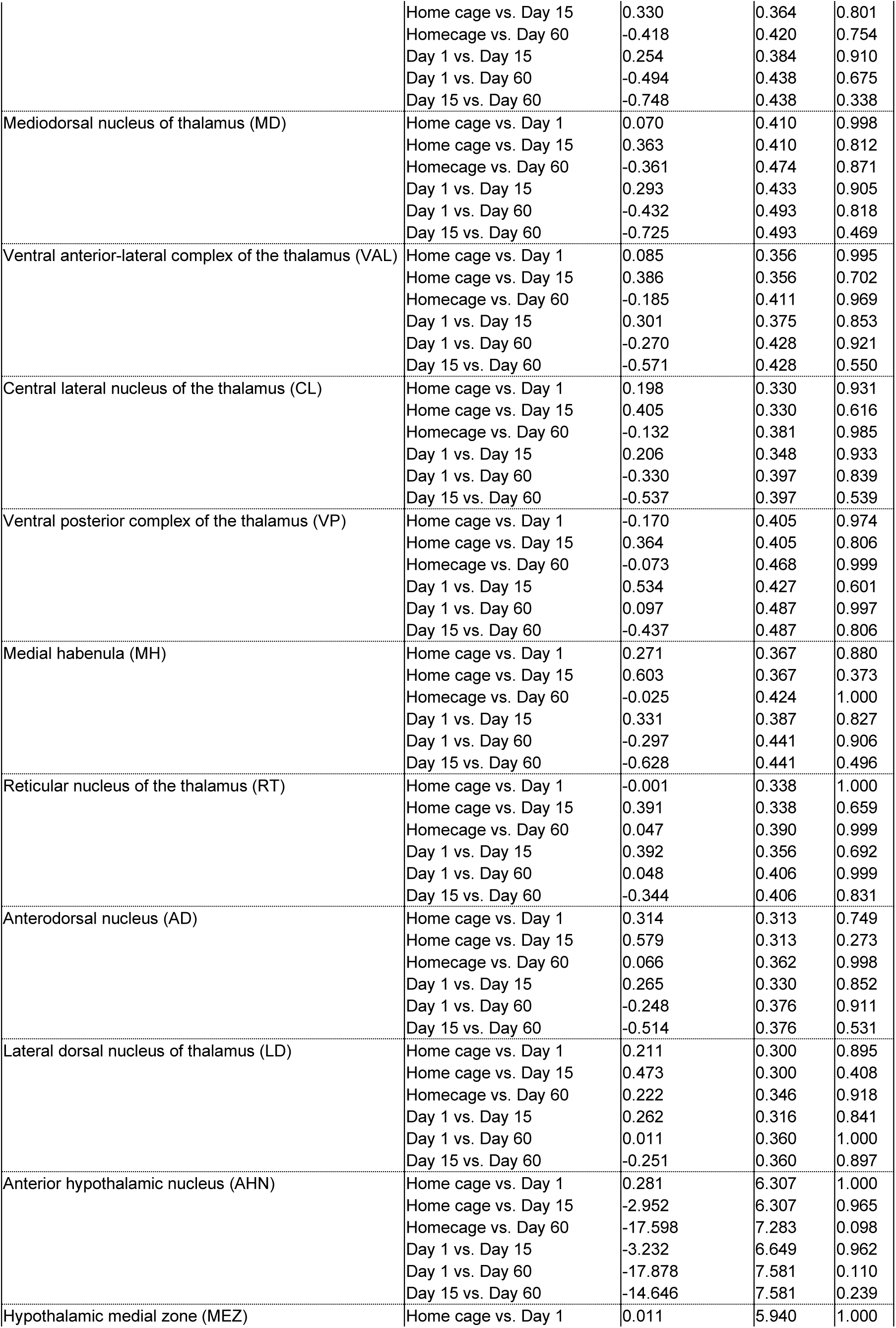
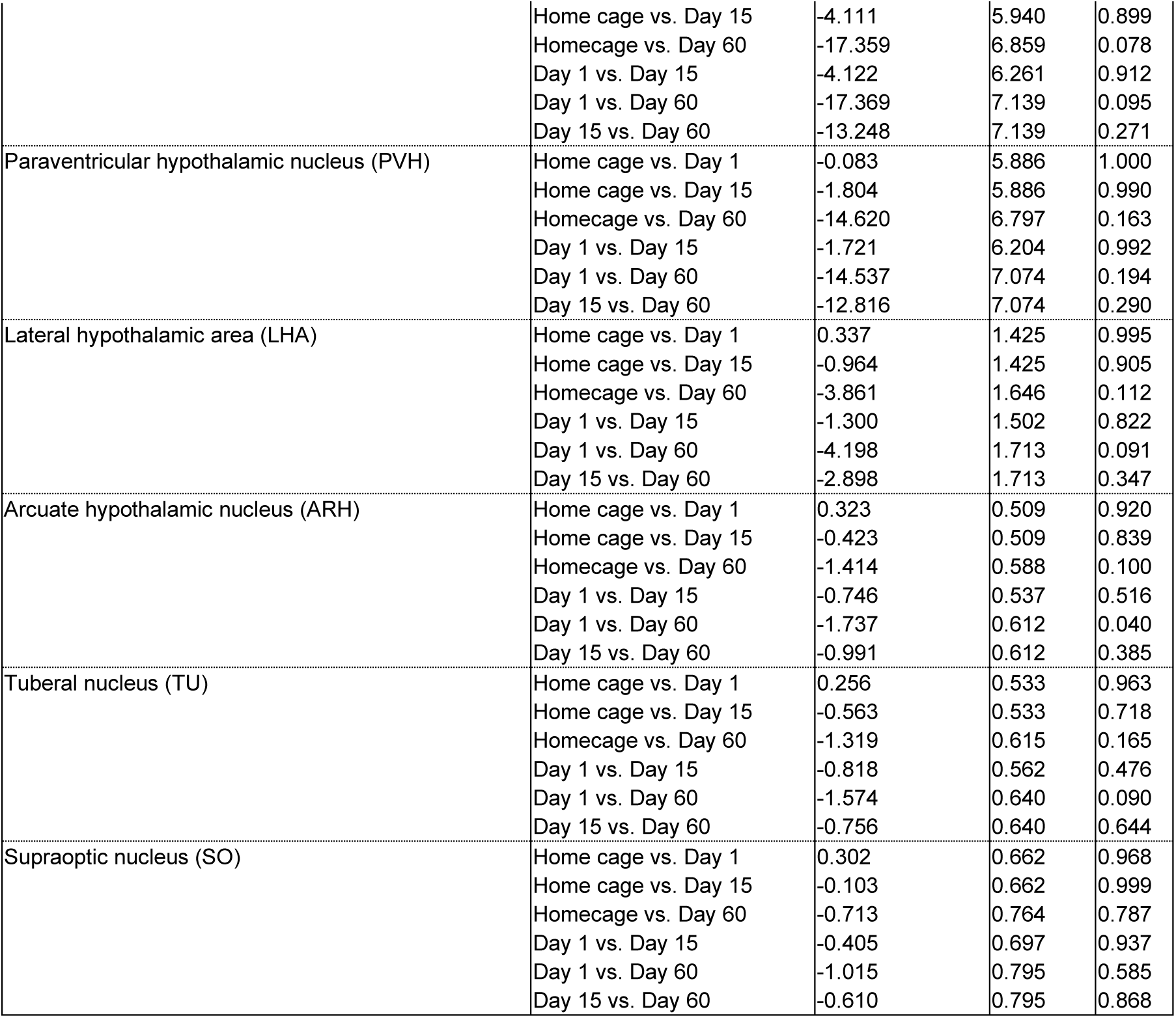
Statistical output for Figure 4D: Analysis of Z-scored counts from 64 subdivisions within AP -1.08 to AP -1.28 coronal subvolume (n=30)

## Notes

**Competing Interest Statement:** The authors declare that they do not have any conflicts of interest (financial or otherwise) related to the text of the paper. The research was supported by the NIDA Intramural Research Program funds to the labs of Yavin Shaham and Bruce Hope. SAG received funding from NIH PRAT 1FI2GM117583-01, NIDA R00DA045662, NIDA R01DA054317, NIDA 1UG3DA053802, NIDA P30DA048736 and NARSAD Young Investigator Award 27082. RM received funding from the NIH Center for Compulsive Behaviors. ES received funding from the Washington Research Foundation Fellowship Program. ORD and DP were supported by the NIDA IRP Scientific Director’s Fellowship for Diversity in Research.

### Competing Interest Statement

The authors have declared no competing interest.

